# Photoswitchable epothilone-based microtubule stabilisers allow GFP-imaging-compatible, optical control over the microtubule cytoskeleton

**DOI:** 10.1101/2021.03.31.437838

**Authors:** Li Gao, Joyce C. M. Meiring, Constanze Heise, Ankit Rai, Adrian Müller-Deku, Anna Akhmanova, Julia Thorn-Seshold, Oliver Thorn-Seshold

## Abstract

Optical methods to modulate microtubule stability and dynamics are promising approaches to reach the micron- and millisecond-scale resolution needed to decrypt the diverse roles of the microtubule cytoskeleton in biology. However, such optical methods have until now focussed nearly exclusively on microtubule destabilisation. Here, we introduce “STEpos” as light-responsive epothilone reagents, designed to photoswitchably bind to tubulin and stabilise lateral contacts in the microtubule lattice. Using a novel styrylthiazole photoswitch, designed to allow the hydrogen-bonding that is key to epothilone potency, we have created the first set of GFP-orthogonal photoswitchable microtubule stabilisers. The STEpos can photocontrol microtubule polymerisation, cell division, and cellular microtubule dynamics with micron- and second-scale spatiotemporal precision. STEpos offer substantial improvements of potency, solubility, and ease-of-use compared to the only previous photopharmaceuticals for microtubule stabilisation. The intriguing structure-photoswitching-activity relationship insights from this work will also assist future developments of improved STEpo reagents, and we anticipate that these will contribute greatly to high-precision cytoskeleton research across the fields of biophysics, cargo transport, cell motility, cell division, development, and neuroscience.

## Introduction

All cellular processes depend on spatiotemporal regulation of protein function. Molecular tools to modulate protein function with the micrometre spatial precision and millisecond temporal precision inherent to these processes, are extremely valuable for precise biological research.^1,2^ Cytoskeleton biology in particular would benefit from such tools, since hundreds of cellular processes rely simultaneously on the cytoskeletal scaffolding networks, with tight spatial regulation and with temporally dynamic response to changing conditions. Therefore, spatiotemporally-specific tools to modulate e.g. the microtubule and actin cytoskeleton, offer unique opportunities to decrypt and modify many different classes of biological functions.^3,4^

Microtubules (MTs) are giant, hollow, tube-like noncovalent polymers of α/β-tubulin heterodimers.^5^ They are formed into a spatially structured network extending throughout the cell, that is rapidly remodelled through regulated cycles of growth and shrinkage in order to fulfil their spatiotemporally-regulated functions. These functions include acting as scaffolding for cell shape and for mechanical processes, as tracks for cargo transport by motor proteins, and as an organising system for proteins and structures within the cell. This makes MT network structure and remodelling dynamics vital in diverse fields, including cell migration, cell division and development (e.g. by supporting the segregation of chromosomes), and neuroscience (e.g. supporting the formation and maintenance of specialized extensions like axons and dendrites).^6,7^

Drugs that modulate MT stability and/or de/polymerisation dynamics are prime research tools for all these fields, being useful to nonspecifically suppress MT-dependent cellular processes. Archetypical MT stabilisers (polymerisers) include taxanes, epothilones, and laulimalide; while notable destabilisers (depolymerisers) include colchicine analogues, vinca alkaloids, auristatins/dolastatins, and maytansines (**Fig 1a**). While both MT destabilisers and MT stabilisers can be used to suppress MT polymerisation dynamics in cell culture, they enable different biological applications particularly *in vivo*. For example, MT stabilisation in damaged mature neurons seems to promote axonal regeneration by reducing the formation of retraction bulbs and modulating glial scar formation, which is relevant to spinal cord injury.^8,9^ Conversely, colchicinoid MT destabilisers currently seem to show promise in reducing Covid-19 mortality^10^ by suppressing inflammatory responses, via a complex mechanism that is initiated by reducing microtubule polymer mass. Both MT stabilisers and destabilisers have substantial societal impact, with several, such as the taxanes, epothilones, and vinca alkaloids, having reached blockbuster status as cancer therapeutics due to their ability to interfere with cell proliferation.^11,12^

**Figure 1:**
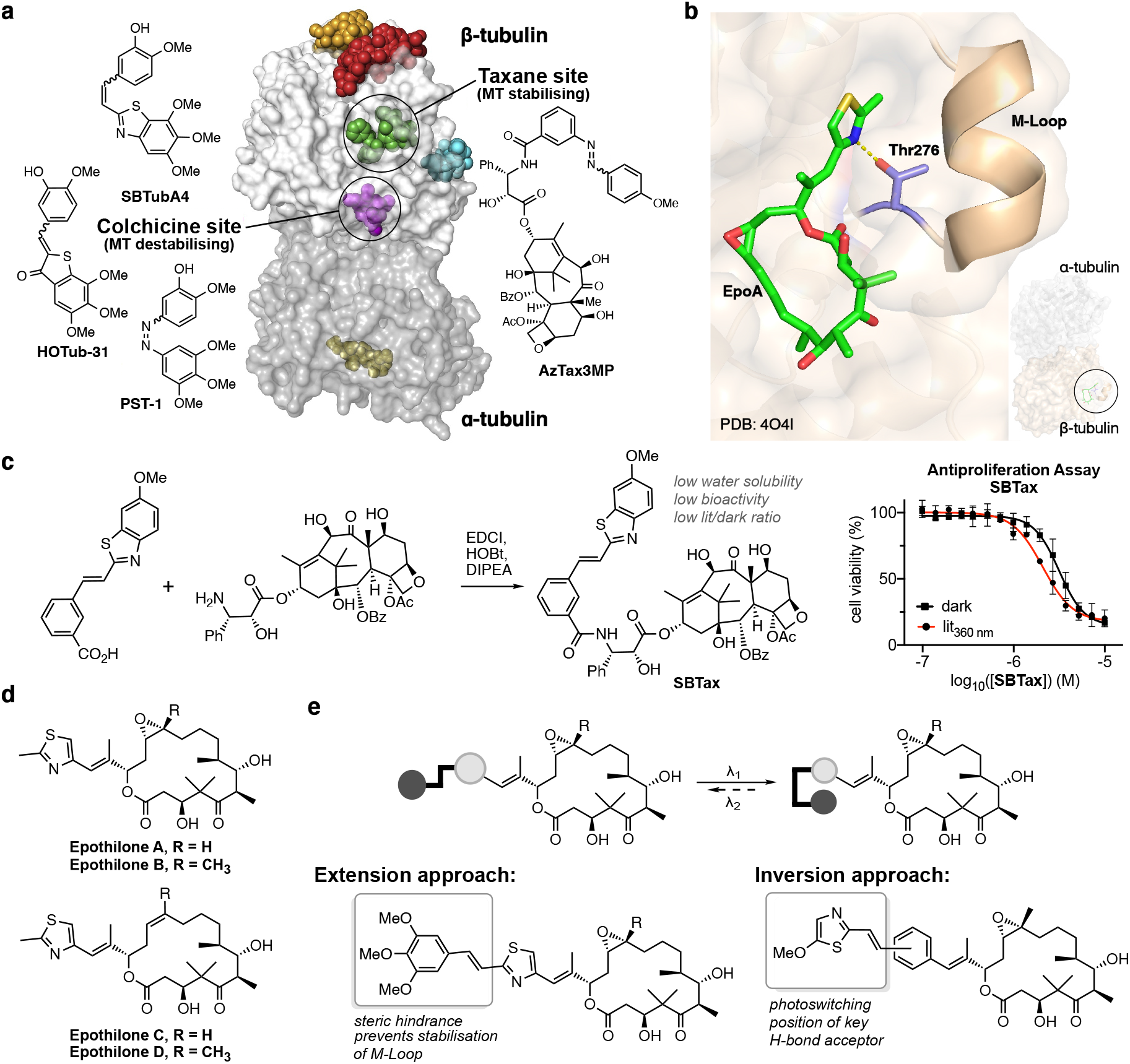
Microtubule stabiliser photopharmacology. **(a)** α/β-tubulin heterodimer with major drug binding sites coloured, and representative structures of the major photopharmaceutical chemotypes (figure from Steinmetz and Prota^21^). **(b)** Epothilone binding to β-tubulin stabilises the M-loop, which stabilises tubulin-tubulin lateral contacts and thus stabilises the MT. A key interaction is the hydrogen bond from Thr276 to the thiazole nitrogen.[adapted from PDB 4o4i]^22^ **(c)** Synthesis and biological evaluation of an SBT photoswitch scaffold based Taxol derivative **(d)** Principal epothilones. The epoxide of EpoA/B is a structural feature and is dispensible for binding (EpoC/D). **(e)** A peripheral photoswitch attachment strategy for epothilone, using the styrylthiazole (ST) photoswitch, was designed to photoswitchably impact the stabilisation of the M-loop and/or the position of an H-bond acceptor.

As research tools however, the lack of spatiotemporal precision with which these drugs can be applied has been a major hurdle for manipulation experiments that would ideally localise their activity on the scale of μm and ms. Indeed, spatiotemporally-specific approaches to modulate the stability and dynamics of the MT cytoskeleton are still in their infancy. Irreversibly photouncaged derivatives of MT inhibitors e.g. taxol and combretastatin A-4^13^ have been developed to improve the spatiotemporal precision of drug application in research, although these are not widely used. We are aware of only one optogenetic tool system for MTs, the photo-inactivatable π-EB1 system from Wittmann^14^, which can reversibly light-suppress MT polymerisation.

In contrast, reversibly photoswitchable analogues of taxane and colchicinoid MT-inhibiting drugs have been developed recently, that do offer cell-specific spatial precision and sub-second-scale temporal precision (**Fig 1a**). These photopharmaceuticals have practical advantages compared to photouncaging, such as fast light response, absence of phototoxic byproducts, easier handling and reduced potential for degradation. Additionally, although there is no directly comparable optogenetic system, the ability to apply these photopharmaceuticals across diverse model organisms, particularly in early developmental stages, is a strong advantage over genetic approaches. In general, optically controlled MT stabilisers are a particularly attractive target for use in cell culture and early development research, since the biological functions of the MT cytoskeleton essentially depend on stabilised or growing MTs. However, only one family of photoswitchable MT stabilisers has been reported; the azobenzene-based “AzTax” taxane analogues, which allowed fast, spatiotemporally precise, reversible control of MT dynamics, with a degree of subcellular control in neurons.^15^ In contrast, work on photoswitchable colchicinoid MT depolymerisers has explored a diversity of photoswitch scaffolds, from azobenzene (PST reagents),^3,16^, to hemithioindigo (HOTub/HITub),^17,18^, spiropyran-merocyanin^19^), and styrylbenzothiazole or “SBT” (SBTub reagents), each with their own drawbacks and advantages (**Fig 1a**).^20^

Notably, the SBT photoswitch offered a particularly attractive combination of advantages for biological research use: (1) The SBT scaffold is completely unaffected by standard GFP imaging at ca. 490 nm excitation, so biological assays can be imaged with common fluorescent proteins or markers without compromising the photopharmaceutical. This is a crucial factor for usability. The vast majority of biological models use GFP or closely-related fluorescent proteins, so GFP-orthogonal photoswitches are needed if researchers are to apply photopharmacology without having to change model (which is particularly problematic with transgenic animal lines that can take years to breed and validate). GFP-orthogonality is also needed to keep imaging channels free for the multiplexed readouts that are increasingly standard in cutting-edge research.^23^ (2) The SBT’s C=C chromophore is ideally situated for efficient photoswitching at 405 nm or 442 nm, so matching the typical photoactivation lasers installed in microscope - again a crucial feature for practical impact. (3) SBT is highly metabolically stable, and avoids biochemical photoswitch scission. This scission is generally problematic for both azobenzene isomers, but they also suffer from faster *Z-*isomer scission, that further complicates assay readouts^24,25^. More problematically still, in typical photopharmaceutical uses, photoswitches are built onto the molecular periphery (as they were in the AzTax reagents), so if either isomer suffers scission, it releases a non-photoswitchable fragment that is typically more potent than the photopharmaceutical and further compromises assay readout (discussed further elsewhere^20^). Taken together, these advantages of the SBTs proved crucial in allowing SBTubs to succeed as MT photoswitches in tissue and animal applications across a variety of animal models, while photopharmaceuticals based on the other photoswitches have not generally succeeded in moving beyond 2D cell culture or early embryo applications.^23^

We were therefore motivated to bring the powerful advantages of the SBT photoswitch to bear on the attractive class of microtubule stabilisers, to generate novel optically-controlled MT-stabilisers with improved practical applicability and scope. We now report the development of such GFP-orthogonal photopharmaceutical MT stabilisers, based on epothilone (**Fig 1b**).

## Results

### Initial approach: SBT-taxanes

We had initially applied the metabolically resistant, GFP-orthogonal SBT photoswitch to the taxane scaffold. We used the same design as we had previously used for the metabolically labile, non-GFP-orthogonal photoswitch azobenzene,^15^ deprotecting the sidechain 3’-amine of docetaxel and coupling it to a SBT-carboxylic acid to give SBT-taxane **SBTax** (**Fig 1c**). However, **SBTax** had poor solubility, unsatisfactory cellular antiproliferative potency, and its bioactivity was not significantly photoisomer-dependent (**Fig 1c**). This may simply reflect the taxanes’ weaknesses for photopharmaceutical adaptation, including their ‘ball and sidechain’ structure (with no obvious basis for rational introduction of isomer-dependent potency), low solubility, already high molecular weight, and structural and chemical complexity that limit reasonable synthetic modifications.

### Final approach: design of ST-epothilones (STEpos)

We therefore decided to switch away from the taxane scaffold, and to explore epothilone-based photopharmaceuticals instead. Epothilones are a structurally simpler class of microtubule stabilisers, that share the same binding pocket as taxanes (**Fig 1b**) but feature higher binding affinity and potency.^26,27^ Epothilones have proven particularly useful as research reagents, in that their greater solubility, bioavailability, and ability to cross the blood-brain barrier, allow applications inaccessible to taxanes: such as systemic (rather than local) administration for stabilising axons, to regenerate the central nervous system after injury.^28^ The synthetic tractability of the epothilones has enabled extensive drug analogue campaigns (**Fig 1d**), clarifying structure-activity relationship (SAR) features;^29,30^ and a range of epothilone derivatives have undergone advanced clinical trials, with ixabepilone being FDA-approved for treatment of metastatic breast cancer. A further advantage of the chemically simpler epothilone is that late-stage modifications that preserve binding activity while altering chemically or biologically problematic groups are known; in our case, this would later prove to be removal of the C12/C13 epoxide of epothilone B, to give the *Z-*alkene epothilone D. The SAR of epothilones suggests that while aryl-aryl substituents are not tolerated around C12/C13,^31^ even relatively large heterocycles can be tolerated in place of the thiazole, with potent thiazole, purine, quinoline and benzothiazole derivatives all being known, and larger rings particularly tolerated in epothilone D derivatives.^29^

X-ray studies also reveal the critical role of the epothilones’ nitrogen-containing aryl ring in forming and orienting the β-tubulin M-loop, so that this loop stabilises tubulin-tubulin lateral contacts, which in turn stabilise microtubules (**Fig 1b**). As well as sterics, the hydrogen bond from Thr276 to the ring nitrogen is a key interaction for this loop formation,^22^ which is underlined by e.g. the 20- to 100-fold potency loss on going from 2-pyridine to near-isosteric 3- and 4-pyridine epothilone analogues.^32^

Using an SBT-like photoswitch to replace the thiazole of epothilones therefore seemed attractive as a rational design principle (**Fig 1e**). (i) The size of the photoswitch would seem to be tolerated, yet sterics around the attachment site play a significant role in bioactivity, giving hope that E/Z isomerisation would influence tubulin-binding potency. (ii) The epothilone scaffold seemed sufficiently water-soluble to support the photoswitch, without incurring the limitations seen with the taxanes. (iii) Finally, the photoswitch’s ring nitrogen raises the exciting opportunity, that photoisomer-dependently repositioning the thiazole portion could modulate its H-bond-accepting capacity, and therefore drastically influencing its M-loop-stabilising activity, and biological potency.

To minimise the overall size and insolubility of the photoswitch, we chose to work with our recently developed styrylthiazole (ST) photoswitches instead of SBTs. STs are isosteric to azobenzenes, but feature similar electronics and absorption spectra as the GFP-orthogonal, metabolically-resistant SBTs (see also below).^20^ We explored two design strategies. Firstly, an “extension approach” aimed to project the photoswitch into the M-loop as the *trans* isomer, alleviating steric pressure as the *cis* isomer, and therefore designing towards a *cis-*active photopharmaceutical (NB: the *cis* isomer of SBTs always places the nitrogen atom accessibly towards the outside,^20^ therefore we did not anticipate that the *cis* isomer would be incapable of acting as an H-bond acceptor). As elsewhere in our work, we chose a trimethoxyphenyl group for this approach, as we find that the combination of the out-of-plane orientation of the central methoxy group that reduces π-stacking, and the additional hydrophilicity of the three extra OMe groups, contribute greatly to water solubility. Secondly, an “inversion approach” tested whether photoswitching could bring an H-bond accepting ring nitrogen sufficiently close to Thr276 to favour *cis*-specific binding (**Fig 1e**). We decided to compare *meta* with *para* photoswitch connection in this inverted approach. We anticipated that the *meta-*connection could bring the thiazole nitrogen close enough to be effective, and although the *para-*connected switch might sterically clash too greatly with the M-loop in both isomers, it could serve as a useful control for e.g. phototoxicity and solubility. As far as we are aware, there is no systematic study of the dependency of ST photochemistry on substituents; but we reasoned that, analogously to azobenzenes, raising the electron density in the thiazole by including an OMe substituent should help shift ST absorption bands towards the visible. It should also be noted that by retaining the vinyl substituent, both extension and inversion designs benefit from an additional shift towards the visible.

### Synthesis of STEpos

Horner-Wadsworth-Emmons olefination of the epothilone methyl ketone **1** is a flexible, established route to attach diverse aryl rings to epothilones,^33^ and we determined to use it to install pre-formed styrylthiazoles (STs). Starting from epothilone B, we cleaved the double bond by ozonolysis^34^ then TES-protected the free OH groups to give protected methyl ketone **2** (**Fig 2a**). The ST photoswitch phosphonates **8** and **12a/b** were synthesised by Arbuzov reaction from the chlorides, which in turn had been accessed from heterocyclic ring closure of the cinnamic acid derivatives (**6** and **10a/b**, **Fig 2b-c**). Initially, methyl ketone **2** was olefinated with phosphonate **12a**, giving inversion photoswitch **STEpo1** after deprotection (**Fig 2d**). Although Mulzer has shown that the epoxide tolerates various basic, oxidative, reductive and electrophilic reagents,^35^ the loss of scarce material by epoxide opening under dilute acid conditions prompted us to deoxygenate the epoxide instead,^36^ which still preserves bioactivity in the epothilone series.^37^ We therefore targeted ST derivatives of epothilone D, and deoxygenated **2** with WCl_6_^38^ to give TES-protected methyl ketone **3** (**Fig 2a**) in good yields. Olefination with ST phosphonates, then deprotection, yielded extension design **STEpo2** and inversion designs **STEpo3** and **STEpo4** in milligram quantities (**Fig 2e**) (see Supporting Information for details).

**Figure 2:**
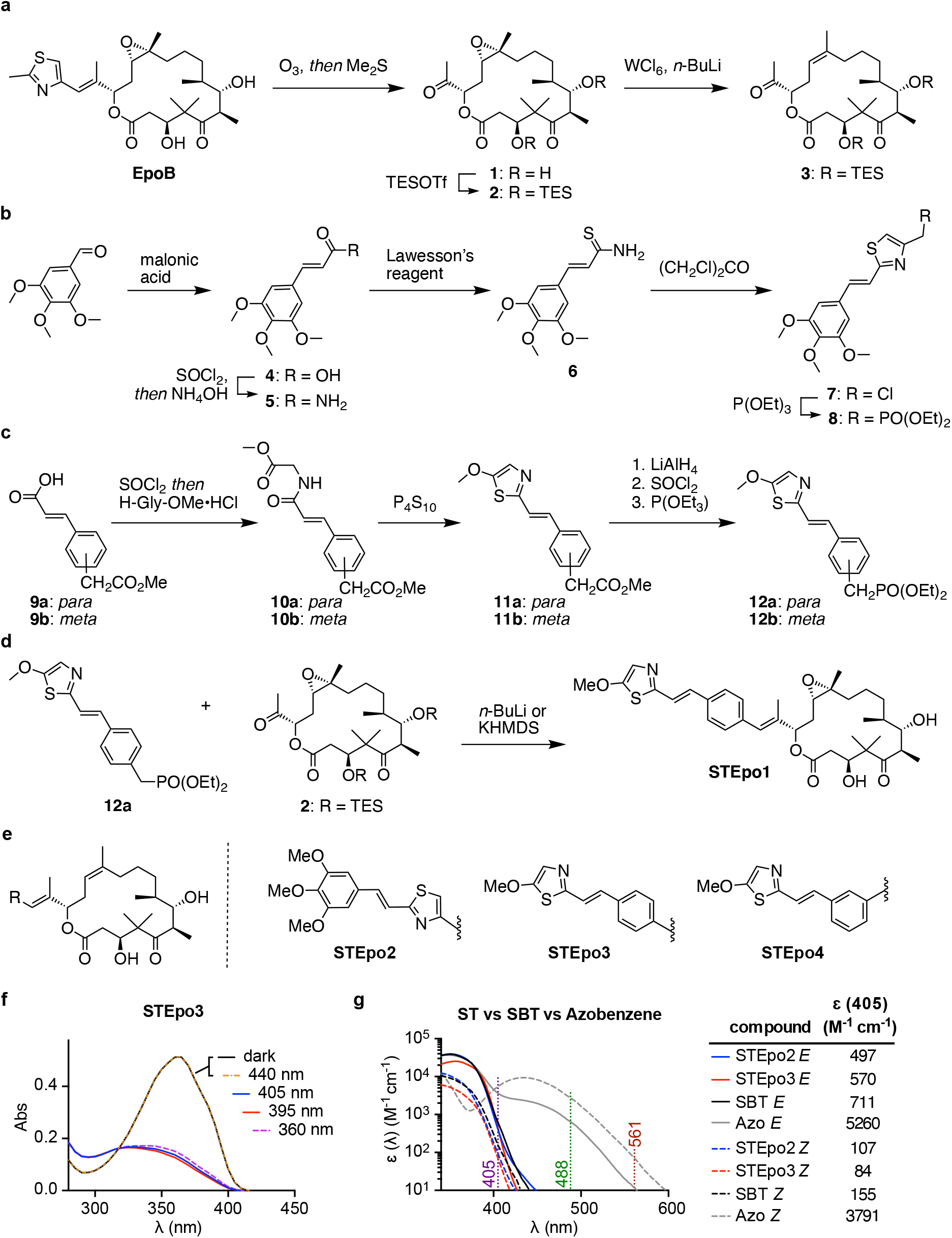
Synthesis and photocharacterisation of STEpo photopharmaceuticals. **(a)** route to epothilone methyl ketones **2** and **3**. **(b)** route to ST phosphonate **8**. **(c)** route to ST phosphonates **12a** and **12b**. **(d)** Epothilone B inversion design derivative, **STEpo1**. **(e)** Epothilone D derivatives **STEpo2** (extension) and **STEpo3**/**STEpo4** (inversion). **(f)** Photostationary state (PSS) equilibrium UV-Vis spectra of **STEpo3** during sequential irradiations starting from the dark state (all-*E*) shows no response to 440 nm light, efficient isomerisation to mostly-*Z* by wavelengths from 405—360 nm, and no degradation under this illumination. **(g)** Comparison of absorption spectra of *E* and *Z* isomers of **STEpos** with an SBT and a typical *para-*methoxylated azobenzene. At 405 nm, the **STEpos** have approx two thirds of the absorption coefficient of an SBT photoswitch. The log scale plot highlights the full orthogonality of the ST (and SBT) photoswitches to 488 nm (GFP) or 561 nm (RFP) imaging, whereas the azobenzene is still responsive to high-intensity focused imaging lasers at these wavelengths. See Supporting Information for details.

### Photoisomerisation and GFP-orthogonality of STEpos

The STEpos’ performance regarding photoisomerisation and photostability was characterised by UV-Vis spectroscopy. Similarly to SBTs,^20^ the STEpos could be reproducibly photoisomerised to majority-*Z* photostationary states (PSSs) by near-UV light (360-410 nm; ca. 85% *Z*; see Supporting Information) (**Fig 2f, Fig S1a-b**). We observed gradual degradation with illumination below 340 nm, presumably by [2+2] cycloaddition, though not by illumination above this wavelength. The vinylthiazole photoswitch **STEpo2** was blueshifted by ca. 10 nm compared to the methoxythiazole (inversion) photoswitches. Surprisingly, the STs were blueshifted by only ca. 10 nm compared to the benzannulated SBTs (**Fig S1d**). The *Z*-STs were thermally stable, giving <1% relaxation to *E-*STs in solution at 37°C after 8 h (**Fig S1c**).

Typically, the major *E*-ST absorption band was centred around 355 nm; the 20%-of-maximum intensity was at ca. 392 nm, with a sharp cutoff above this wavelength (ε dropping by a factor of 10 every 12 nm). The major *Z*-ST absorption band was centred around 325 nm; the 20%-of-maximum intensity was at 382 nm, also with a sharp cutoff above this wavelength (ε dropping by a factor of 10 every 16 nm) (**Fig 2g**). These sharp cutoffs are crucial features. These are responsible for the ST’s absorption being effectively zero above 440 nm, which should ensure that it is entirely unaffected by GFP imaging either with lasers (488 nm) or with broader filtered excitation sources (typically 490±25 nm). This is in sharp contrast to azobenzene photoswitches, particularly those that benefit from polymethoxylation, which are significantly impacted by imaging of GFP, YFP, and even RFP (**Fig 2g**).^20^

### STEpos give photoisomer-dependent cellular bioactivity

We first evaluated the GFP-orthogonal, photoswitchable STEpos for their photoisomer-dependent antiproliferative activity in cells in culture, which MT inhibitors cause by blocking mitosis. We incubated HeLa cervical cancer cells with *E-*STEpos in the dark (all-*E*) or lit by non-phototoxic pulsed illuminations from low-power 360 nm LEDs (<1 mW/cm^2^; <1% duty cycle; *in situ* photoswitching to mostly-*Z*) as described previously,^3,20^ and assessed cell viability 44 h later.

All compounds showed antiproliferative activity in the submicromolar range, and all compounds had reproducibly light-dependent bioactivity with up to ca. 8-fold potency shift upon isomerisation (**Fig 3a**). The epothilone B *para-*inversion analogue **STEpo1** (IC_50_(lit) ca. 3 nM, IC_50_(dark) ca. 12 nM) was ca. 100-fold more potent than its epothilone D inversion counterpart **STEpo3**, and ca. 35-fold more than epothilone D extension analogue **STEpo2**, which were likewise lit-active with similar 4-fold photoswitchability of bioactivity. Interestingly, epothilone D *meta-*inversion analogue **STEpo4** was instead dark-active, with *Z*-**STEpo4** being equipotent to *E*-**STEpo3**, and *E*-**STEpo4** being equipotent to *Z*-**STEpo3**. This reversal suggests an intriguing structural basis for photoisomer-dependent activity in the STEpo series, which is an ongoing challenge in our research. It also highlights the absence of phototoxicity from either the ST or the illumination protocols, since this would otherwise result in consistent lit-toxic effects.

**Figure 3:**
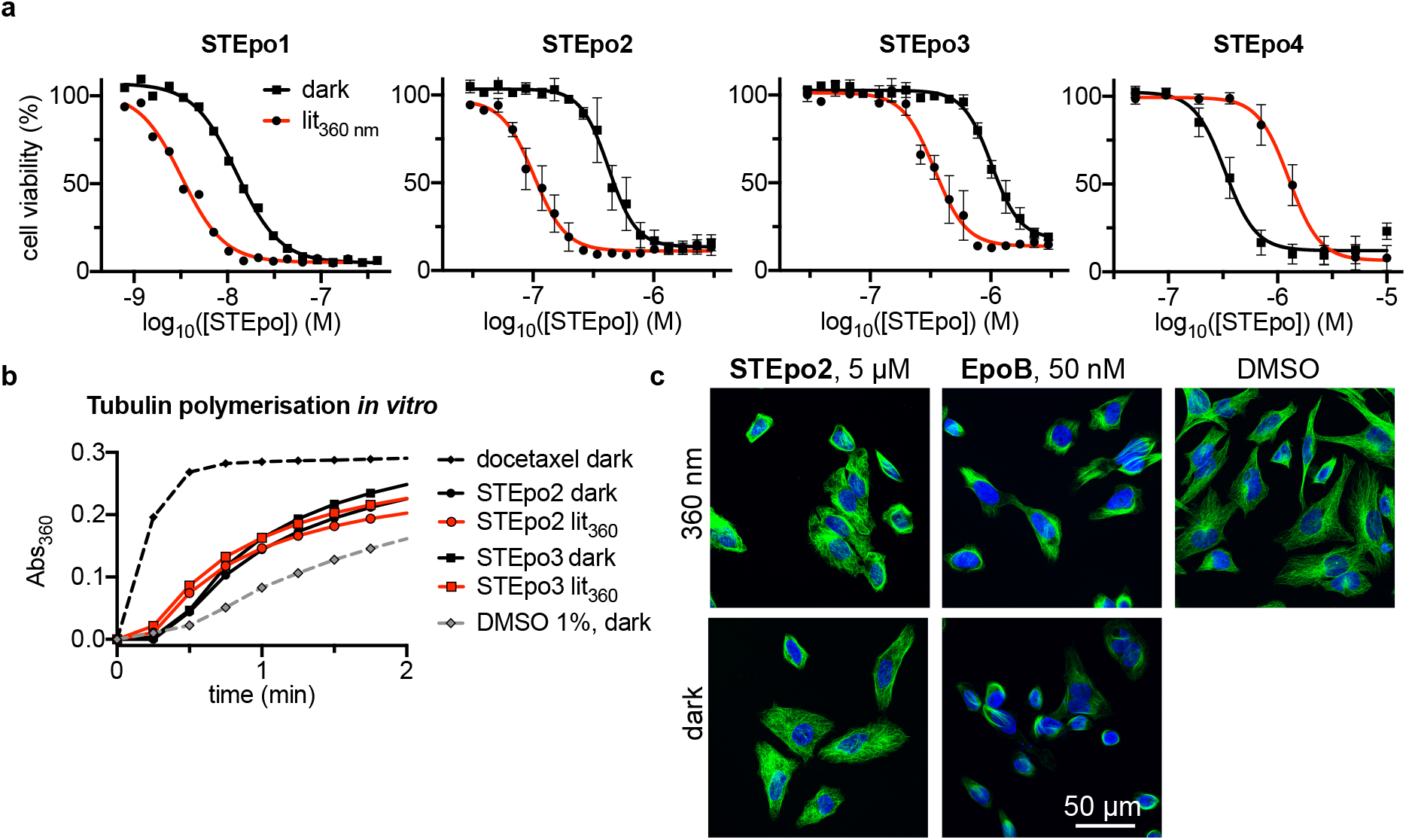
Cellular bioactivity and mechanism of action of STEpos. **(a) STEpos** have potent and light-dependent antiproliferative activity (HeLa cells, 44 h incubation; all-*E* dark conditions versus mostly-*Z* UV-lit conditions; **STEpo1**: one representative of three independent experiments shown; **STEpo2-4**: three replicates, mean+SD). **(b) STEpo2/3** promote tubulin polymerisation (turbidimetric *in vitro* assay; greater absorbance corresponds to a greater degree of polymerisation; time to onset of polymerisation should be examined) with a small light-dependency visible (earlier onset). **(c)** Immunofluorescence imaging of cells treated with **STEpo2** shows breakdown of MT architecture and mitotic arrests under lit conditions (mostly-*Z*), but nearly no disorganization in the dark (all-*E*) (HeLa cells, 20 h incubation; α-tubulin in green, DNA stained with DAPI in blue).

We did not continue with **STEpo1** due to its lower synthetic availability, and since **STEpo2** offered an identical photoswitchability of bioactivity (ratio of apparent cellular activity, comparing limiting illumination conditions) while not facing solubility problems. Since lit-active photopharmaceuticals present the greatest practical advantage for biological use, we also did not continue with **STEpo4**.

### Mechanism of action of STEpo

To begin testing their molecular mechanism of light-dependent bioactivity in a simplified cell-free system, we assayed the photoswitchably cytotoxic, lit-active **STEpo2/3** for light-dependent enhancement of polymerisation of purified tubulin (polymerisation in bulk). Both isomers were polymerisation enhancers, indicating that they act directly on tubulin as do the epothilones; and the time to polymerisation onset was noticeably shorter with pre-lit mostly-*Z* stocks, than with all-*E* stocks: matching their cellular activity pattern (**Fig 3b**).

Next, we imaged the network architecture of the MT cytoskeleton in cells incubated with **STEpo2/3** to examine their cellular mechanism of isomer-dependent bioactivity. Dark (all-*E*) assays showed markedly less impact on the MT network architecture than *in situ* lit assays (mostly *Z*-isomer) (**Fig 3c**). The mitotic arrests and MT depolymerisation that are particularly visible on the lit assays are hallmarks of MT inhibitors, matching the assumption that their photoisomer-dependent cytotoxicity arises from their *Z-*isomers more potently inhibiting MT dynamics and stability in cells.

Finally, we studied the capacity of **STEpo2** to permit *in situ* photocontrol over microtubule polymerisation in cell-free settings: with spatial resolution on the μm scale allowing us to optically target *individual microtubules*, and with temporal resolution on the scale of seconds.

We used TIRF microscopy to image a reconstituted microtubule polymerisation system. This uses the non-hydrolysable GTP analogue GMPCPP to form stable microtubule “seeds” that are fixed to a glass surface for spatially-localised imaging; free tubulin is then applied together with the drug under testing and as well as MT plus end-tracking protein EB3, added to improve visualization of dynamic MT tips (**Fig 4a**)^39,40^; finally, GTP is supplied and the seeds initiate normal cycles of growth and shrinkage using the free tubulin in the mixture. Using the labelled seed of each MT as a static reference point, kymographs tracking the growing and shrinking tips of the MT reveal the frequency of shrinkage events as the “spikes” in the kymograph. Typically, MTs depolymerise back to the seed before restarting polymerisation. MT-stabilising drugs decrease the frequency of shrinkage events and can stabilize a shrinking MT before the seed is reached (known as a rescue). In the event that the drug stabilises a different protofilament number of MT than the seeds have (14 for GMPCPP), stable rescue sites will be observed, while if the drug stabilizes the protofilament number matching the seed, rescues will occur stochastically at different sites along the microtubule,^39^ so offering mechanistic insights into the nature of MT stabilisation. As epothilones are 14-protofilament MT stabilisers, we expected that if our **STEpos** retained the same MT-stabilising properties with *Z-*specific potency, they would permit these MT rescues after illumination (**Fig 4b**): which we aimed to apply with spatial specificity to regions near single selected MT tips.

**Figure 4:**
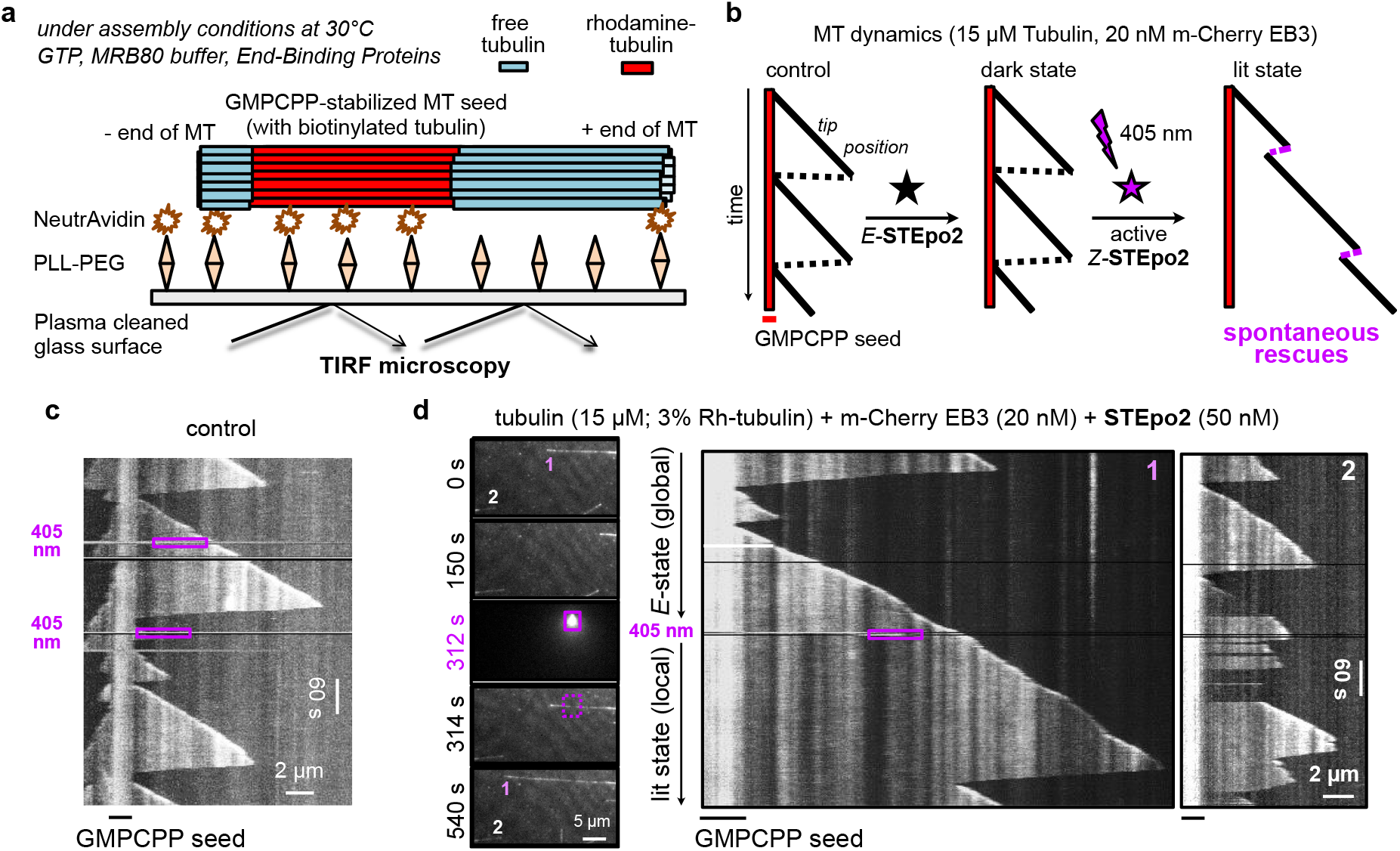
Spatiotemporal control over MT dynamics in cell-free systems by photocontrolling STEpo2: **(a)** Schematic of the *in vitro* MT dynamics imaging assay, and **(b)** cartoon of kymographs illustrating MT growth and catastrophe patterns during normal conditions, then expected patterns for **SBEpo2** as a 405 nm-dependent 14-protofilament MT stabiliser. **(c)** No-drug control kymograph showing MT shrinkage frequency and polymerisation restarts at the seed (405 nm illuminations in the purple dotted boxes). **(d)** Time-lapse images at left, showing two microtubules in the field of view (labelled 1 and 2), with their kymographs at right. 405 nm illumination at 312 s is in the purple box region of time-lapse panel/kymographs (0.01% of the total field of view).

During control assays, kymographs revealed that shrinkage events proceeded all the way down to the seed before polymerisation restarted, as expected, and this MT behaviour was unaffected by 405 nm illumination (**Fig 4c, Fig S2a**). When *E-***SBEpo2** was added at 50 nM concentration, MT shrinkage events were similar to control conditions, reaching the GMPCPP seeds before restart (first 5 min, **Fig 4d**), although higher concentrations predictably caused MT stabilisation (**Fig S2b**). Local 405 nm illumination near a single selected microtubule (labelled “1” in **Fig 4d**) stabilised it, increasing MT length and showing rescue. By contrast, other MTs which were further from the illumination zone (e.g. “2”) maintained similar shrinkage frequency as before, though with some spontaneous rescues (next 5 min, **Fig 4d**; **Movie S1**; see too **Fig S2c**). These stringent assays confirm that **STEpo2** is an epothilone-like, *Z-*isomer-specific stabiliser of 14-protofilament MTs, which can be spatiotemporally targeted in cell-free settings with micron-and-second scale precision.

### Live cell photocontrol with STEpo

To test the STEpos’ ability to enable *in situ* photocontrol of MT dynamics with spatiotemporal precision, we imaged **STEpo2**-treated cells transfected with the fluorescently-labelled MT end binding protein EB3-GFP. EB3 marks the GTP cap region of MTs, thus revealing the dynamics of polymerising MT plus ends as dynamic “comets”;^41^ and EB3 imaging is an effective spatiotemporally-resolved readout for isomerisation-dependent control of MT dynamics by photoswitchable inhibitors.^20^

Cells were therefore imaged continually with excitation at 488 nm. Imaging was begun before **STEpo2** application to establish baseline MT dynamics. Low intensity 405 nm pulsing was applied, inducing negligible comet count reduction. Then **STEpo2** was applied at 0.6 μM, again resulting in no noticeable change of EB3 statistics. That 488 nm imaging of **STEpo2**-treated cells did not induce inhibition of MT dynamics (unlike the performance of azobenzene-based photopharmaceuticals)^20^ matches their design for GFP-compatibility by absorption cut-off. However, in a fourth phase, 405 nm photoactivation pulses were additionally applied, and now MT dynamics were rapidly suppressed (EB3 comet count halved within seconds; P < 0.0001) (**Fig 5**; **Movie S2**). These assays show that *in situ* photoswitching of **STEpo2** is an effective approach to precise, noninvasive MT control in cellular settings.

**Figure 5:**
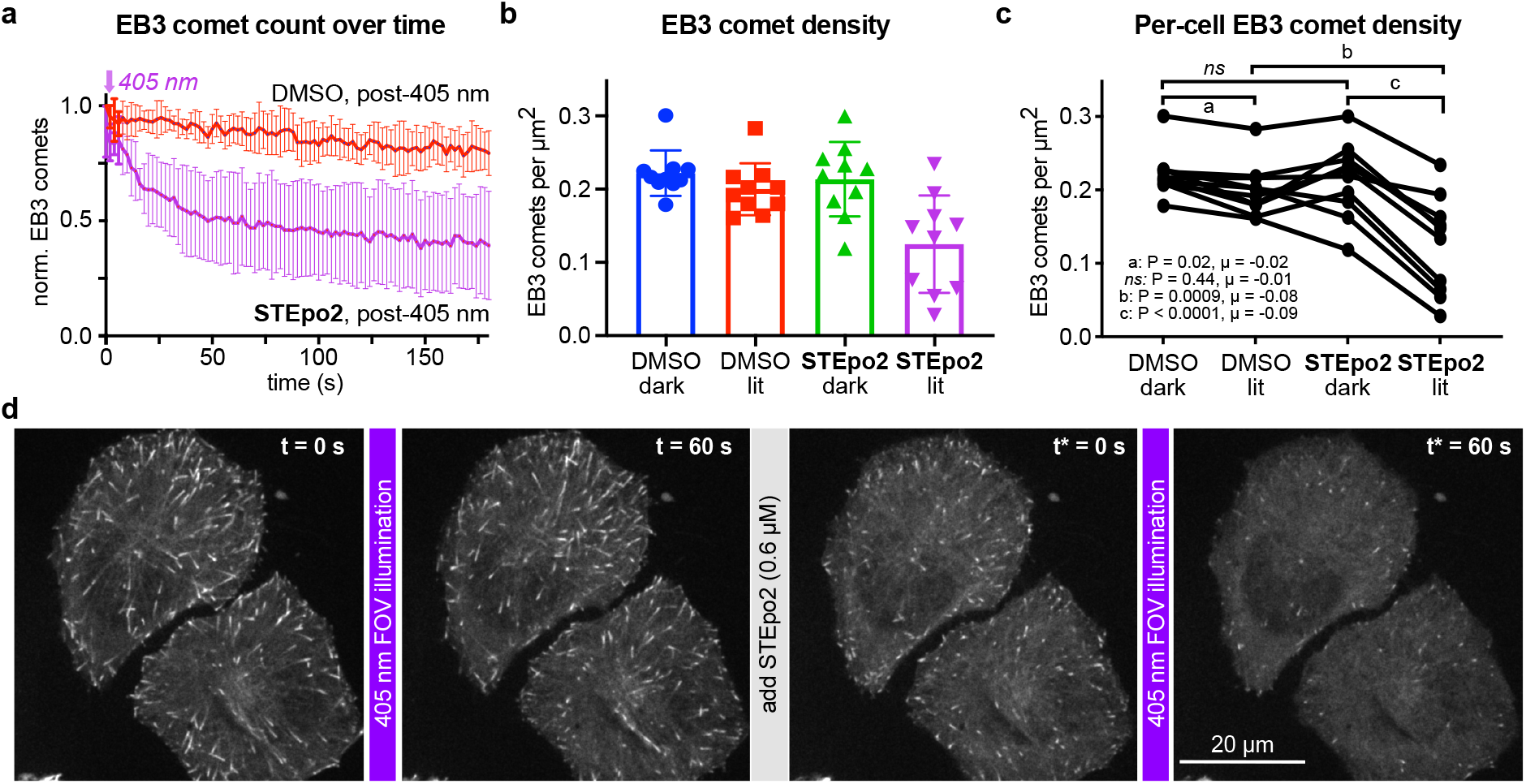
Spatiotemporal control over cellular MT dynamics with STEpo2: Live cell EB3-GFP “comets” during GFP imaging with 488 nm (HeLa cells). MT inhibition with **STEpo2** is initiated upon 405 nm illumination (cells first treated with 1% DMSO, then imaged for 60 s for baseline, then photoactivated from time t = 0 with 405 nm during imaging over 3 min; then an additional 0.6 μM *E*-**STEpo2** was applied, and 405 nm/imaging applied from time t* = 0. 10 cells acquired). **(a)** mean ± SD comet count (each cell normalised to t=0 of the DMSO control). **(b-c)** Comet density statistics show large and significant differences upon **STEpo2** photoactivation (dark at t = 0 or t* = 0, lit at t = 60 s or t* = 60 s; **b** shows pooled data, **c** shows longitudinal traces per cell; P-values and μ mean differences as annotated). **(d)** Stills from representative movie (data related to **Movie S2**).

## Conclusions

Methods to modulate biological processes with the spatiotemporal resolution appropriate to their endogenous functioning, have become increasingly powerful and effective over recent years. Photopharmacology in particular is rapidly evolving into an elegant technique for high-precision, non-invasive biological control in fast-response and/or spatially-localised applications from neuroscience^42,43^ to cytoskeleton research^3,4^. As the optical instrumentation of biological research becomes ever more sophisticated and widespread, the value of robust optically-controlled tools will continue to rise: particularly for studies of inherently (sub)cellularly- and subsecond-resolved processes, such as those that the microtubule cytoskeleton supports.^44^

In this work we report the first photopharmaceuticals based on the epothilone MT stabiliser scaffold. The mid-nanomolar potencies, reliable solubility, and the robust photoswitchability of bioactivity of all the **STEpos** in this work are very promising features for future research, and can be favourably compared to the micromolar potencies, solubility problems, and variable photoswitchability of bioactivity we have found here and previously^15^ for photoswitch-derivatised taxanes. We feel that designing towards isomer-dependent interference with a partially disordered structural element (here, the tubulin M-loop) represents an exciting general principle for photopharmacology, which currently focusses on relatively rigid steric clashes that may be unrealistic to identify and photocontrol. By exploring uncommonly large epothilone derivatives, we have also revealed aspects of epothilone SAR that can lead to identifying which steric and polarity aspects determine **STEpo** binding and/or M-loop orientation, and so will inspire the rational design of still-more-potent systems with still greater photoswitchability of bioactivity.

This is also, we believe, the first use of the compact styrylthiazole (ST) as a photoswitch scaffold for biology: one that is isosteric to azobenzene, yet offers significant practical benefits. Here we highlight its GFP-orthogonality, that enables easy imaging, and its intriguing and biologically relevant potential to act as an H-bond acceptor, which matched the requirements of the epothilone system. We feel that GFP-orthogonality is currently an under-appreciated goal amongst chemical reagent makers. The race toward NIR-capable photoswitches for deep tissue photoisomerisation is understandable, even though deep tissue isomerisation has to contend with significant spatial scattering of illumination (as is seen by shining a red laser pointer through a finger) and so may sacrifice the spatial precision that is one of photopharmacology’s greatest advantages. A recent design shift, driven by biologists, is however underway: emphasising biochemically stable, imaging-orthogonal photoswitch scaffolds that can be easily employed in tissue slice and embryo/early animal research across a variety of models.^20,23^ This shift has great potential to serve the biological research community with a valuable palette of rapid-response, non-phototoxic, byproduct-free reagents for studies in diverse fields. To continue this progress will rely on expanding our scope of photopharmaceutical design principles and of photoswitch scaffolds. In this context, unlocking a useful new photoswitch scaffold, may prove as valuable as unlocking a new drug or binding site for a key, central biological player.

There are several challenges for the ongoing development of **STEpos**. Topics in focus in our group include: (i) To understand and to harness the greater potency seen for the epoxide-bearing derivative with milder chemical routes. (ii) To improve the speed of photoactivation at the 405 nm laser line by slightly redshifting the ST absorption bands (just a 15 nm shift would produce a 10-fold rate enhancement due to the steepness of absorption cutoff). We have elsewhere achieved this with increasing substituent donor strength (e.g. *para*-NMe_2_Ph-instead of Ph- for the extension design)^23^; for the inverted design, further photochemical exploration of the ST scaffold will be needed, although again we anticipate that increasing the electron donating capacity of the thiazole will be key. (iii) To achieve bidirectional photoswitching with a photoswitch offering the GFP-orthogonality of an ST (and the metabolic robustness of an SBT). This requires increasing the separation between the absorption bands of the *E* and *Z* isomers, which is an object of ongoing study.

In conclusion, the **STEpos** are reliably soluble, easy to apply, GFP-orthogonal, potent MT-stabilising photoswitches for cell culture, and should also easily find applications through to *in vivo* GFP/multiplexed imaging experiments in near-surface settings. These **STEpos**, or future reagents, will undoubtedly bring great value to cytoskeleton research across fields from development and motility to transport and cell division. In particular we expect that the **STEpo** toolset will benefit research into axonal repair following injury, which can be favoured by MT stabilisers, though by still-unknown mechanisms whose elucidation has so far been stymied by poor spatiotemporal precision of MT stabilisation.^8,9,15,28^ Lastly, we are optimistic that by contributing to innovations in the realms of photoswitch scaffold chemistry and of rational photopharmaceutical design, the **STEpos** are a promising advance not only for high-precision microtubule biology, but also towards the refinement of high-performance photopharmacology against other protein targets.

## Acknowledgements

This research was supported by funds from the German Research Foundation (DFG: Emmy Noether grant number 400324123; SFB 1032 project B09 number 201269156; SFB TRR 152 project P24 number 239283807; and SPP 1926 project number 426018126 to O.T.-S.). J.C.M.M. acknowledges support from an EMBO Long Term Fellowship. We thank Jan Huebner and Bayer for the gift of EpoB, Dirk Trauner for enabling the material transfers, Monique Preusse for early cell viability testing, and Rebekkah Hammar for performing the tubulin polymerisation assay. We are grateful to Henrietta Lacks, now deceased, and to her surviving family members for their contributions to biomedical research.

## Author contributions

L.G. designed and performed the epothilone syntheses, performed photocharacterisation and cell viability assays, and coordinated data assembly. J.C.M.M. performed live cell EB3 imaging during photoswitching and coordinated data assembly. C.H. performed cell viability assays, immunofluorescence staining, and cell cycle analysis. A.M.-D. synthesised the SBT-taxane. A.A. supervised EB3 imaging. J.T.-S. performed cell cycle analysis, coordinated data assembly and supervised all other cell biology. O.T.-S. designed the concept and experiments, supervised all other experiments, coordinated data assembly and wrote the manuscript.

