## Supplemental Information for "Photoswitchable epothilone-based microtubule stabilisers allow GFP-imaging-compatible, optical control over the microtubule cytoskeleton"

### Part A: Chemical Synthesis

#### Conventions

Abbreviations: The following abbreviations are used: Hex – distilled isohexanes, EA – ethyl acetate, DCM – dichloromethane, Et – ethyl, Ac – acetyl, Me – methyl, MeCN – acetonitrile, DMSO – dimethylsulfoxide, PBS – phosphate buffered saline, FA – Formic Acid, TFA – trifluoro acetic acid, TEA – triethyl amine, LR – Lawesson's reagent.

Safety Hazards: no unexpected or unusually high safety hazards were encountered.

Reagents and Conditions: Unless stated otherwise, (1) all reactions and characterizations were performed with unpurified, undried, non-degassed solvents and reagents, used as obtained, under closed air atmosphere without special precautions; (2) "hexane" used for chromatography was distilled from commercial crude isohexane fraction by rotary evaporation; (3) "column" and "chromatography" refer to manual flash column chromatography on Merck silica gel Si-60 (40−63 μm); (4) MPLC flash column chromatography refers to purification on a Biotage Isolera Spektra, using prepacked silica cartridges from Biotage; (5) procedures and yields are unoptimized; (6) yields refer to isolated chromatographically and spectroscopically pure materials, corrected for residual solvent content; (7) all eluent and solvent mixtures are given as volume ratios unless otherwise specified, thus "1:1 Hex:EA" indicates a 1:1 (v/v) mixture of hexanes and ethyl acetate; (8) chromatography eluents e.g. "0→25% EA:Hex" indicate a linear gradient of eluent composition.

Thin-layer chromatography (TLC) was run on 0.25 mm Merck silica gel plates (60, F-254), typically with Hex:EA eluents, except where indicated. UV light (254 nm) was used as a visualizing agent, with cross-checking by 365 nm UV lamp. Compounds not containing chromophores were stained with cerium ammonium molybdate instead. TLC characterizations are abbreviated as $\mathbf{R}_{\boldsymbol{f}}$ = 0.64 (EA:Hex = 1:1).

NMR: Standard NMR characterization was by $^{1}$H- and $^{13}$C-NMR spectra on a Bruker Ascend 400 (400 **MHz** & 100 **MHz** for $^{1}$H and $^{13}$C respectively) or a Bruker Ascend 500 (500 **MHz** & 125 **MHz** for $^{1}$H and $^{13}$C, respectively). Known compounds were checked against literature data and their spectral analysis is not detailed unless necessary. Chemical shifts (δ) are reported in ppm calibrated to residual non-perdeuterated solvent as an internal reference[1]. Peak descriptions singlet (s), doublet (d), triplet (t), quartet (q) and multiplet (m).

Analytical HPLC and Mass Spectra: Analytical HPLC-MS measurements were performed on an Agilent 1100 SL coupled HPLC-MS system with (a) a binary pump to deliver $H_2O$:MeCN eluent mixtures containing 0.1% formic acid at a 0.4 mL/min flow rate, (b) Thermo Scientific Hypersil GOLD™ C18 column (1.9 μm; 3 × 50 mm) maintained at 25°C, whereby the solvent front eluted at $t_{ret}$ = 0.5 min, (c) an Agilent 1100 series diode array detector used to acquire peak spectra of separated compounds/isomers in the range 200-550 nm after manually

baselining across each elution peak of interest to correct for eluent composition effects, (d) a Bruker Daltonics HCT-Ultra mass spectrometer used in ESI mode at unit mass resolution. Run conditions were a linear gradient of $H_2O$:MeCN eluent composition from the starting ratio through to 10:90, applied during the separation phase (first 5 min), then 0:100 maintained until all peaks of interest had been observed (typically 2 min more); the column was equilibrated with the $H_2O$:MeCN eluent mixture for 2 minutes before each run. HRMS was carried out by the Zentrale Analytik of the LMU Munich using ESI or EI ionization as specified. LRMS was carried out on an expression CMS by Advion with either APCI or ESI as ionization source.
Preperative HPLC (prep-HPLC): Prep-HPLC purification was carried out on a 1260 Infinity II Preparative LC System by Agilent using an Agilent reversed phase Prep-HT C18 column (21.2 x 250 mm, 10 µm) at a 20 mL/min flow rate.

#### Synthesis of STEpos

##### TES protected STEpo1:

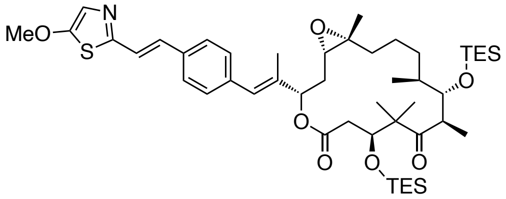

To a solution of styrylthiazole phosphonate **12a** (50 mg, 0.14 mmol, 2.5 eq) in dry THF (1.5 mL) was added sodium bis(trimethylsilyl)amide (2 M in THF, 50 µL, 0.10 mmol, 1.8 eq) and the solution was stirred at –78°C for 30 min. A solution of methyl ketone **2** (35 mg, 55 µmol, 1 eq) in THF (1 mL) was added dropwise, and the red reaction mixture was stirred for 2 h at –78°C. Then the reaction mixture was quenched with sat. aq. $NH_4Cl$ (2 mL), and allowed to warm to room temperature. The two phases were separated, and the aqueous layer was extracted with EA (3 × 5mL). The combined organic layers were dried over $Na_2SO_4$ and concentrated under reduced pressure. The obtained residue was purified by normal phase MPLC (0→20% EA:Hex, elutes at 17% EA) to afford TES protected **STEpo1** (8.3 mg, 9.7 µmol, 18%) as colorless oil.
**$^1$H-NMR (500 MHz, CDCl$_3$):** δ = 7.45 (d, *J* = 8.2 Hz, 2H), 7.29 (d, *J* = 8.0 Hz, 2H), 7.15 – 7.05 (m, 3H), 6.55 (s, 1H), 5.29 (dd, *J* = 8.9, 2.8 Hz, 1H), 4.14 (dd, *J* = 7.2, 4.8 Hz, 1H), 3.95 (s, 3H), 3.91 (d, *J* = 9.1 Hz, 1H), 3.09 – 3.00 (m, 1H), 2.84 (dd, *J* = 9.9, 3.6 Hz, 1H), 2.73 – 2.60 (m, 2H), 2.22 (dt, *J* = 14.9, 3.2 Hz, 1H), 1.93 (d, *J* = 1.4 Hz, 3H), 1.92 – 1.84 (m, 1H), 1.74 (dt, *J* = 12.5, 6.0 Hz, 1H), 1.44 – 1.31 (m, 4H), 1.28 (s, 3H), 1.25 (s, 1H), 1.21 (s, 1H), 1.18 (s, 3H), 1.14 (s, 3H), 1.10 (d, *J* = 6.8 Hz, 3H), 1.00 – 0.97 (m, 12H), 0.93 (t, *J* = 8.0 Hz, 9H), 0.64 (dd, *J* = 16.5, 8.1 Hz, 12H) ppm. **$^{13}$C-NMR (125 MHz, CDCl$_3$):** δ = 215.4, 170.8, 162.3, 155.1, 137.1,

136.8, 134.5, 131.7, 129.6, 127.0, 126.6, 122.6, 121.7, 79.8, 77.1, 75.7, 62.5, 62.1, 61.4, 53.5, 48.1, 39.9, 36.7, 33.9, 32.2, 31.3, 29.7, 24.9, 23.8, 22.4, 19.6, 17.5, 14.1, 7.2, 7.0, 5.5, 5.3 ppm. **$R_f$** = 0.48 (EA:Hex = 2:8). **HRMS (ESI, positive):** calc. for $C_{47}H_{76}NO_7SSi_2^+$ [M+H]$^+$ 854.4876, found 854.4864.

**STEpo1:**

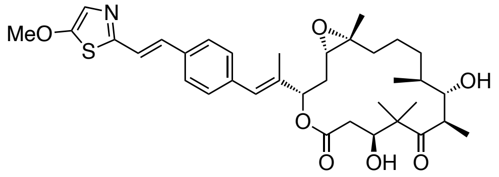

TES protected **STEpo1** (8.3 mg, 9.7 µmol, 1 eq) was dissolved in dry THF (1 mL) and cooled to 0°C. HF•pyr (70%, 40 µL, excess) was added and the reaction mixture was allowed to warm to room temperature. After 5 h, the reaction mixture was carefully quenched with sat. aq. $NaHCO_3$ (10 mL). The two layers were separated, and the aqueous layer was extracted with EA (3 × 10 mL). The combined organic layers were dried over $Na_2SO_4$ and concentrated under reduced pressure. Purification by prep-HPLC (40→100% MeCN:$H_2O$) afforded **STEpo1** (1 mg, 1.6 µmol, 16%) as colorless solid.

**$^1$H-NMR (500 MHz, $CDCl_3$):** δ = 7.46 (d, *J* = 8.3 Hz, 2H), 7.28 (d, *J* = 8.4 Hz, 2H), 7.13 – 7.05 (m, 3H), 6.58 (s, 1H), 5.48 (dd, *J* = 7.0, 4.0 Hz, 1H), 4.08 (dd, *J* = 9.3, 3.1 Hz, 1H), 3.95 (s, 3H), 3.81 (t, *J* = 4.4 Hz, 1H), 3.35 – 3.26 (m, 1H), 2.83 (dd, *J* = 7.2, 5.5 Hz, 1H), 2.57 (dd, *J* = 14.7, 9.6 Hz, 1H), 2.48 (d, *J* = 3.5 Hz, 1H), 2.35 (s, 1H), 2.13 – 2.03 (m, 2H), 2.02 – 1.92 (m, 5H), 1.70 (dd, *J* = 15.1, 7.9 Hz, 3H), 1.57 – 1.43 (m, 3H), 1.35 (s, 3H), 1.30 (s, 3H), 1.18 (d, *J* = 7.0 Hz, 3H), 1.11 (s, 3H), 1.01 (d, *J* = 7.0 Hz, 3H) ppm. **$R_f$** = 0.24 (EA:Hex = 6:4). **HRMS (ESI, positive)**: calc. for $C_{35}H_{48}NO_7S^+$ [M+H]$^+$ 626.3146, found 626.3144.

**TES protected STEpo2:**

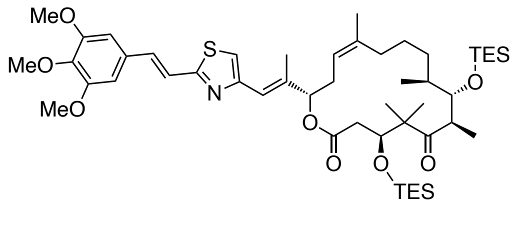

To a solution of styrylthiazole phosphonate **8** (45 mg, 0.11 mmol, 3.5 eq) in dry THF (1.5 mL) was added *n*-BuLi (2.5 M in hexane, 30 µL, 76 µmol, 2.5 eq) and the purple solution was stirred at –78°C for 45 min. A solution of methyl ketone **3** (19 mg, 30 µmol, 1 eq) in THF (1 mL) was added dropwise, and the reaction mixture was allowed to slowly warm to 0°C. After 2 h the orange reaction mixture was quenched with sat. aq. $NH_4Cl$ (2 mL), and transferred to a separation funnel. The two layers were separated, and the aqueous layer was extracted with

EA (3 × 5mL). The combined organic layers were dried over $Na_2SO_4$ and concentrated under reduced pressure. The obtained residue was purified by normal phase MPLC (0→20% EA:Hex, elutes at 10% EA) to afford TES protected **STEpo2** (12 mg, 13 µmol, 44%) as colorless oil.

**$^1$H-NMR (500 MHz, $C_6D_6$):** δ = 7.52 (d, *J* = 16.1 Hz, 1H), 7.35 (d, *J* = 16.1 Hz, 1H), 6.78 (s, 1H), 6.62 (s, 1H), 6.53 (s, 2H), 5.33 (d, *J* = 9.8 Hz, 1H), 5.27 – 5.21 (m, 1H), 4.23 (d, *J* = 8.9 Hz, 1H), 4.15 (d, *J* = 9.4 Hz, 1H), 3.82 (s, 3H), 3.32 (s, 6H), 3.06 – 2.96 (m, 1H), 2.87 – 2.75 (m, 2H), 2.64 (ddd, *J* = 20.2, 11.8, 4.9 Hz, 2H), 2.38 (s, 3H), 2.03 (dd, *J* = 14.3, 6.3 Hz, 1H), 1.93 – 1.78 (m, 4H), 1.75 (s, 3H), 1.39 – 1.34 (m, 2H), 1.21 (d, *J* = 6.9 Hz, 3H), 1.19 (s, 3H), 1.12 – 1.07 (m, 21H), 0.86 – 0.80 (m, 6H), 0.77 (t, *J* = 7.9 Hz, 6H), 0.73 (s, 3H) ppm. **$^{13}$C-NMR (125 MHz, $C_6D_6$):** δ = 214.0, 170.9, 166.0, 154.9, 154.4, 140.6, 140.4, 139.4, 135.1, 131.4, 121.2, 120.3, 120.1, 116.7, 105.1, 80.6, 76.7, 60.6, 55.8, 53.4, 48.0, 39.4, 38.1, 32.7, 32.5, 32.0, 30.2, 28.1, 25.2, 23.4, 23.3, 19.5, 17.7, 15.3, 7.5, 7.4, 6.1, 5.9 ppm. **$R_f$** = 0.56 (EA:Hex = 2:8). **LRMS (APCI, positive):** calc. for $C_{49}H_{80}NO_8SSi_2$ $[M+H]^+$ 898.5138, found 899.5.

**STEpo2:**

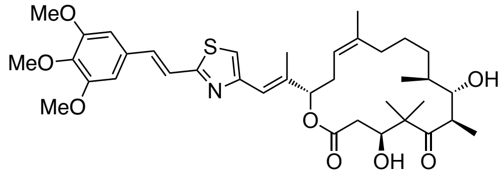

TES protected **STEpo2** (12 mg, 13 µmol, 1 eq) was dissolved in dry THF (1 mL) and cooled to 0°C. HF•pyr (70%, 75 µL, excess) was added and reaction mixture was allowed to warm to room temperature. After 1.5 h, only mono-deprotected TES **STEpo2** was observed. Another 50 µL HF•pyr was added at 0°C and stirring was continued another 2 h at room temperature. The reaction mixture was carefully quenched with sat. aq. $NaHCO_3$ (10 mL). The two layers were separated, and the aqueous layer was extracted with EA (3 × 10 mL). The combined organic layers were dried over $Na_2SO_4$ and concentrated under reduced pressure. Purification by prep-HPLC (40→100% MeCN:$H_2O$ + 0.1% FA) afforded **STEpo2** (6.5 mg, 9.7 µmol, 73%) as slightly yellowish solid.

**$^1$H-NMR (500 MHz, $C_6D_6$):** δ = 7.53 (d, *J* = 16.1 Hz, 1H), 7.30 (d, *J* = 16.1 Hz, 1H), 6.85 (t, *J* = 1.4 Hz, 1H), 6.63 (s, 1H), 6.56 (s, 2H), 5.48 – 5.43 (m, 1H), 5.20 (dd, *J* = 10.2, 5.2 Hz, 1H), 4.22 (dd, *J* = 11.0, 2.8 Hz, 1H), 3.83 (s, 3H), 3.74 (dd, *J* = 4.1, 2.8 Hz, 1H), 3.34 (s, 6H), 2.97 (qd, *J* = 6.8, 2.8 Hz, 1H), 2.69 (dt, *J* = 15.0, 9.8 Hz, 1H), 2.39 (dd, *J* = 14.9, 11.0 Hz, 1H), 2.26 (d, *J* = 1.3 Hz, 4H), 2.18 – 2.10 (m, 2H), 1.81 (dt, *J* = 19.0, 5.7 Hz, 2H), 1.64 (t, *J* = 1.5 Hz, 4H), 1.28 – 1.20 (m, 3H), 1.10 (d, *J* = 6.8 Hz, 3H), 1.03 (d, *J* = 7.0 Hz, 3H), 1.01 (s, 3H), 0.97 (s, 3H) ppm. **$^{13}$C-NMR (125 MHz, $C_6D_6$):** δ = 219.6, 170.1, 166.1, 154.6, 154.4, 140.7, 139.6, 138.6, 135.2, 131.4, 121.4, 120.8, 119.7, 116.3, 105.2, 79.4, 74.5, 72.8, 60.6, 55.8, 53.6, 42.1,

40.1, 38.7, 32.8, 32.0, 31.7, 25.9, 23.2, 22.8, 18.4, 16.1, 15.9, 13.9 ppm. **R*f*** = 0.31 (EA:Hex = 4:6). **HRMS (ESI, positive):** calc. for $C_{37}H_{52}NO_8S^+$ $[M+H]^+$ 670.3408, found 670.3410.

**TES protected STEpo3:**

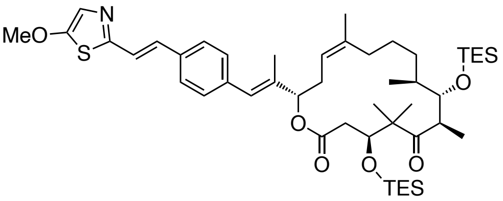

To a solution of styrylthiazole phosphonate **12a** (39 mg, 0.11 mmol, 3 eq) in dry THF (1.5 mL) was added sodium bis(trimethylsilyl)amide (2 M in THF, 44 µL, 88 µmol, 2.5 eq) and the red solution was stirred at –78°C for 50 min. A solution of methyl ketone **3** (22 mg, 35 µmol, 1 eq) in THF (1 mL) was added dropwise, and the reaction mixture was allowed to slowly warm to 0°C. After 2.5 h the reaction mixture was quenched with sat. aq. $NH_4Cl$ (5 mL), and the reaction mixture was allowed to warm to room temperature. The two layers were separated, and the aqueous layer was extracted with EA (3 × 5mL). The combined organic layers were dried over $Na_2SO_4$ and concentrated under reduced pressure. The obtained residue was purified by normal phase MPLC (5→25% EA:Hex, elutes at 11%) to afford TES protected **STEpo3** (12 mg, 14 µmol, 41%) as colorless oil.
**$^1$H-NMR (400 MHz, CDCl$_3$):** δ = 7.47 (d, *J* = 8.2 Hz, 2H), 7.29 (d, *J* = 8.1 Hz, 2H), 7.17 (d, *J* = 6.7 Hz, 2H), 7.08 (s, 1H), 6.51 (s, 1H), 5.19 (t, *J* = 8.2 Hz, 1H), 5.12 (d, *J* = 9.9 Hz, 1H), 4.12 (dd, *J* = 8.0, 3.9 Hz, 1H), 3.97 (s, 3H), 3.94 (d, *J* = 9.0 Hz, 1H), 3.11 – 2.98 (m, 1H), 2.73 – 2.68 (m, 2H), 2.44 (d, *J* = 8.5 Hz, 1H), 2.08 (dd, *J* = 14.9, 5.7 Hz, 1H), 1.94 (s, 3H), 1.77 – 1.63 (m, 5H), 1.60 – 1.49 (m, 2H), 1.25 (s, 3H), 1.18 (s, 3H), 1.11 (d, *J* = 7.3 Hz, 6H), 0.99 (t, *J* = 7.8 Hz, 12H), 0.90 (t, *J* = 7.9 Hz, 9H), 0.67 (q, *J* = 8.0 Hz, 6H), 0.59 (q, *J* = 7.5 Hz, 6H) ppm.
**$^{13}$C-NMR (125 MHz, CDCl$_3$):** δ = 215.6, 171.1, 162.2, 155.8, 140.6, 138.1, 134.0, 129.7, 127.0, 126.1, 119.4, 80.1, 80.0, 76.1, 61.7, 53.7, 48.0, 39.7, 37.5, 32.6, 32.1, 32.1, 31.3, 29.9, 27.5, 24.4, 24.0, 23.2, 22.8, 19.3, 17.7, 14.6, 14.3, 7.3, 7.1, 5.7, 5.4 ppm. **R*f*** = 0.28 (EA:Hex = 15:85). **HRMS (ESI, positive)**: calc. for $C_{47}H_{76}NO_6SSi_2$ $[M+H]^+$ 838.4926, 838.4926 found.

**STEpo3:**

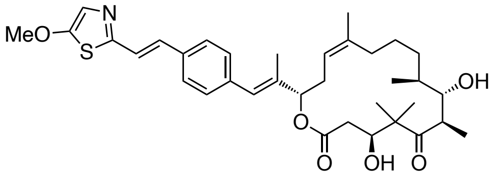

TES protected **STEpo3** (12 mg, 14 µmol, 1 eq) was dissolved in dry THF (2 mL) and cooled to 0°C. HF•pyr (70%, 100 µL, excess) was added and reaction mixture was allowed to warm to room temperature. After 5 h the reaction mixture was carefully quenched with sat. aq.

$NaHCO_3$ (10 mL). The two layers were separated, and the aqueous layer was extracted with EA (3 × 10 mL). The combined organic layers were dried over $Na_2SO_4$ and concentrated under reduced pressure. Purification by prep-HPLC (40→100% MeCN:$H_2O$ + 0.1% FA) afforded desired product *E*-**STEpo3** (1.7 mg, 28 µmol, 20%) as colorless solid and *Z*-**STEpo3** (1.3 mg, 21 µmol, 25%) as colorless solid.

**$^1$H-NMR (500 MHz, $C_6D_6$):** δ = 7.27 (d, *J* = 16.1 Hz, 1H), 7.21 – 7.18 (m, 4H), 7.13 (d, *J* = 12.3 Hz, 1H), 7.03 (s, 1H), 6.72 (s, 1H), 5.45 (dd, *J* = 9.9, 2.2 Hz, 1H), 5.20 (dd, *J* = 10.1, 5.2 Hz, 1H), 4.07 (ddd, *J* = 10.4, 6.7, 2.9 Hz, 1H), 3.73 (s, 1H), 3.17 (s, 3H), 2.93 (qd, *J* = 6.8, 3.0 Hz, 1H), 2.80 (s, 1H), 2.68 (dt, *J* = 15.0, 9.9 Hz, 1H), 2.34 – 2.27 (m, 2H), 2.14 (dd, *J* = 15.3, 2.9 Hz, 1H), 2.11 – 2.07 (m, 1H), 2.00 (d, *J* = 7.0 Hz, 1H), 1.86 (s, 3H), 1.84 – 1.78 (m, 2H), 1.64 (s, 3H), 1.62 – 1.56 (m, 1H), 1.23 – 1.18 (m, 3H), 1.08 (d, *J* = 6.8 Hz, 3H), 1.02 (d, *J* = 7.1 Hz, 3H), 0.92 (s, 3H), 0.89 (s, 3H) ppm. **$^{13}$C-NMR (125 MHz, $C_6D_6$):** δ = 219.3, 170.0, 162.7, 154.8, 138.6, 137.7, 137.5, 135.0, 131.7, 129.9, 127.1, 126.8, 123.4, 122.3, 121.4, 79.7, 74.5, 73.1, 60.5, 53.2, 42.3, 40.0, 38.7, 32.7, 31.9, 30.2, 26.1, 23.2, 22.5, 18.8, 16.2, 14.9, 14.0 ppm. **$R_f$** = 0.43 (EA:Hex = 6:4). **HRMS (ESI, positive)**: calc. for $C_{35}H_{48}NO_6S^+$ $[M+H]^+$ 610.3197, found 610.3201.

**STEpo4:**

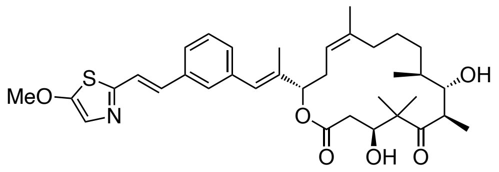

To a solution of styrylthiazole phosphonate **12b** (30 mg, 82 µmol, 4.6 eq) in dry THF (1 mL) was added *n*-BuLi (2 M in THF, 25 µL, 63 µmol, 3.5 eq) and the solution was stirred at –78°C for 1 h. A solution of methyl ketone **3** (11 mg, 18 µmol, 1 eq) in THF (1 mL) was added dropwise, and the reaction mixture was allowed to slowly warm to 0°C. Reaction progress was checked by TLC after 2.5 h, showing a weak fluorescent spot **$R_f$** = 0.32 (EA:Hex = 1:9). The reaction mixture was quenched with sat. aq. $NH_4Cl$ (4 mL), and the reaction mixture was allowed to warm to room temperature. The two layers were separated, and the aqueous layer was extracted with EA (3 × 5mL). The combined organic layers were dried over $Na_2SO_4$ and concentrated under reduced pressure. The crude product was filtered over a short silica plug (EA:Hex =3:7) to remove phosphonate **12b** and the filtrate was concentrated. The crude was redissolved in dry THF (1 mL) and cooled to 0°C. HF•pyr (70%, 100 µL, excess) was added and reaction mixture was allowed to warm to room temperature. After 3 h the reaction mixture was carefully quenched with sat. aq. $NaHCO_3$ (10 mL). The two layers were separated, and the aqueous layer was extracted with EA (3 × 10 mL). The combined organic layers were dried over $Na_2SO_4$ and concentrated under reduced pressure. Purification over prep-HPLC

(40→100% MeCN:$H_2O$ + 0.1% FA) afforded **STEpo4** (0.8 mg, 1.3 µmol, 7%) as colorless solid.

**$^1$H-NMR (400 MHz, $C_6D_6$):** δ = 7.37 (s, 1H), 7.27 (s, 1H), 7.23 (s, 1H), 7.21 (s, 1H), 7.08 – 7.04 (m, 2H), 7.01 (s, 1H), 6.73 (s, 1H), 5.46 (dd, *J* = 9.7, 2.1 Hz, 1H), 5.21 (dd, *J* = 9.8, 4.8 Hz, 1H), 4.09 (dd, *J* = 10.8, 2.4 Hz, 1H), 3.73 (t, *J* = 3.3 Hz, 1H), 3.46 (br, 1H), 3.16 (s, 3H), 2.93 (dd, *J* = 6.8, 3.0 Hz, 1H), 2.81 (br, 1H), 2.69 (dt, *J* = 14.8, 9.8 Hz, 1H), 2.32 (dd, *J* = 15.1, 10.9 Hz, 2H), 2.18 – 2.09 (m, 2H), 1.87 – 1.77 (m, 5H), 1.65 (d, *J* = 1.5 Hz, 3H), 1.60 (d, *J* = 4.0 Hz, 1H), 1.23 (s, 3H), 1.07 (d, *J* = 6.8 Hz, 3H), 1.02 (d, *J* = 7.0 Hz, 3H), 0.92 (s, 3H), 0.89 (s, 3H) ppm. **$R_f$** = 0.51 (EA:Hex = 6:4). **HRMS (ESI, positive):** calc. for $C_{35}H_{48}NO_6S^+$ $[M+H]^+$ 610.3197, found 610.3203.

#### Synthesis of epothilone methyl ketones

Synthesis of the Epothilone B/ Epothilone D methyl ketones was performed closely following previously reported protocols by Nicolaou et al[2,3] and characterization matched the literature.

##### Synthesis of methyl ketone 1

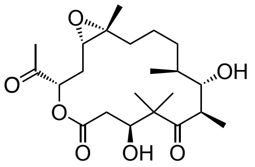

Freshly generated ozone was bubbled through a stirred solution of Epothilone B (117 mg, 230 µmol, 1.0 eq) in dichloromethane (5 mL) at −78°C. The reaction mixture was immediately quenched with dimethyl sulfide (170 µL, 2.31 mmol, 10 eq) when the color turned blue. The reaction mixture was warmed to room temperature and stirred for 1 h before the solvent was removed under reduced pressure and purified by MPLC (20→60% EA:Hex) to give methyl ketone 1 (76 mg, 184 µmol, 80%) as colorless solid and recovered starting material Epothilone B (11 mg, 22 µmol, 9%).

**$^1$H-NMR (500 MHz, $CDCl_3$):** δ = 5.30 (dd, *J* = 10.5, 1.9 Hz, 1H), 4.30 (dd, *J* = 10.6, 3.2 Hz, 1H), 3.69 (t, *J* = 4.3 Hz, 1H), 3.24 (qd, *J* = 6.8, 4.7 Hz, 1H), 2.82 (dd, *J* = 9.4, 3.1 Hz, 1H), 2.55 (d, *J* = 10.6 Hz, 1H), 2.52 (d, *J* = 10.6 Hz, 1H), 2.34 (dt, *J* = 15.3, 2.6 Hz, 1H), 2.28 (s, 3H), 2.25 (d, *J* = 3.0 Hz, 1H), 1.79 – 1.70 (m, 2H), 1.68 – 1.61 (m, 1H), 1.51 – 1.43 (m, 1H), 1.41 (s, 3H), 1.37 – 1.29 (m, 2H), 1.28 (s, 3H), 1.27 – 1.22 (m, 2H), 1.19 (d, *J* = 6.8 Hz, 3H), 1.08 (s, 3H), 0.98 (d, *J* = 6.9 Hz, 3H) ppm. **$^{13}$C-NMR (125 MHz, $CDCl_3$):** δ = 220.6, 204.9, 170.7, 76.8, 74.5, 71.6, 62.4, 62.1, 53.4, 42.7, 39.9, 37.4, 32.8, 31.2, 28.9, 26.3, 23.2, 22.5, 22.4, 18.0, 17.2, 14.4 ppm. **$R_f$** = 0.28 (EA:Hex = 1:1). **LRMS (ESI, positive):** calc for $C_{22}H_{37}O_7^+$ $[M+H]^+$ 413.2534, 413.3 found.

**Synthesis of TES protected methyl ketone 2.**

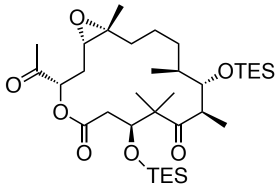

To a stirred solution of methyl ketone **1** (44 mg, 107 µmol, 1 eq) in dry dichloromethane (2 mL) was added 2,6-lutidine (37 µL, 320 µmol, 3 eq) and triethylsilyl trifluoromethanesulfonate (58 µL, 265 µmol, 2.4 eq) at –78°C. The reaction mixture was stirred for 30 min, before quenching with water (5 mL) and warming up to room temperature. The reaction mixture was transferred to a separation funnel and the organic layer was separated. The aqueous layer was extracted with EA (3 × 5 mL) and the combined organic layers were dried over $Na_2SO_4$. The solvent was removed under reduced pressure and purification by normal phase MPLC (10→20% EA:Hex) afforded desired compound **2** (55 mg, 85.8 µmol, 80%) as colorless foam. **$^1$H-NMR (500 MHz, $CDCl_3$):** δ = 5.01 (dd, $J$ = 9.9, 2.1 Hz, 1H), 4.04 (dd, $J$ = 10.0, 2.3 Hz, 1H), 3.91 (d, $J$ = 9.4 Hz, 1H), 3.02 (dq, $J$ = 9.2, 6.7 Hz, 1H), 2.91 (dd, $J$ = 16.3, 2.4 Hz, 1H), 2.84 (dd, $J$ = 10.1, 4.0 Hz, 1H), 2.75 (dd, $J$ = 16.4, 10.0 Hz, 1H), 2.35 (ddd, $J$ = 15.0, 4.1, 2.1 Hz, 1H), 2.22 (s, 3H), 1.70 (ddt, $J$ = 18.5, 10.1, 4.9 Hz, 2H), 1.63 – 1.53 (m, 2H), 1.52 – 1.43 (m, 2H), 1.38 (d, $J$ = 12.1 Hz, 1H), 1.27 (s, 3H), 1.22 (s, 3H), 1.15 (s, 3H), 1.07 (d, $J$ = 6.8 Hz, 3H), 1.05 – 1.01 (m, 1H), 0.99 – 0.96 (m, 12H), 0.91 (t, $J$ = 7.8 Hz, 9H), 0.64 (q, $J$ = 7.8 Hz, 6H), 0.58 (q, $J$ = 7.8 Hz, 6H) ppm. **$^{13}$C-NMR (125 MHz, $CDCl_3$):** δ = 215.2, 203.4, 171.8, 80.2, 76.5, 76.2, 62.5, 62.2, 53.4, 48.5, 39.4, 36.8, 32.0, 31.0, 30.3, 25.9, 24.9, 24.6, 23.6, 22.6, 19.6, 17.8, 7.3, 7.0, 5.7, 5.3 ppm. **$R_f$** = 0.53 (EA:Hex = 2:8). **LRMS (ESI, positive):** calc for $C_{34}H_{65}O_7Si_2^+$ $[M+H]^+$ 641.4263, 641.3 found.

**Synthesis of deepoxidised TES protected methyl ketone 3.**

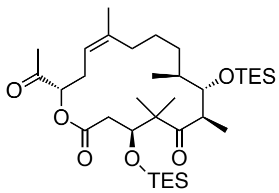

A solution of tungsten hexachloride (27 g, 0.069 mmol, 2.0 eq) in dry THF (2 mL) was cooled to –78°C and n-BuLi (2.5 M in hexanes, 55 µL, 0.137 mmol, 4.0 eq) was added dropwise. The suspension was stirred for 10 min, before the reaction mixture was warmed to 25°C and stirred another 30 min. The black suspension was cooled to –20°C, and a solution of methyl ketone **2** (0.22 mg, 0.034 mmol,1.0 eq) in tetrahydrofuran (1 mL) was added dropwise, and the reaction mixture was warmed to 0°C. After 3 h, the reaction mixture was quenched with sat. aq. $NH_4Cl$ (5 mL), and allowed to warm to room temperature. The reaction mixture was transferred into a separation funnel and the two phases were separated. The aqueous layer was extracted with ethyl acetate (3 × 5 mL). The combined organic layers were dried over $Na_2SO_4$, filtered and concentrated under reduced pressure. The obtained residue was purified by MPLC purification (0→15% EA:Hex) to afford deepoxidised methyl ketone **3** (10.6 g, 0.017 mmol, 49%) as a colorless oil.

**$^1$H-NMR (500 MHz, $CDCl_3$):** δ = 5.16 (t, *J* = 8.2 Hz, 1H), 4.84 (d, *J* = 9.7 Hz, 1H), 4.04 (dd, *J* = 10.4, 1.9 Hz, 1H), 3.91 (d, *J* = 9.1 Hz, 1H), 3.08 – 2.96 (m, 1H), 2.92 (dd, *J* = 15.7, 1.8 Hz, 1H), 2.76 (dd, *J* = 16.2, 10.2 Hz, 1H), 2.54 (dt, *J* = 14.7, 9.8 Hz, 1H), 2.42 (t, *J* = 11.0 Hz, 1H), 2.24 (dd, *J* = 14.7, 7.4 Hz, 1H), 2.19 (s, 3H), 1.78 – 1.69 (m, 2H), 1.69 (s, 3H), 1.55 – 1.48 (m, 2H), 1.22 (s, 3H), 1.14 (s, 3H), 1.09 (d, *J* = 6.7 Hz, 3H), 1.07 – 1.02 (m, 2H), 1.01 – 0.96 (m, 12H), 0.88 (t, *J* = 7.9 Hz, 9H), 0.65 (q, *J* = 7.9 Hz, 6H), 0.56 (q, *J* = 7.8 Hz, 6H) ppm. **$R_f$** = 0.58 (EA:Hex = 15:85). **LRMS (ESI, positive):** calc for $C_{34}H_{65}O_7Si_2^+$ $[M+H]^+$ 641.4263, 641.3 found.

#### Synthesis of styrylthiazole phosphonates

##### Synthesis of styrylthiazole phosphonate 8:

*General remarks: Synthesis of phosphonate photoswitches was performed with as little purification as possible for time efficiency reasons. It is recommended to remove most of the side products after synthesis of* ***6****.*

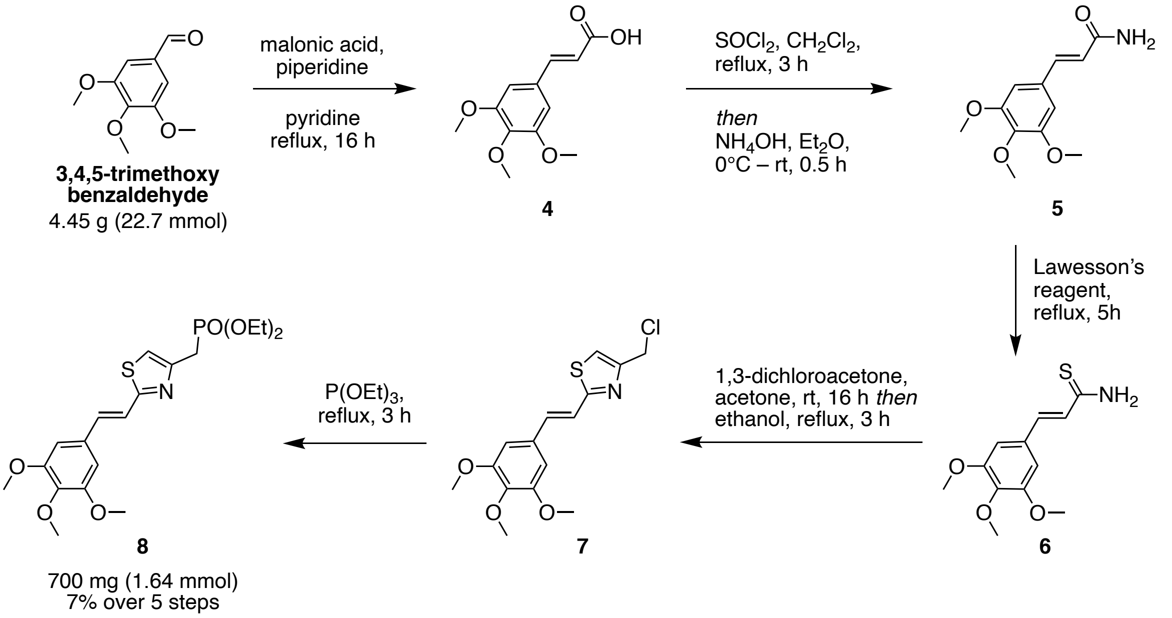

A solution of 3,4,5-trimethoxybenzaldehyde (4.45 g, 22.7 mmol, 1.0 eq.), malonic acid (2.83 g, 27.2 mmol, 1.2 eq) and piperidine (0.5 mL) in pyridine (20 mL) was stirred overnight at reflux. After cooling down to room temperature, the solvent was removed under reduced pressure and redissolved in EA (30 mL). The organic layer was washed with 2 M HCl (15 mL), water

(15 mL) and brine (15 mL), dried over $MgSO_4$ and filtered to afford cinnamic acid **4** (5.40 g, 22.7 mmol, quant.) without further purification.
**$^{1}$H-NMR (400 MHz, $CDCl_3$):** δ = 7.71 (d, *J* = 15.9 Hz, 1H), 6.78 (s, 2H), 6.36 (d, *J* = 15.9 Hz, 1H), 3.90 (s, 6H), 3.90 (s, 3H) ppm. **$^{13}$C-NMR (100 MHz, $CDCl_3$):** δ = 172.1, 153.6, 147.2, 140.7, 129.6, 116.5, 105.7, 61.1, 56.3 ppm. **$R_f$** = 0.57 (EA:Hex:FA = 1:1:0.01). **HRMS (EI, positive):** calc. for $C_{12}H_{14}O_5^{+\bullet}$ $[M]^{+\bullet}$ 238.0836, found 238.0836.

The cinnamic acid **4** (5.40 g, 22.7 mmol, 1 eq) was dissolved in dichloromethane (100 mL). Thionyl chloride (16.5 mL, 227 mmol, 10 eq.) was added slowly and the reaction mixture was stirred at reflux for 3 h. The solvent was removed under reduced pressure and redissolved in $Et_2O$ (80 mL) and $NH_4OH$ (10 mL) was added slowly at 0°C. Stirring was continued for 30 min. The precipitate was washed with $Et_2O$ several times to give cinnamic amide **5** (4.95 g, 20.9 mmol, 91%).
**$^{1}$H-NMR (400 MHz, DMSO-*d₆*):** δ = 7.49 (s, 1H), 7.35 (d, *J* = 15.8 Hz, 1H), 7.06 (s, 1H), 6.88 (s, 2H), 6.57 (d, *J* = 15.8 Hz, 1H), 3.80 (s, 6H), 3.67 (s, 3H) ppm. **$^{13}$C-NMR (100 MHz, DMSO-$d_6$):** δ = 166.9, 153.1, 139.5, 138.7, 130.6, 121.7, 105.0, 60.2, 55.9 ppm. **$R_f$** = 0.23 (EA = 100%). **HRMS (ESI, positive):** calc. for $C_{12}H_{16}NO_4^{+}$ $[M+H]^{+}$ 238.1074, found 238.1075.

The cinnamic amide **5** (5.40 g, 22.7 mmol, 1 eq) was dissolved in THF (100 mL). Lawesson's reagent (4.95 g, 11.3 mmol, 0.5 eq.) was added portionwise and the reaction mixture was stirred at reflux for 5 h. The solvent was removed under reduced pressure and redissolved in dichloromethane (100 mL) transferred into a separation funnel and washed with 2M HCl (3 × 30 mL) and brine (30 mL), dried over $MgSO_4$ and filtered. All volatiles were removed by rotary evaporation and filtered. Purification by normal phase MPLC (40→60% EA:Hex, elutes at 60% EA) to remove most of the Lawesson's reagent byproducts afforded cinnamic thioamide **6** (840 mg, 3.32 mmol, 14% over 3 steps).
**$^{1}$H-NMR (400 MHz, $CDCl_3$):** δ = 7.71 (d, *J* = 15.8 Hz, 1H), 6.78 (s, 2H), 6.36 (d, *J* = 15.8 Hz, 1H), 3.90 (s, 6H), 3.89 (s, 3H) ppm. **$^{13}$C-NMR (100 MHz, $CDCl_3$):** δ = 172.1, 153.6, 147.2, 140.7, 129.6, 116.5, 105.7, 61.1, 56.3 ppm. **$R_f$** = 0.26 (EA:Hex = 1:1). **HRMS (ESI, positive):** calc. for $C_{12}H_{16}NO_3S^{+}$ $[M+H]^{+}$ 254.0845, found 254.0847.

A solution of cinnamic thioamide **6** (840 mg, 3.32 mmol, 1 eq) in acetone (25 mL) was added dropwise to a solution of 1,3-dichloroacetone (421 mg, 3.32 mg, 1 eq) in acetone (25 mL) at 50°C. The reaction mixture was allowed to cool down to room temperature and stirred overnight. The acetone was removed under reduced pressure, redissolved in ethanol (30 ml) and refluxed for 3 h. After cooling down to room temperature the solvent was removed *in* *vacuo* and purified by MPLC (30→40% EA:Hex) to give compound **7** (811 mg, 2.49 mmol, 75%) as slightly yellowish solid.

**$^1$H NMR (400 MHz, CDCl$_3$):** δ = 7.33 (d, *J* = 16.2 Hz, 1H), 7.24 – 7.16 (m, 2H), 6.76 (s, 2H), 4.69 (d, *J* = 0.7 Hz, 2H), 3.90 (s, 6H), 3.88 (s, 3H) ppm. **$^{13}$C-NMR (100 MHz, CDCl$_3$):** δ = 168.0, 153.6, 152.3, 139.5, 136.0, 131.0, 120.1, 116.6, 104.5, 61.1, 56.3, 40.5 ppm. **R$_f$** = 0.38 (EA:Hex = 3:7). **HRMS (ESI, positive):** calc. for $C_{15}H_{17}NO_3ClS^+$ $[M+H]^+$ 326.0612, found 326.0615.

A stirred solution styrylthiazole **7** (811 mg, 2.49 mmol, 1 eq) in triethyl phosphite (6 mL, 35 mmol, 10.5 eq) was heated to 160 °C. After 3 h, the triethyl phosphite was removed under a steady flow of nitrogen gas, and the reaction mixture was allowed to cool to 25°C. The crude material was purified by normal phase MPLC (20→100% EA:Hex, elutes at 100% EA) to give stryrylthiazole phosphonate **8** (700 mg, 1.64 mmol, 66%) as yellow oil.

**$^1$H-NMR (500 MHz, CDCl$_3$):** δ = 7.30 (d, *J* = 16.1 Hz, 1H), 7.19 (dd, *J* = 16.1, 0.7 Hz, 1H), 7.16 (d, *J* = 3.4 Hz, 1H), 6.75 (s, 2H), 4.11 (dq, *J* = 8.3, 7.1 Hz, 4H), 3.90 (s, 6H), 3.87 (s, 3H), 3.40 (dd, *J* = 21.0, 0.8 Hz, 2H), 1.30 (t, *J* = 7.1 Hz, 6H) ppm. **$^{13}$C-NMR (125 MHz, CDCl$_3$):** δ = 166.5, 153.6, 147.5, 147.4, 139.1, 134.4, 131.4, 121.0, 115.6, 115.6, 104.3, 62.5, 62.4, 61.1, 56.3, 30.3, 29.1, 16.6, 16.5 ppm. **R$_f$** = 0.13 (EA:Hex = 9:1). **HRMS (ESI, positive):** calc. for $C_{19}H_{27}NO_6PS^+$ $[M+H]^+$ 428.1291, found 428.1293.

##### Synthesis of styrylthiazole phosphonate 12a

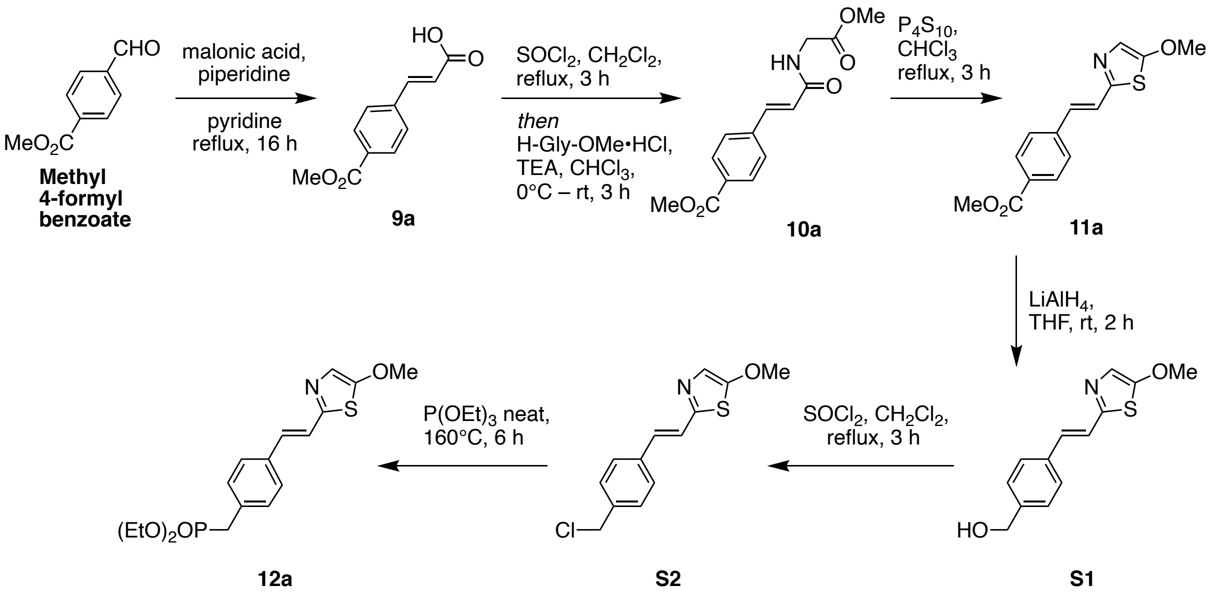

A mixture of methyl 4-formylbenzoate (1.64 g, 10 mmol, 1 eq), malonic acid (1.25 g, 12 mmol, 1.2 eq) and piperidine (99 µL, 1 mmol, 0.1 eq) in pyridine (20 mL) was refluxed overnight. The reaction mixture was cooled down to room temperature, the solvent was removed, redissolved

in EA (30 mL) and transferred into a separation funnel. The organic layer was washed with 2M HCl (2 × 20 mL), water (20 mL) and brine (20 mL). After drying over $MgSO_4$, the solvent was filtered and concentrated *in vacuo* to give crude cinnamic acid **9a** (2.06 g, 10 mmol, quant.) as colorless solid, which was directly resuspended in DCM (30 mL). $SOCl_2$ (7.25 mL, 10 mmol, 10 eq) was added in a dropwise manner. The reaction mixture was stirred at reflux for 1.5 h, cooled down to room temperature and all volatiles were removed by rotary evaporation. The residuals were redissolved in $CHCl_3$ (30 mL) and added slowly to a stirring suspension of glycine methyl ester hydrochloride (1.38 g, 11 mmol, 1.1 eq) and TEA (3.1 mL, 22 mmol, 2.2 eq) at 0°C. The reaction mixture was allowed to warm up to room temperature and stirring was continued for 3 h. The solution was transferred into a separation funnel and washed with water (20 mL) and brine (20 mL), dried over $MgSO_4$, filtered and concentrated under reduced pressure to give crude amide **10a**, which was directly redissolved in chloroform (50 mL) and phosphorus pentasulfide (4.45 g, 10 mmol, 1 eq) was added. The reaction mixture was stirred at 80°C for 3 h after cooling down to room temperature the reaction was carefully quenched with 2M NaOH (100 mL) and extracted with EA (5 x 40 mL). The combined organic layers were dried over $MgSO_4$, filtered and purified by normal phase MPLC (10→40% EA:Hex) to give styrylthiazole **11a** (678 mg, 2.46 mmol, 25% over three steps) as brown solid.
**$^1$H-NMR (500 MHz, $CDCl_3$):** δ = 8.02 (d, *J* = 8.4 Hz, 2H), 7.53 (d, *J* = 8.5 Hz, 2H), 7.22 (s, 1H), 7.11 (d, *J* = 16.3 Hz, 1H), 7.09 (d, *J* = 0.8 Hz, 1H), 3.95 (s, 3H), 3.91 (s, 3H) ppm. **$^{13}$C-NMR (125 MHz, $CDCl_3$):** δ = 166.8, 163.0, 154.3, 140.5, 130.7, 130.2, 129.8, 126.7, 125.0, 122.2, 61.6, 52.3 ppm. **$R_f$** = 0.29 (EA:Hex = 3:7). **HRMS (ESI, positive):** calc. for $C_{14}H_{14}NO_3S^+$ $[M+H]^+$ 276.0689, found 276.0691.

**11a** (552 mg, 2 mmol, 1 eq) was dissolved in dry THF (20 mL) and $LiAlH_4$ (2.21 mL, 2.21 mmol, 1.1 eq) was added dropwise at 0°C. The mixture was stirred at room temperature until complete reduction, and then cooled to 0°C, quenched with ice water, and filtered through Celite. The filtrate was dried over $MgSO_4$ and concentrated *in vacuo* to give benzyl alcohol **S1** (495 mg, 2 mmol, quant.) as colorless solid.
**$^1$H-NMR (400 MHz, $CDCl_3$):** δ = 7.38 (d, *J* = 8.0 Hz, 2H), 7.26 (d, *J* = 7.9 Hz, 2H), 7.01 (d, *J* = 0.9 Hz, 2H), 6.96 (s, 1H), 4.61 (s, 2H), 3.85 (d, *J* = 0.9 Hz, 3H) ppm. **$^{13}$C-NMR (125 MHz, $CDCl_3$):** δ = 162.4, 155.2, 141.4, 135.5, 131.9, 127.6, 127.2, 122.8, 121.8, 65.2, 61.6 ppm. **$R_f$** = 0.45 (EA:Hex = 1:1). **HRMS (ESI, positive):** calc. for $C_{13}H_{14}NO_2S^+$ $[M+H]^+$ 248.0740, found 248.0742.

The benzyl alcohol **S1** (495 mg, 2 mmol, 1 eq) was dissolved in DCM (20 mL) and $SOCl_2$ (0.73 ml, 10.1 mmol, 5 eq) and DMF (0.5 mL) was added at 0°C. The reaction mixture was stirred at room temperature for 2 h before all volatiles were removed under reduced pressure and purified by normal phase MPLC (10→80% EA:Hex, elutes at 40% EA) to give benzyl chloride **S2** (460 mg, 1.73 mmol, 87%).

**$^1$H-NMR (400 MHz, CDCl$_3$):** δ = 7.48 (d, *J* = 8.3 Hz, 2H), 7.39 (d, *J* = 8.3 Hz, 2H), 7.12 (s, 1H), 7.11 (s, 1H), 7.08 (s, 1H), 4.59 (s, 2H), 3.96 (s, 3H) ppm. **$^{13}$C-NMR (125 MHz, CDCl$_3$):** δ = 162.5, 154.8, 137.7, 136.2, 131.3, 129.1, 127.1, 123.3, 121.7, 61.4, 45.9 ppm. **R$_f$** = 0.45 (EA:Hex = 4:6). **HRMS (ESI, positive):** calc. for $C_{13}H_{13}ClNOS^+$ [M+H]$^+$ 266.0401, found 266.0405.

The benzyl chloride **S2** (460 mg, 1.73 mmol, 1 eq) was dissolved in 2 mL triethyl phosphite and stirred at 160°C for 6 h. The solvent was removed over a steady stream of $N_2$ gas. And the remaining crude was purified over normal phase MPLC (10→100% EA:Hex, elutes at 100% EA) to give phosphonate **12a** (470 mg, 1.28 mmol, 74%) as a yellow oil.

**$^1$H-NMR (400 MHz, CDCl$_3$):** δ = 7.43 (dd, *J* = 8.4, 1.2 Hz, 2H), 7.29 (dd, *J* = 8.3, 2.5 Hz, 2H), 7.08 (s, 2H), 7.05 (s, 1H), 4.02 (dqd, *J* = 8.2, 7.1, 2.0 Hz, 4H), 3.94 (s, 3H), 3.15 (d, *J* = 21.9 Hz, 2H), 1.25 (td, *J* = 7.1, 0.5 Hz, 6H) ppm. **$^{13}$C-NMR (100 MHz, CDCl$_3$):** δ = 162.3, 155.0, 134.7, 134.6, 132.2, 132.1, 131.6, 131.6, 130.3, 130.2, 127.0, 127.0, 122.6, 122.6, 121.7, 62.2, 62.2, 61.4, 34.4, 33.0, 16.4, 16.4 ppm. **R$_f$** = 0.14 (EA:Hex = 9:1). **HRMS (ESI, positive):** calc. for $C_{17}H_{23}NO_4PS^+$ [M+H]$^+$ 368.1080, found 368.1083.

**Synthesis of styrylthiazole phosphonate 12b**

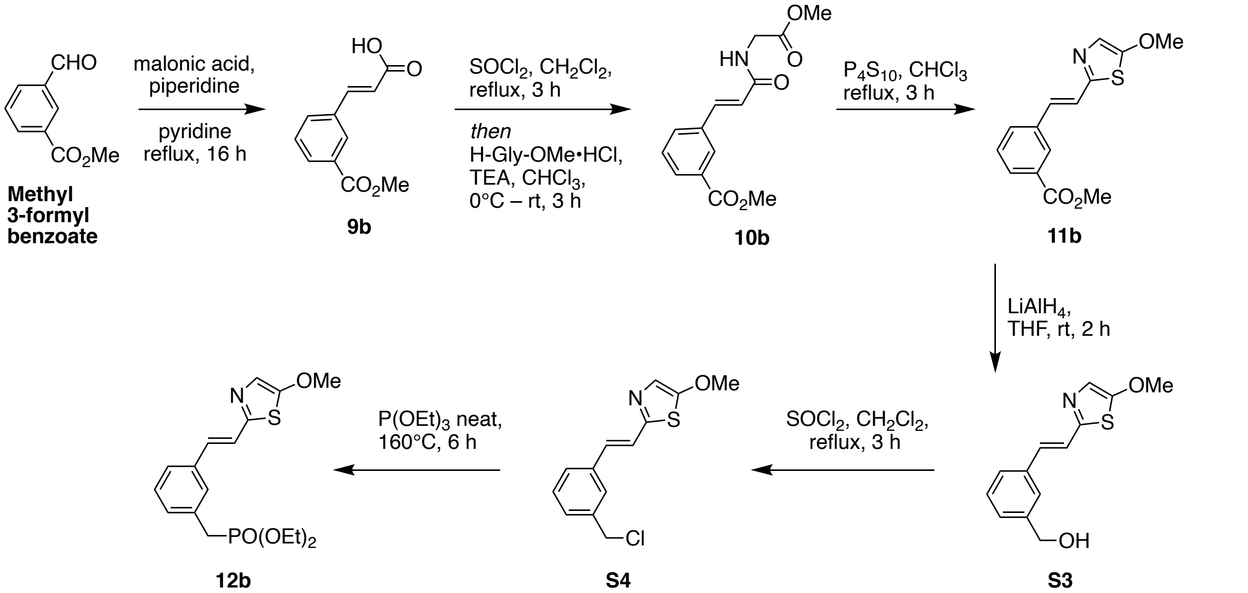

Styrylthiazole **11b** was synthesized following the same procedures for **11a**, starting from Methyl 3-formylbenzoate (2 g, 12.2 mmol, 1 eq) to give **11b** (766 mg, 2.78 mmol, 23% over three steps) as brown solid.

**$^1$H-NMR (400 MHz, $CDCl_3$):** δ = 8.12 (t, *J* = 1.8 Hz, 1H), 7.91 (ddd, *J* = 7.8, 1.7, 1.2 Hz, 1H), 7.64 – 7.58 (m, 1H), 7.39 (t, *J* = 7.8 Hz, 1H), 7.15 (d, *J* = 16.3 Hz, 1H), 7.08 (d, *J* = 16.3 Hz, 1H), 7.05 (s, 1H), 3.91 (s, 3H), 3.90 (s, 3H) ppm. **$^{13}$C-NMR (100 MHz, $CDCl_3$):** δ = 166.7, 162.6, 154.3, 136.3, 130.8, 130.8, 130.7, 129.3, 128.9, 127.9, 123.8, 121.9, 61.4, 52.2 ppm. **$R_f$** = 0.51 (EA:Hex = 1:1). **HRMS (ESI, positive):** calc. for $C_{14}H_{14}NO_3S^+$ $[M+H]^+$ 276.0689, found 276.0687.

Reduction of **11b** (433 mg, 2 mmol, 1 eq) following the same procedure as for benzyl alcohol **S1** gave benzyl alcohol of **S3** (367 mg, 1.48 mmol, 94%).
**$^1$H-NMR (500 MHz, $CDCl_3$):** δ =7.50 (s, 1H), 7.42 (d, *J* = 7.6 Hz, 1H), 7.36 (t, *J* = 7.6 Hz, 1H), 7.30 (d, *J* = 7.5 Hz, 1H), 7.13 (d, *J* = 1.8 Hz, 2H), 7.06 (s, 1H), 4.72 (s, 2H), 3.95 (s, 3H) ppm. **$^{13}$C-NMR (125 MHz, $CDCl_3$):** 162.5, 155.1, 141.6, 136.4, 131.9, 129.2, 127.2, 126.2, 125.4, 123.1, 121.9, 65.3, 61.6 ppm. **$R_f$** = 0.20 (EA:Hex = 3:7). **HRMS (ESI, positive):** calc. for $C_{13}H_{14}NO_2S^+$ $[M+H]^+$ 248.0740, found 248.0739.

Chlorination of benzyl alcohol of **S3** (330 mg, 1.33 mg, 1 eq) was obtained following the same the same procedure as for benzyl chloride **S2** to yield benzyl chloride **S4** (315 mg, 1.19 mmol, 89%).
**$^1$H-NMR (400 MHz, $CDCl_3$):** δ = 7.50 (d, *J* = 1.8 Hz, 1H), 7.45 (dt, *J* = 7.5, 1.6 Hz, 1H), 7.36 (t, *J* = 7.5 Hz, 1H), 7.32 (dt, *J* = 7.7, 1.6 Hz, 1H), 7.15 (d, *J* = 16.2 Hz, 1H), 7.10 (d, *J* = 16.1 Hz, 2H), 4.60 (s, 2H), 3.96 (s, 3H) ppm. **$^{13}$C-NMR (100 MHz, $CDCl_3$):** δ = 162.6, 154.8, 138.2, 136.7, 131.4, 129.4, 128.7, 127.1, 126.8, 123.5, 122.0, 61.6, 46.1 ppm. **$R_f$** = 0.49 (EA:Hex = 3:7). **HRMS (ESI, positive):** calc. for $C_{13}H_{13}ClNOS^+$ $[M+H]^+$ 266.0401, found 266.0400.

Styrylthiazole phosphonate **12b** was synthesized from the benzyl chloride of **S4** (315 mg, 1.19 mmol, 1 eq), following the same procedure as for phosphonate **12a** to obtain desired product **12b** (211 mg, 0.574 mmol, 49%) as yellow oil.
**$^1$H-NMR (400 MHz, $CDCl_3$):** δ = 7.43 – 7.34 (m, 2H), 7.31 (t, *J* = 7.6 Hz, 1H), 7.24 (ddt, *J* = 7.6, 2.9, 1.5 Hz, 1H), 7.10 (s, 1H), 7.10 (s, 1H), 7.06 (s, 1H), 4.03 (dqd, *J* = 8.3, 7.1, 2.1 Hz, 4H), 3.94 (s, 3H), 3.16 (d, *J* = 21.6 Hz, 2H), 1.25 (t, *J* = 7.1 Hz, 6H) ppm. **$^{13}$C-NMR (100 MHz, $CDCl_3$):** δ = 162.4, 154.9, 136.3, 136.3, 132.5, 132.4, 131.8, 130.1, 130.0, 129.1, 129.1, 128.4, 128.4, 125.3, 125.3, 123.0, 121.8, 62.3, 62.2, 61.5, 34.5, 33.1, 16.5, 16.5 ppm. **$R_f$** = 0.12 (EA:Hex = 9:1). **HRMS (ESI, positive):** calc. for $C_{17}H_{23}NO_4PS^+$ $[M+H]^+$ 368.1080, found 368.1083.

#### Synthesis of SBTax

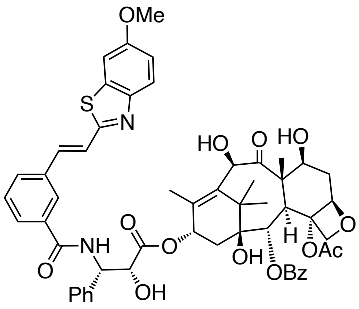

A flask was charged with docetaxel (19 mg, 24 µmol, 1 eq) and DCM (2 mL) and the solution was stirred at 0°C for 2 min. TFA (2 mL) was added and the mixture was stirred at 0°C for 1 hour. The solution was added into rapidly stirred sat. aq. $NaHCO_3$ (15 mL). Solid $NaHCO_3$ was added until all TFA was neutralized. The mixture was extracted with DCM (3 × 10 mL). The combined organic layers were washed with sat. aq. $NaHCO_3$ (10 mL), brine (10 mL), dried on $Na_2SO_4$, filtered and concentrated to give crude deprotected docetaxel as a colorless foam. The crude was dissolved in DMF (1 mL). The SBT photoswitch (22.4 mg, 72 µmol 3 eq) was dissolved in DMF (1 mL), HOBt•$H_2O$ (8.1 mg, 60 µmol, 2.5 eq) and EDCI (8.4 mg, 54 µmol, 2.25 eq) were added and the solution stirred at room temperature for 5 min. DIPEA (25 µL,144 µmol, 6.0 eq) was added and stirring was continued for 10 min. The solution of crude deprotected docetaxel was added and the solution was stirred for 12 h at room temperature, then poured into 10% aq. $NaHCO_3$ (20 mL) and extracted with DCM (3 × 10 mL). The combined organic layers were washed with sat. aq. $NaHCO_3$ (10 mL), sat. aq. LiCl (10 mL), brine (10 mL), dried on $Na_2SO_4$, filtered and concentrated to a yellow solid. Chromatography on silica (EA:Hex = 3:7→1:1 then DCM:MeOH = 1:0→9:1) yielded **SBTax** as a colorless solid (6 mg, 6 µmol, 25%).

**$^1$H-NMR (400 MHz, CDCl$_3$):** δ = 8.14 (d, *J* = 7.3 Hz, 2H), 7.92 (s, 1H), 7.86 (d, *J* = 9.0 Hz, 1H), 7.73 (d, *J* = 7.8 Hz, 1H), 7.67 (d, *J* = 7.8 Hz, 1H), 7.61 (t, *J* = 7.3 Hz, 1H), 7.54 – 7.30 (m, 10H), 7.20 (d, *J* = 9.0 Hz, 1H), 7.07 (dd, *J* = 9.0, 2.5 Hz, 1H), 6.25 (t, *J* = 8.8 Hz, 1H), 5.85 – 5.79 (m, 1H), 5.69 (d, *J* = 7.1 Hz, 1H), 5.20 (s, 1H), 4.96 (d, *J* = 9.3 Hz, 1H), 4.81 (s, 1H), 4.35 – 4.17 (m, 4H), 3.93 (d, *J* = 7.1 Hz, 1H), 3.89 (s, 3H), 3.64 (s, 1H), 2.61 (dt, *J* = 14.9, 8.1 Hz, 1H), 2.42 (s, 3H), 2.31 (t, *J* = 8.8 Hz, 2H), 1.89 (t, *J* = 12.8 Hz, 1H), 1.79 (m, 5H), 1.78 – 1.71 (m, 1H), 1.22 (s, 3H), 1.13 (s, 3H) ppm. **$^{13}$C-NMR (100 MHz, CDCl$_3$):** δ = 211.4, 172.5, 170.9, 167.1, 166.7, 164.2, 158.3, 148.3, 138.1, 136.4, 136.3, 135.9, 135.5, 134.7, 133.9, 130.7, 130.4, 129.5, 129.3, 129.2, 128.9, 128.5, 127.7, 127.2, 125.8, 123.8, 123.6, 116.0, 104.2, 84.3, 81.4, 78.9, 74.9, 74.7, 73.6, 72.5, 72.2, 57.8, 56.0, 55.4, 46.8, 43.2, 37.3, 36.0, 29.9, 26.8, 22.7, 20.6, 14.7, 10.1 ppm. **$R_f$** = 0.22 (DCM:MeOH = 97:3). **HRMS (ESI, positive):** calc. for $C_{55}H_{57}N_2O_{14}S^+$ $[M+H]^+$ 1001.3525, found 1001.3517.

**Synthesis of SBT photoswitch**

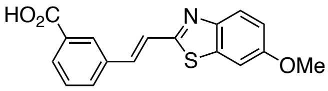

Was prepared following a procedure from Gao et al.[4] from 6-methoxy-2-methylbenzothiazole (358 mg, 2 mmol, 1 eq) and methyl 3-formylbenzoate (328 mg, 2 mmol, 1 eq). A light-yellow solid crashed out of DMSO. The precipitate was collected by filtration and washed with $H_2O$ (2 × 50 mL), acetone (2 × 50 mL) and methanol (2 × 50 mL). No further purification was needed. The product was obtained as a light-yellow solid (468 mg, 1.49 mmol, 75%).

**$^1$H-NMR (400 MHz, DMSO-*d$_6$*):** δ = 8.25 (t, *J* = 1.8 Hz, 1H), 8.03 (dt, *J* = 7.9, 1.5 Hz, 1H), 7.93 (dt, *J* = 7.7, 1.3 Hz, 1H), 7.87 (d, *J* = 8.9 Hz, 1H), 7.68 (d, *J* = 2.6 Hz, 1H), 7.66 – 7.63 (m, 2H), 7.56 (t, *J* = 7.7 Hz, 1H), 7.11 (dd, *J* = 8.9, 2.6 Hz, 1H), 3.85 (s, 3H), 2.54 (s, 1H) ppm. **$^{13}$C-NMR (100 MHz, DMSO-*d$_6$*):** δ =167.0, 163.6, 157.7, 147.9, 135.8, 135.8, 135.3, 131.5, 131.2, 129.8, 129.2, 128.5, 123.2, 123.1, 115.9, 104.8, 55.8 ppm. **$R_f$** = 0.30 (DCM:MeOH = 97:3). **HRMS (EI, positive):** calc. for $C_{17}H_{12}NO_3S^+$ $[M-H]^+$ 310.0532, found 310.0521.

### Part B: Photocharacterization

Absorption spectra in cuvette (“UV-Vis”) were acquired on a Cary 60 UV-Vis Spectrophotometer from Agilent (1 cm pathlength). For photoisomerisation measurements, Hellma microcuvettes (HL105-200-15-40) taking 150 µL volume to top of optical window were used with the test solution concentrations of 10-20 µM. Measurements were performed in pure DMSO or in PBS/DMSO 1:1 indicated by asterisk. Photoisomerisations and relaxation monitoring (**Fig S1c**) were performed at room temperature unless indicated otherwise. Medium-power LEDs (H2A1-models spanning 360–490 nm from Roithner Lasertechnik) were used to deliver high-intensity and relatively monochromatic light (FWHM ~25 nm) into the cuvette, for rapid PSS determinations (**Fig S1a,d**) that were also predictive of what would be obtained in LED-illuminated cell culture. Spectra of pure *E* and *Z* isomers (**Fig S1b**) were acquired from the inline Diode Array Detector during analytical separation on the HPLC (injection of 5-10 µL, 5→100% MeCN:$H_2O$ over 20 min), after a DMSO stock (0.5 – 2.5mM) was irradiated with a 395 nm LED (~ 5 min) or from the inline Diode Array Detector during **STEpo** purification on the prep-HPLC.

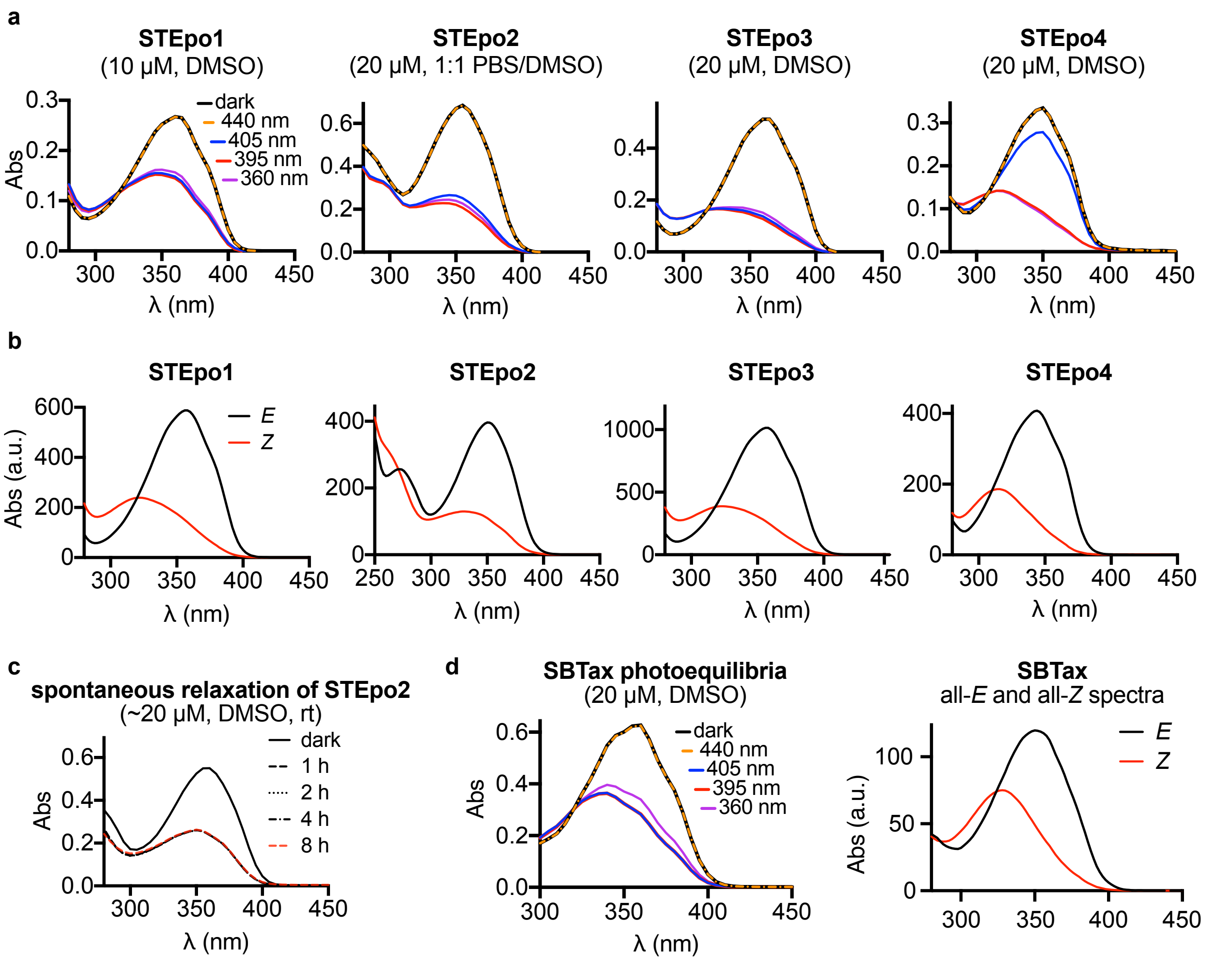

**Fig S1: (a)** UV-Vis spectra of **STEpo**s at various photostationary states, shows that the ST photoswitch scaffold photoisomerizes from an all-*E* isomer state to a *Z*-isomer enriched state when irradiated with UV light (~400 nm). **(b)** all-*E* and all-*Z* spectra obtained from inline HPLC-DAD show that PSS spectra of **STEpos** irradiated with 395 nm photoisomerization **(c) STEpo2** shows no spontaneous relaxation after illumination with 395 nm light at room temperature over 8 hours in DMSO. **(d)** Photocharacterization of **SBTax**.

### Part C: Biological Data

#### Tubulin polymerisation

99% tubulin from porcine brain was obtained from Cytoskeleton Inc. (cat. #T240). The polymerisation reaction was performed at 5 mg/mL tubulin, in polymerisation buffer BRB80 (80 mM piperazine-N,N′-bis(2-ethanesulfonic acid) (PIPES) pH = 6.9; 0.5 mM EGTA; 2 mM $MgCl_2$), in a cuvette (120 µL final volume, 1 cm path length) in a Agilent CaryScan 60 with Peltier cell temperature control unit maintained at 37°C; with glycerol (10 µL). Tubulin was first incubated for 10 min at 37°C with “lit”- (395 nm-pre-illuminated; mostly-*Z*-), dark- (all-*E*) **STEpo** (final concentration 1 µM) or docetaxel (positive ctrl, 10 µM) in buffer with 3% DMSO, without GTP. Then GTP was added to achieve final GTP concentration 1 mM (with mixing), and the change in absorbance at 340 nm was monitored for 15 min, scanning at 15 s intervals.

#### General cell culture

HeLa and Jurkat cells were maintained under standard cell culture conditions in Dulbecco's modified Eagle's medium (DMEM; PAN-Biotech: P04-035550) supplemented with 10% fetal calf serum (FCS), 100 U/mL penicillin and 100 U/mL streptomycin. Cells were grown and incubated at 37°C in a 5% $CO_2$ atmosphere. Cells were typically transferred to phenol red free medium prior to assays (DMEM; PAN-Biotech: P04-03591). Substrates and cosolvent (DMSO; 1% final concentration) were added *via* a D300e digital dispenser (Tecan). Cells were either incubated under "lit" or "dark" conditions; "lit" indicates a pulsed illumination protocol was applied by multi-LED arrays to create the *Z*-isomers of the compounds in *situ* in cells and then maintain the wavelength-dependent PSS isomer ratio throughout the experiment, as described previously.[4] Typical "lit" timing conditions were 75 ms pulses of ~1 mW/cm$^2$ applied every 15 s. "Dark" indicates that **STEpo** biostocks were used with an all-*trans* state, as determined by analytical HPLC, kept in the dark at 4°C, applied while working under red-light conditions, and cells were then incubated in light-proof boxes to shield from ambient light, thereby maintaining the all-*E*-isomer population throughout the experiment.

#### Resazurin antiproliferation assay

HeLa cells were seeded in 96-well plates at 5,000 cells/well and left to adhere for 24 h before treating with test compounds. *E*-**STEpos** were added for 48 h (final well volume 100 µL, 1% DMSO; three technical replicates); the "cosolvent control" ("ctrl") indicates treatment with DMSO only. Cells were then treated with Resazurin 150 mg/mL for 3 h. Fluorescence was measured at 590 nm (excitation 544 nm) using a FLUOstar Omega microplate reader (BMG Labtech). Absorbance data was averaged over the technical replicates, then normalized to viable cell count from the cosolvent control cells (%control) as 100%, where 0% viability was assumed to correspond to fluorescence signal in PBS only with no cells. Three independent experiments were performed. Data were plotted against the log of **STEpo** concentration ($log_{10}$([**STEpo**]) (M)) with mean ±SD.

#### Immunofluorescence staining

HeLa cells were seeded on glass coverslips in 24-well plates (50,000 cells/well) and treated with **STEpo2** the next day under "dark" or "lit" conditions for 24 h. Cells were washed with pre-warmed (37°C) MTSB buffer (80 mM PIPES pH 6.8; 1 mM $MgCl_2$, 5 mM ethylene glycol tetraacetic acid (EGTA) dipotassium salt; 0.5% Triton X-100) for 30 s then fixed with 0.5% glutaraldehyde for 10 min. After quenching with 0.1% $NaBH_4$ (7 min), samples were blocked with PBS + 10% FCS (30 min). The cells were treated with primary antibody (1:400 rabbit alpha-tubulin; Abcam ab18251) in PBS containing 10% FCS for 1 h and then washed with

PBS. Cells were incubated with secondary antibody (1:400 goat-anti-rabbit Alexa fluor 488; Abcam ab150077) in PBS containing 10% FCS for 1 h. After washing with PBS, the coverslips were mounted onto glass slides using Roti-Mount FluorCare DAPI (Roth) and imaged with a Zeiss LSM Meta confocal microscope (CALM platform, LMU). Images were processed using the free Fiji software. Postprocessing was only performed to improve visibility.

#### *In vitro* microtubule dynamics imaging

*Reagents:* GMPCPP was obtained from Jena Biosciences. Biotinylated poly(l-lysine)-[g]-poly(ethylene glycol) (PLL-PEG-biotin) was obtained from Susos AG. NeutrAvidin was obtained from Invitrogen. Ethylene glycol-bis(2-aminoethylether)-N,N,N',N'-tetraacetic acid (EGTA), potassium chloride, potassium hydroxide, κ-casein, 1,4-piperazinediethanesulfonic acid (PIPES), GTP, methylcellulose, glucose oxidase from Aspergillus niger, catalase from bovine liver, dithiothreitol (DTT), magnesium chloride and glucose were obtained from Sigma-Aldrich. Different types of labeled and unlabeled purified tubulin used in the assays were purchased from Cytoskeleton.

The *in vitro* microtubule dynamics assay was performed as described previously (Bieling et al., 2007; Rai et al., 2020)[5,6]. GMPCPP (a slowly hydrolyzable GTP analog) stabilized microtubule seeds were prepared by two rounds of polymerization and depolymerization in the presence of GMPCPP. A solution of 20 µM porcine brain tubulin mix containing 12% rhodamine-labeled tubulin and 18% biotin-labeled tubulin was polymerized in MRB80 buffer (80 mM K-PIPES, pH 6.8, 4 mM $MgCl_2$, 1 mM EGTA) in the presence of GMPCPP (1 mM) at 37°C for 30 min. After polymerization, the mixture was pelleted by centrifugation at 119,000 g for 5 min in an Airfuge centrifuge. The pellet obtained was resuspended in MRB80 buffer and microtubules were depolymerized further on ice for 20 min. The resuspended mixture was again polymerized in the presence of GMPCPP. After the second round of polymerization and pelleting, GMPCPP-stabilized microtubule seeds were stored in MRB80 in the presence of 10% glycerol.

Microscopy slides with plasma-cleaned glass coverslips were used to prepare *in vitro* flow chambers using two strips of double-sided tape. Flow chambers were sequentially incubated with 0.2 mg/mL PLL-PEG-biotin and 1 mg/mL NeutrAvidin in MRB80 buffer. The chamber was further incubated with GMPCPP stabilized microtubule seeds followed by treatment with 1 mg/ml κ-casein in MRB80 buffer. The reaction mixtures containing 15 µM porcine brain tubulin supplemented with 3% rhodamine-tubulin, 20 nM mCherry-EB3, 0.1% methylcellulose, 0.2 mg/mL κ-casein, 50 mM KCl, 1 mM GTP and oxygen scavenger mixture (50 mM glucose, 400 µg/mL glucose oxidase, 200 µg/mL catalase, and 4 mM DTT in MRB80 buffer) without or with **STEpo2** (50 nM) were added to the flow chambers after centrifugation in an Airfuge for

5 minutes at 119,000 g. The chambers were sealed with vacuum grease and microtubule dynamics was recorded at 30 °C using TIRF microscopy.

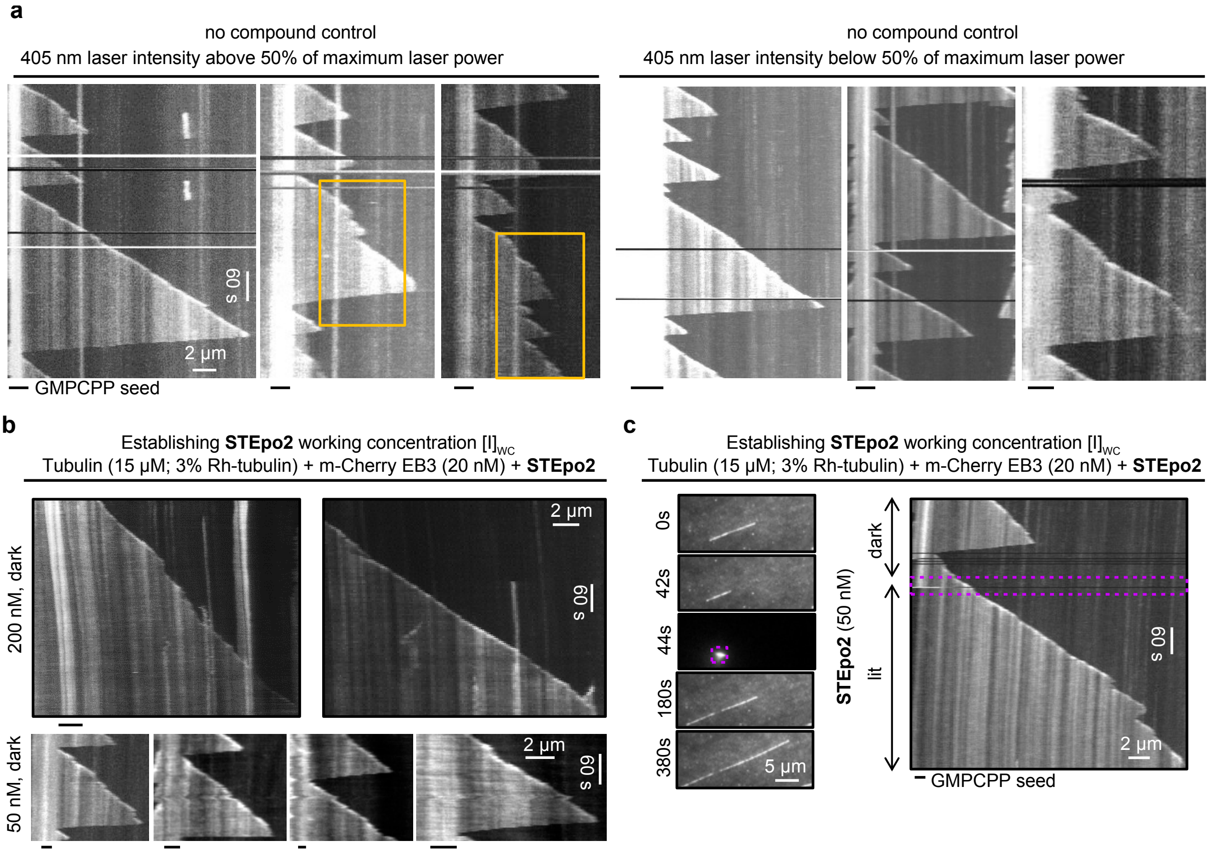

**Fig S2: (a)** Representative kymographs showing MT dynamics during control conditions with different laser intensities of 405 nm. The assay was performed in the presence of tubulin (15 µM supplemented with 3% of rhodamine tubulin) and 20 nM m-Cherry EB3. MT were showing some random rescues or catastrophes (yellow boxes) when the laser was used above 50% of maximum laser power. No significant change in microtubule dynamics was observed when the laser was used below 50% of maximum laser power. A laser intensity of 20-30% of maximum laser power was used for the activation of **STEpo2** (two independent experiments). **(b)** Representative kymographs (two independent experiments) showing microtubule dynamics in the presence of different concentrations of **STEpo2** without 405 nm laser illumination (dark). MT stabilizing dark activity of *E*-**STEpo2** is observed at 200 nM. When decreasing **STEpo2** concentration to 50 nM, MT behave similar to control. 50 nM of STEpo2 was selected as viable working concentration $[I]_{WC}$ for light dependent MT stabilization. **(c)** Time-lapse images and representative kymographs (from three independent experiments) show MT dynamics before (dark) and after (lit) 405 nm photoactivation (20% of maximum laser power, purple box). MT growth events were similar to control conditions in the presence of *E*-**STEpo2** (catastrophes reaching to GMPCPP seeds). After 405 nm photoactivation, MT was showing processive growth event with spontaneous rescues indicating activation of microtubule stabilization by *Z*-**STEpo2**.

#### Image acquisition by TIRF microscopy

Imaging was performed on a TIRF microscope setup (inverted research microscope Nikon Eclipse Ti-E) which was equipped with a perfect focus system (PFS) (Nikon) and Nikon CFI Apo TIRF 100x 1.49 N.A. oil objective (Nikon, Tokyo, Japan). The microscope was supplemented with TIRF-E motorized TIRF illuminator modified by Roper Scientific France/PICT-IBiSA Institut Curie, and a stage top incubator model INUBG2E-ZILCS (Tokai Hit) was used to regulate the temperature of the sample. Image acquisition was performed using Prime BSI sCMOS camera (Teledyne Photometrics, final magnification 0.068 µm/pixel) and controlled with MetaMorph 7.7 software (Molecular Devices, CA). Images were captured with 1 frame/2 s in time-lapse mode.

A TIRF microscope equipped with an ILas system (Roper Scientific/PICTIBiSA) was used to photoactivate **STEpo2** with 405 nm light. A region of microtubules was illuminated using a focused laser beam. *In vitro* microtubule dynamics assay was performed in the presence of GMPCPP-stabilized microtubule seeds with 15 µM tubulin, supplemented with 3% rhodamine-tubulin without (control) or with 50 nM **STEpo2**. Kymographs representing the life history of microtubule dynamics were generated using KymoResliceWide v.0.4 (https://github.com/ekatrukha/KymoResliceWide (Katrukha, 2015)) plugin in ImageJ.

#### Live cell microtubule dynamics: imaging

HeLa cells were transfected with EB3-tdTomato using FuGENE 6 (Promega) according to manufacturer's instructions. Experiments were imaged on a Nikon Eclipse Ti microscope equipped with a perfect focus system (Nikon), a spinning disk-based confocal scanner unit (CSU-X1-A1, Yokogawa), an Evolve 512 EMCCD camera (Photometrics) attached to a 2.0× intermediate lens (Edmund Optics), a Roper Scientific custom-made set of Vortran Stradus 405 nm (100 mW), 488 nm (487 nm / 150 mW) and Cobolt Jive 561 nm (110 mW) lasers, a set of ET-DAPI, ET-GFP and ET-mCherry filters (Chroma), a motorized stage MS-2000-XYZ, a stage top incubator INUBG2E-ZILCS (Tokai Hit) and lens heating calibrated for incubation at 37°C with 5% $CO_2$. Microscope image acquisition was controlled using MetaMorph 7.7 and images were acquired using a Plan Apo VC 40× NA 1.3 oil objective. Imaging conditions were initially optimized to minimize tdTomato bleaching and phototoxicity for untreated cells. For compound treated acquisitions 0.6 µM **STEpo2** diluted in prewarmed cell medium was applied to cells in a dark room with only red ambient light, cells were protected from all ambient light after application of drug. Drug was incubated on cells for at least 1 min before commencing experiment. Comet count analysis was performed in ImageJ using the ComDet plugin (E. Katrukha, University of Utrecht, Netherlands, https://github.com/ekatrukha/ComDet).

#### Live cell microtubule dynamics: quantification and statistics

All relevant assays were done in independent biological replicates. All attempts at replication were successful and no data were excluded from analysis. Blinding was not performed as assay readout is mostly unbiased (plate reader, flow cytometry, Fiji/ImageJ plugins). Microscopic evaluation was performed independently by two separate scientists. Data were analysed using Prism 9 software (GraphPad). Two-tailed unpaired t tests were used in pairwise group comparisons; * was used for $P < 0.05$, ** for $P < 0.01$, *** for $P < 0.001$, **** for $P < 0.0001$.

### Part D: NMR Spectra

#### [1]H-NMR of TES protected STEpo1:

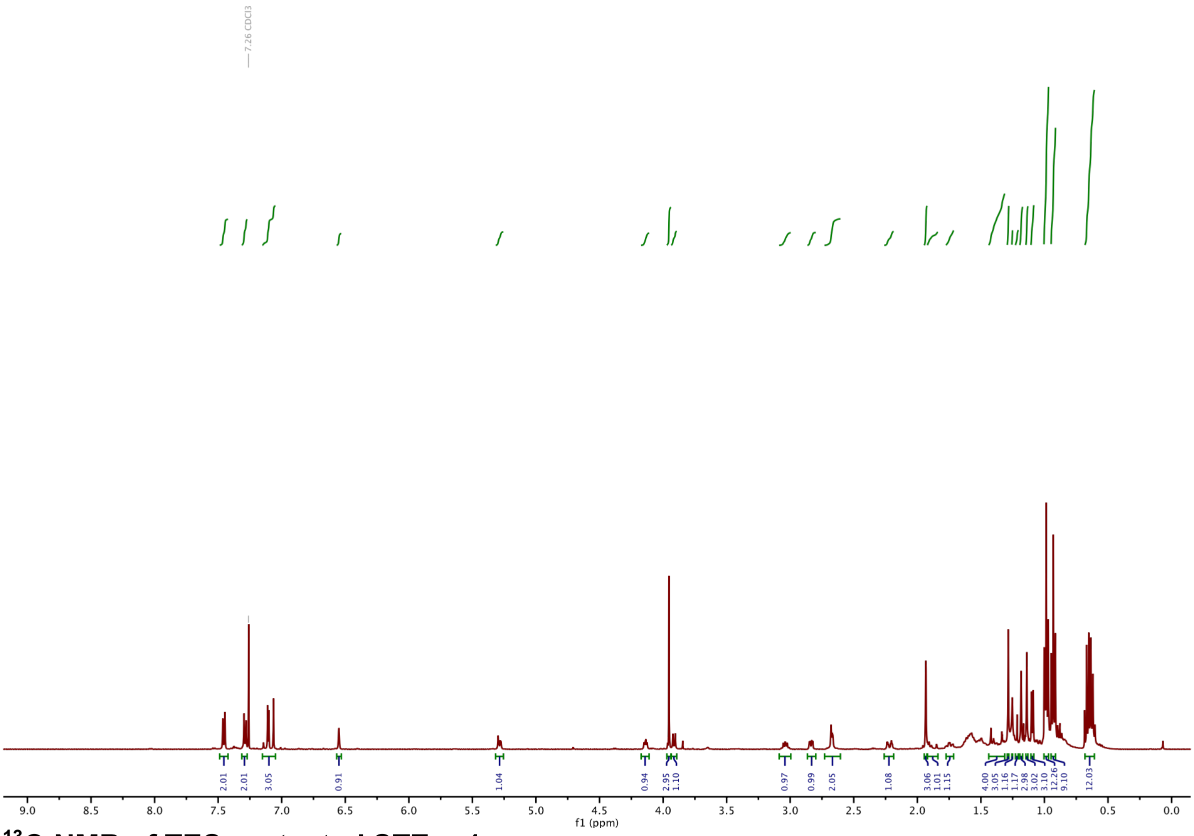

#### [13]C-NMR of TES protected STEpo1:

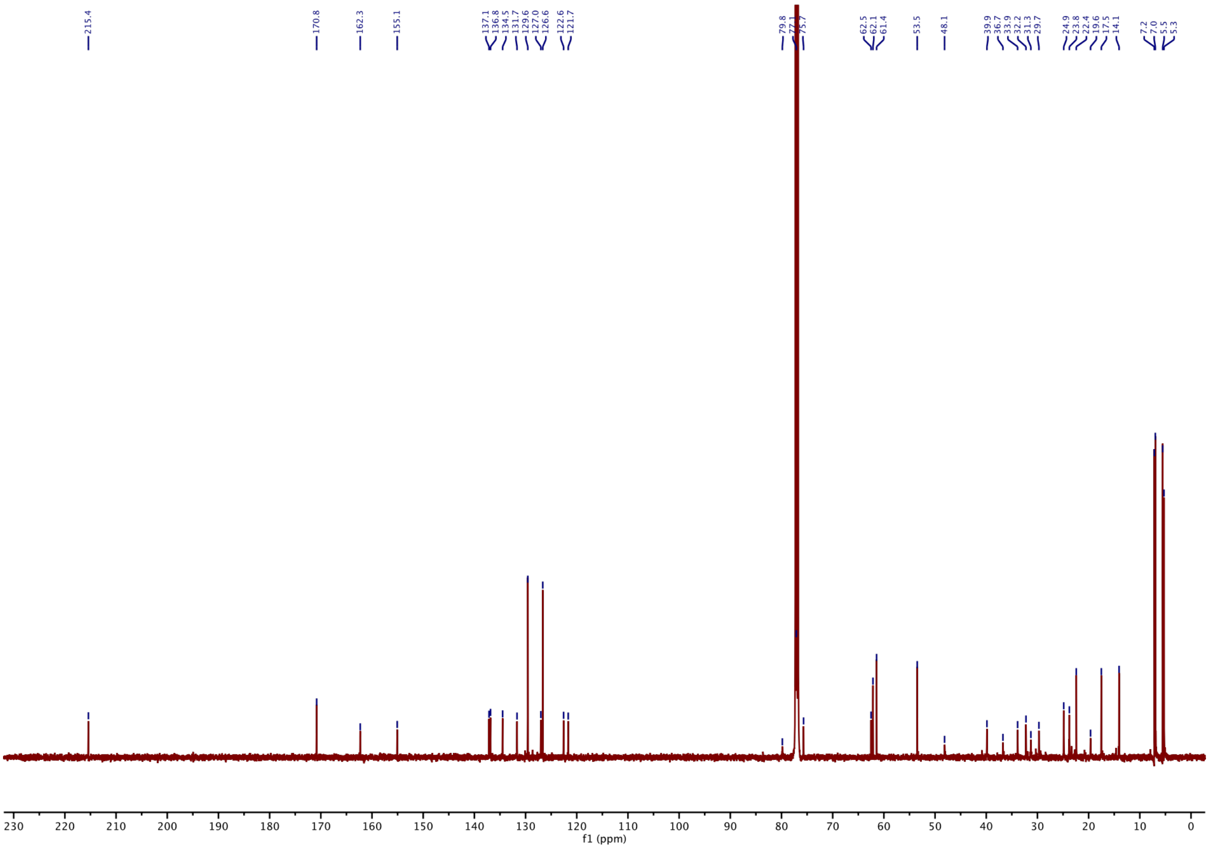

#### $^{1}$H-NMR of STEpo1:

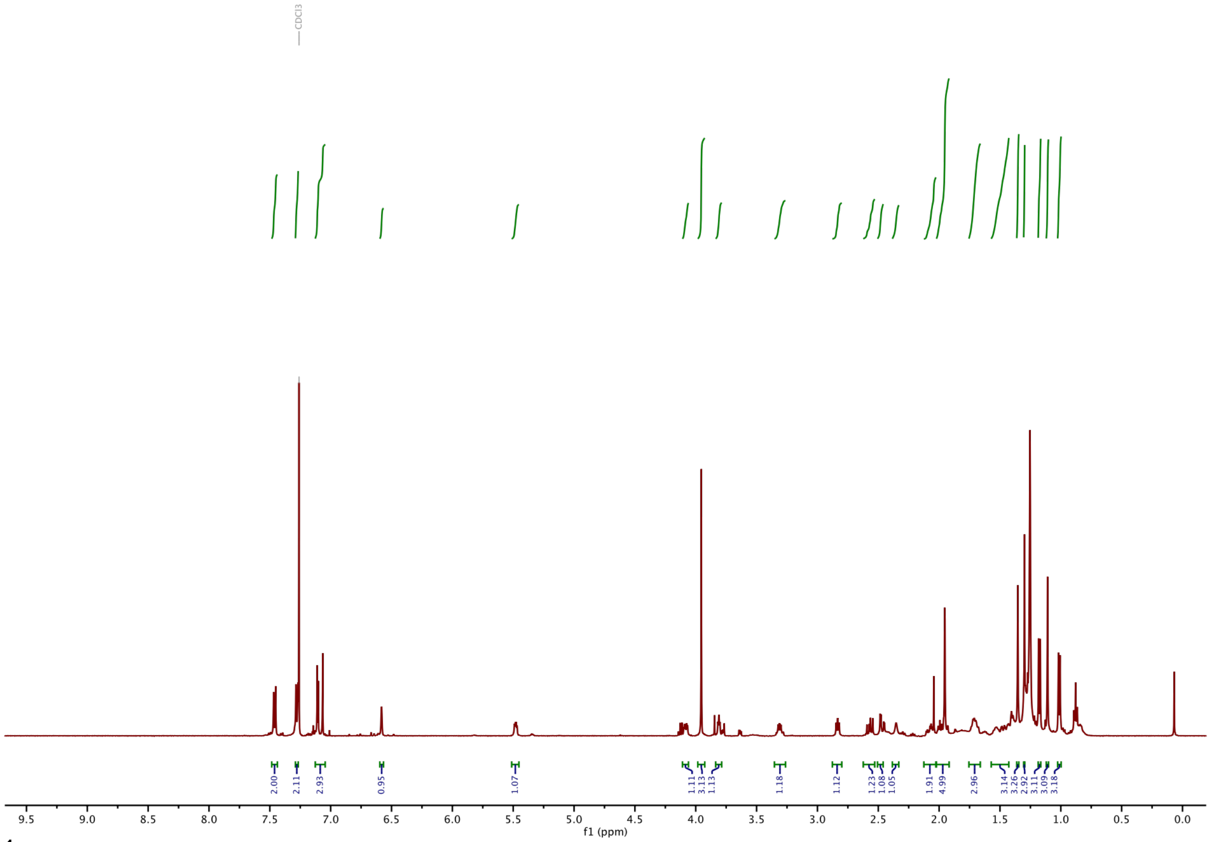

#### $^{1}$H-NMR of TES protected STEpo2:

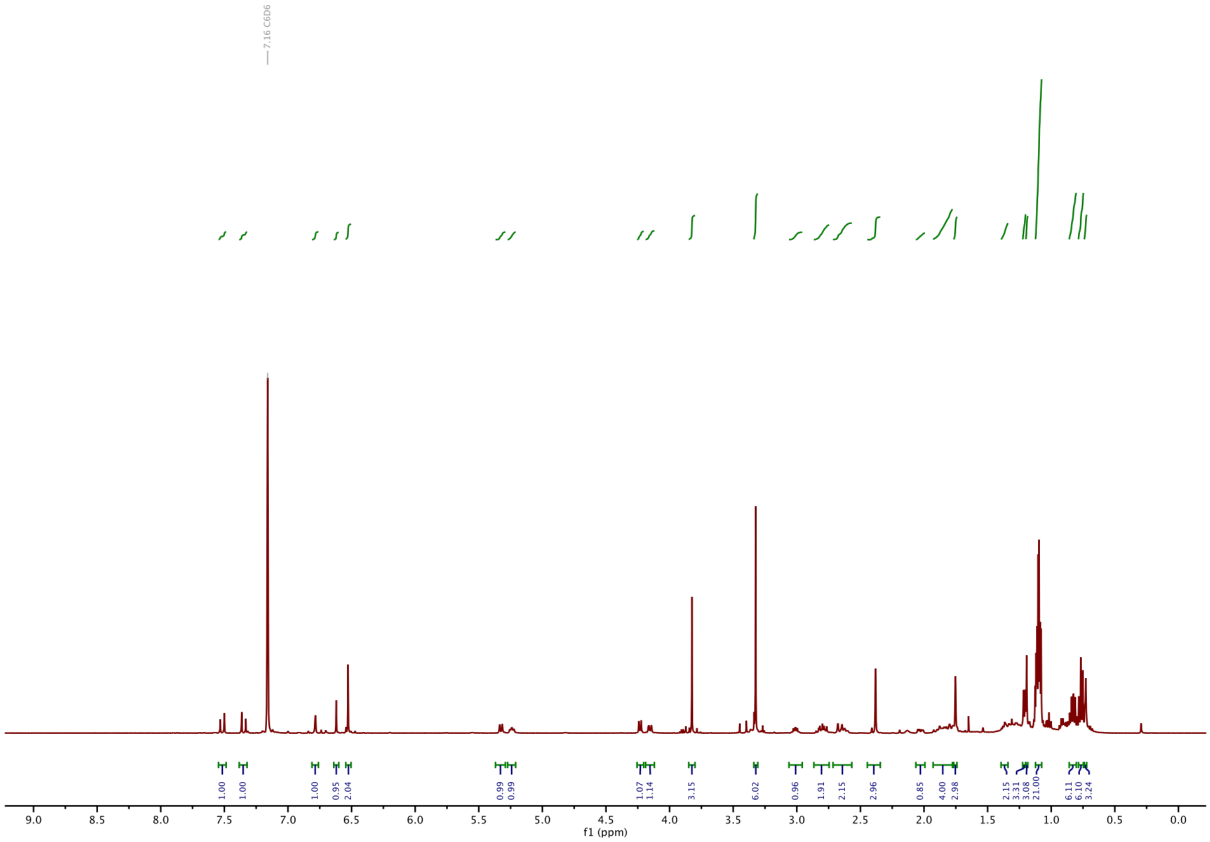

#### $^{13}$C-NMR of TES protected STEpo2:

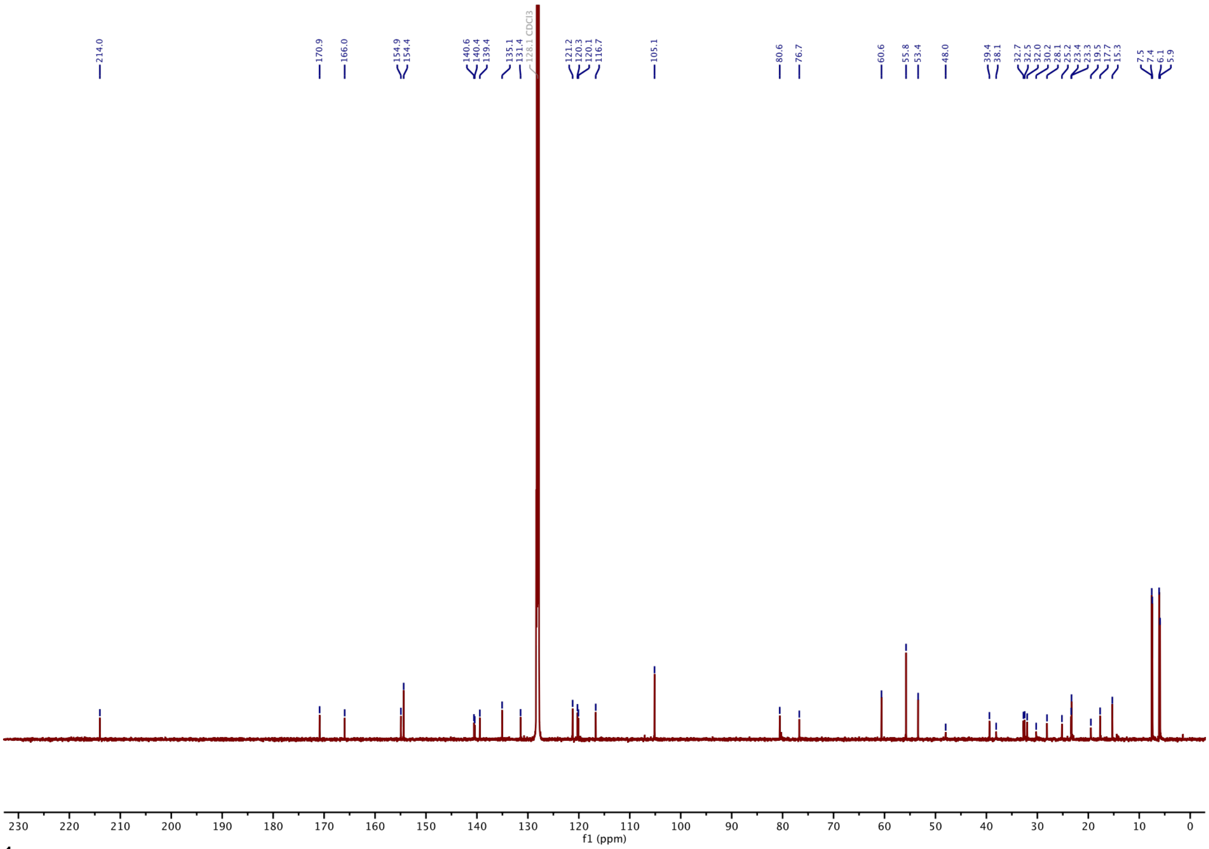

#### $^{1}$H-NMR of STEpo2:

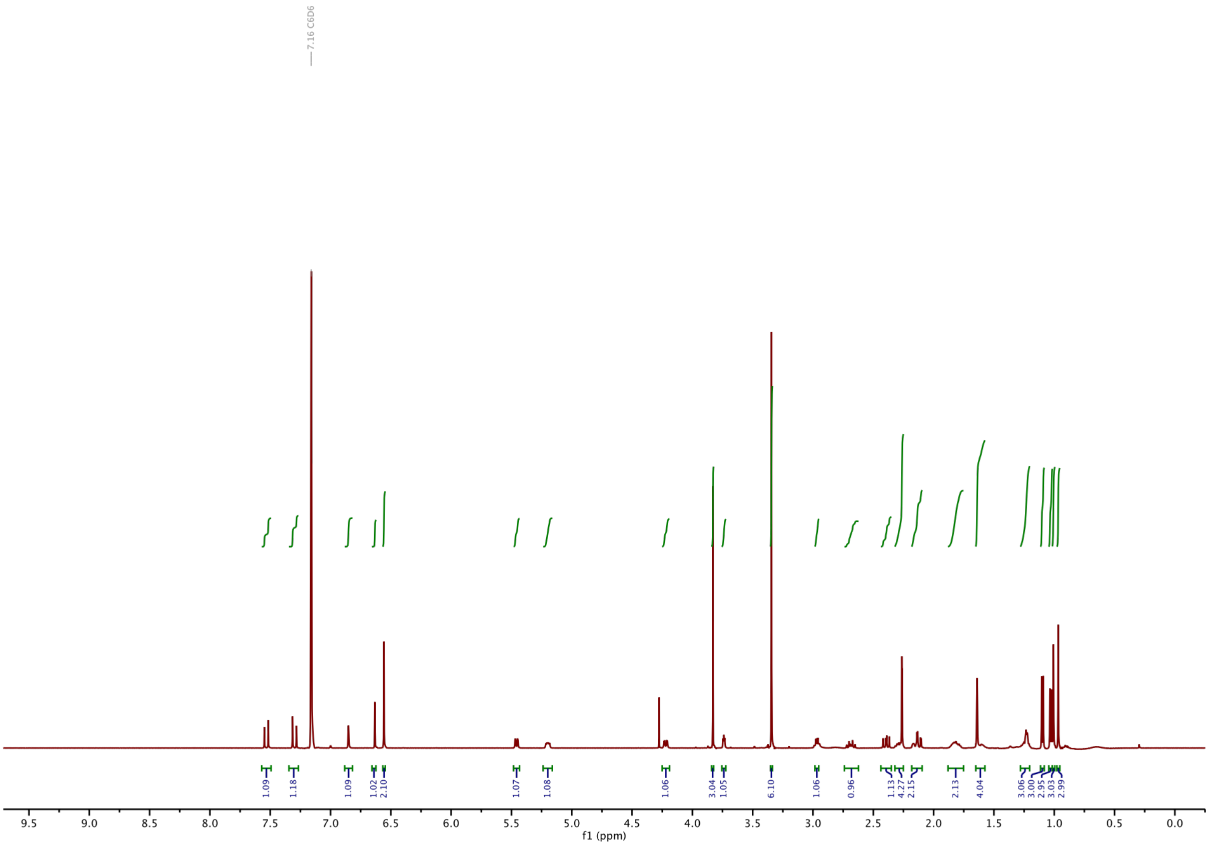

#### $^{13}$C-NMR of STEpo2:

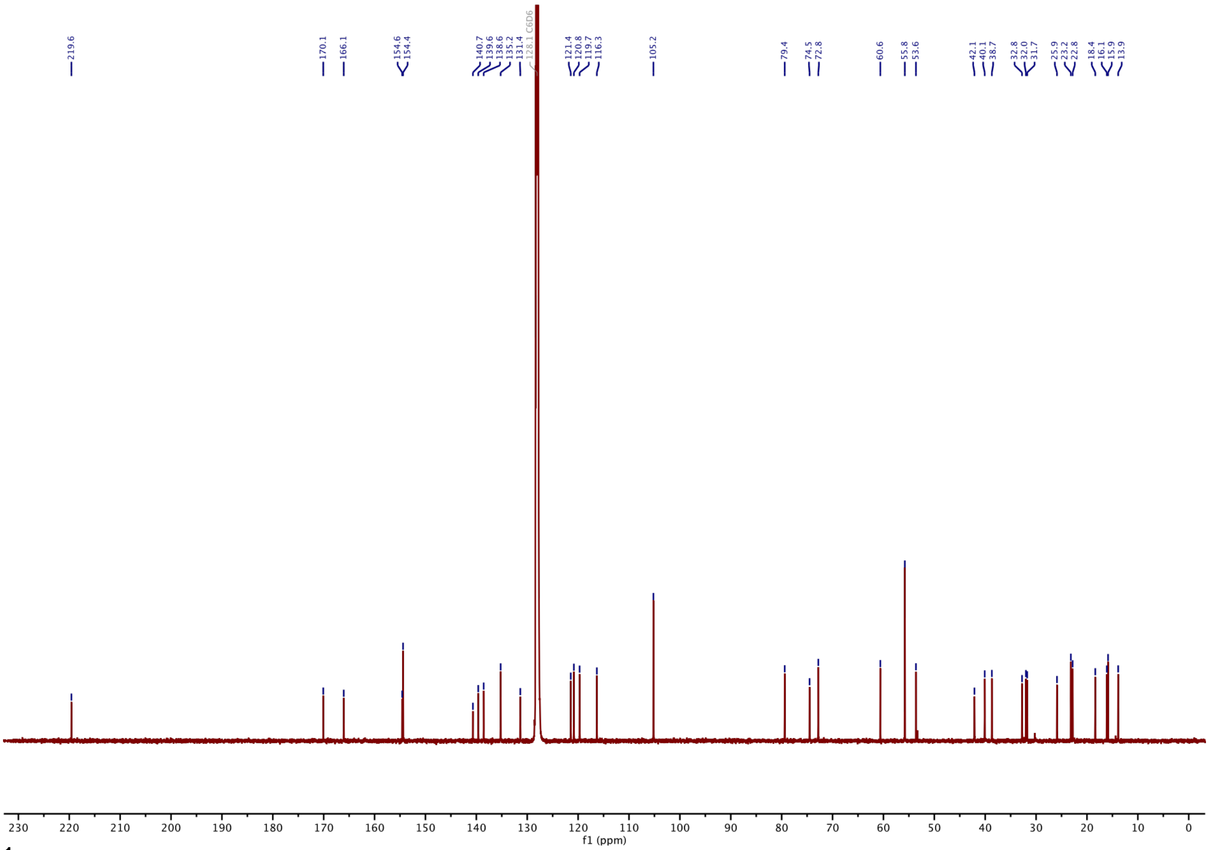

#### $^{1}$H-NMR of TES protected STEpo3:

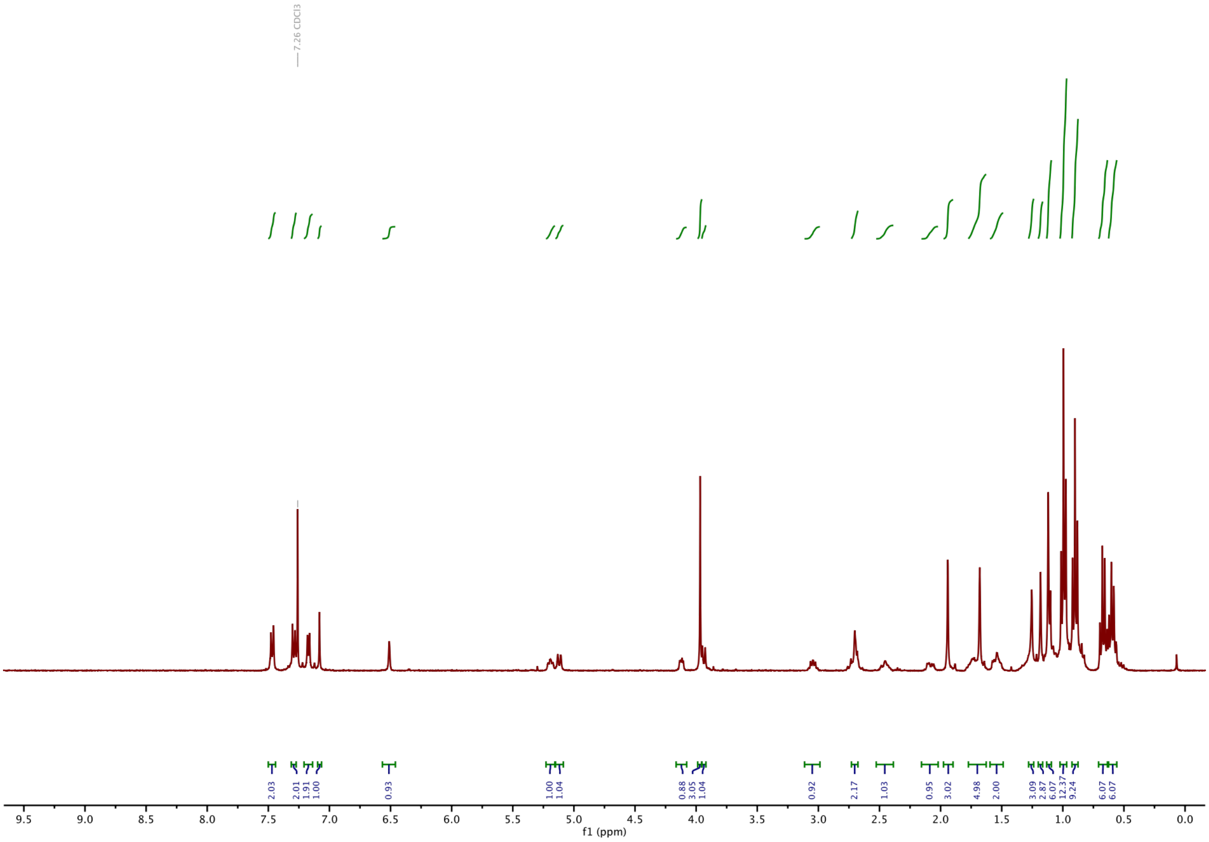

**[13]C-NMR of TES protected STEpo3:**

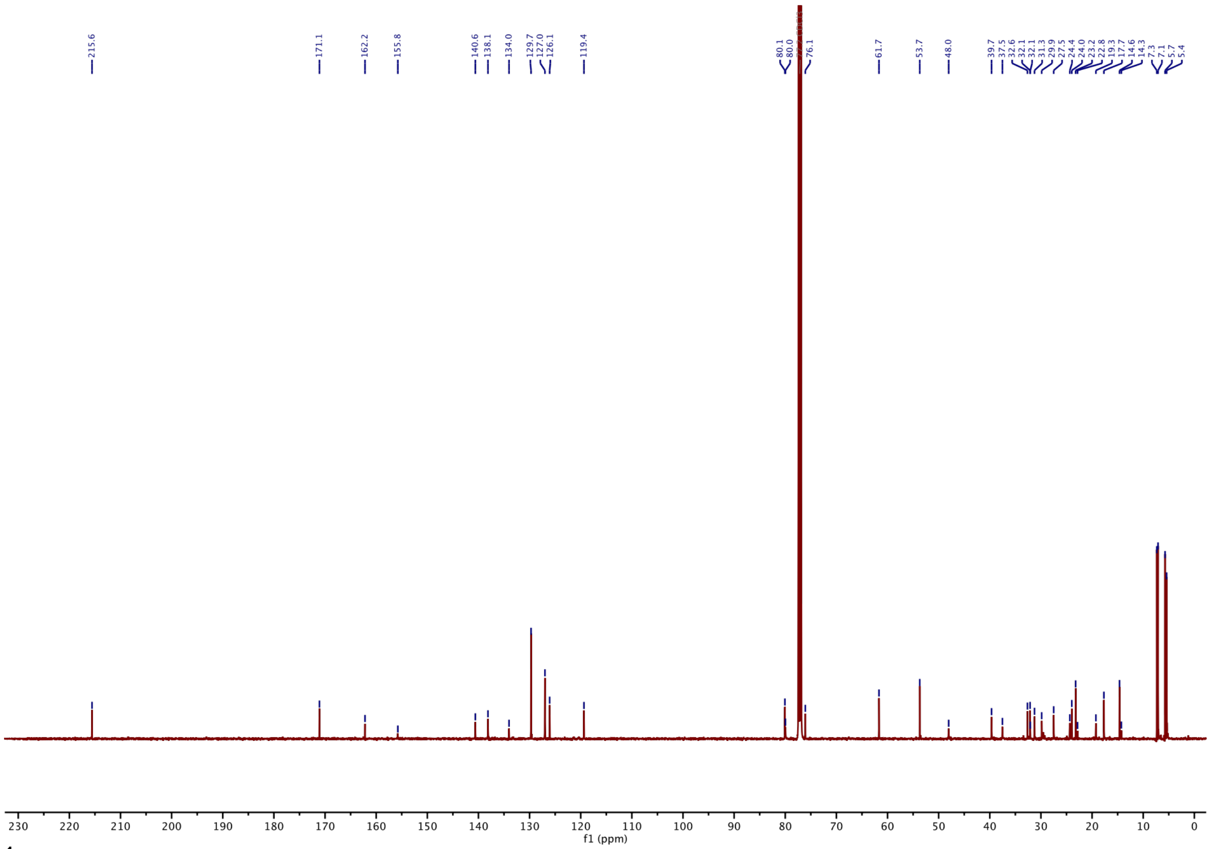

**[1]H-NMR of STEpo3:**

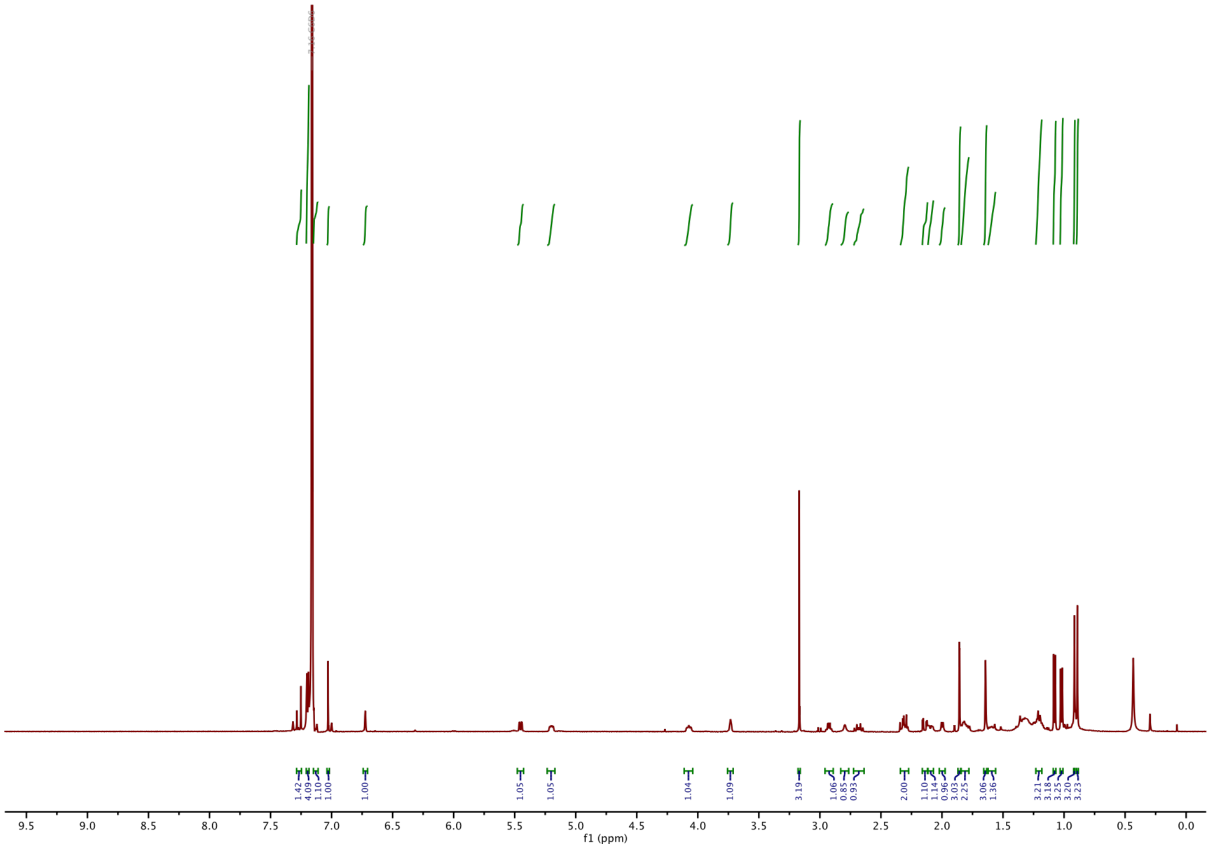

#### $^{13}$C-NMR of STEpo3:

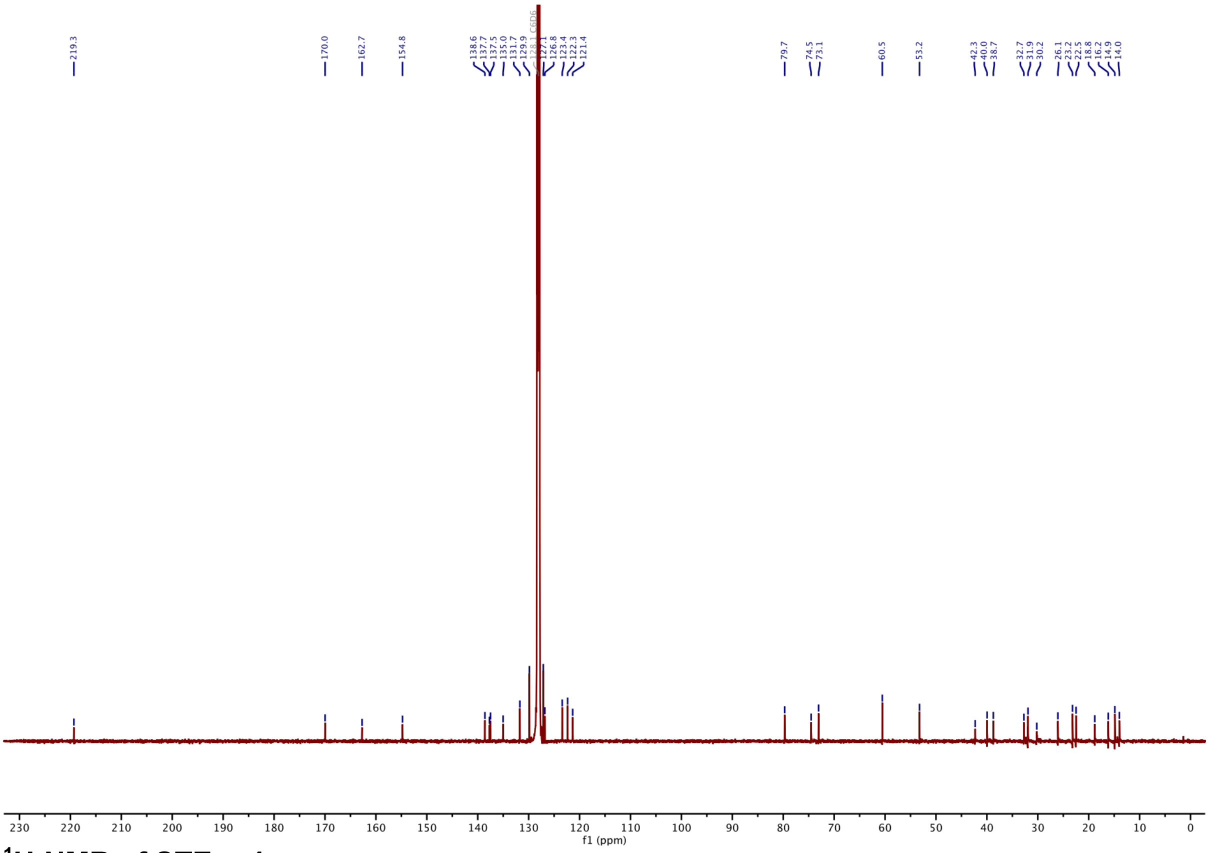

#### $^{1}$H-NMR of STEpo4:

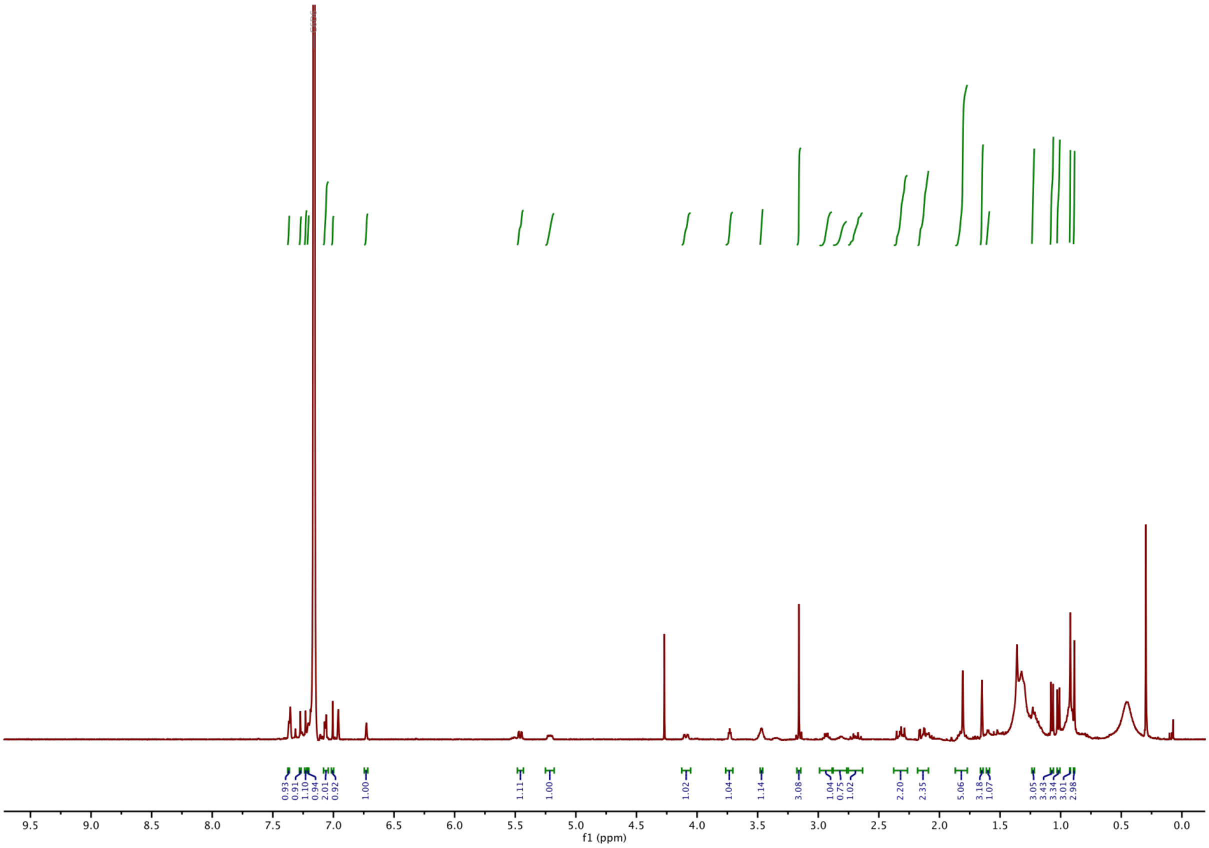

**[1]H-NMR of 1:**

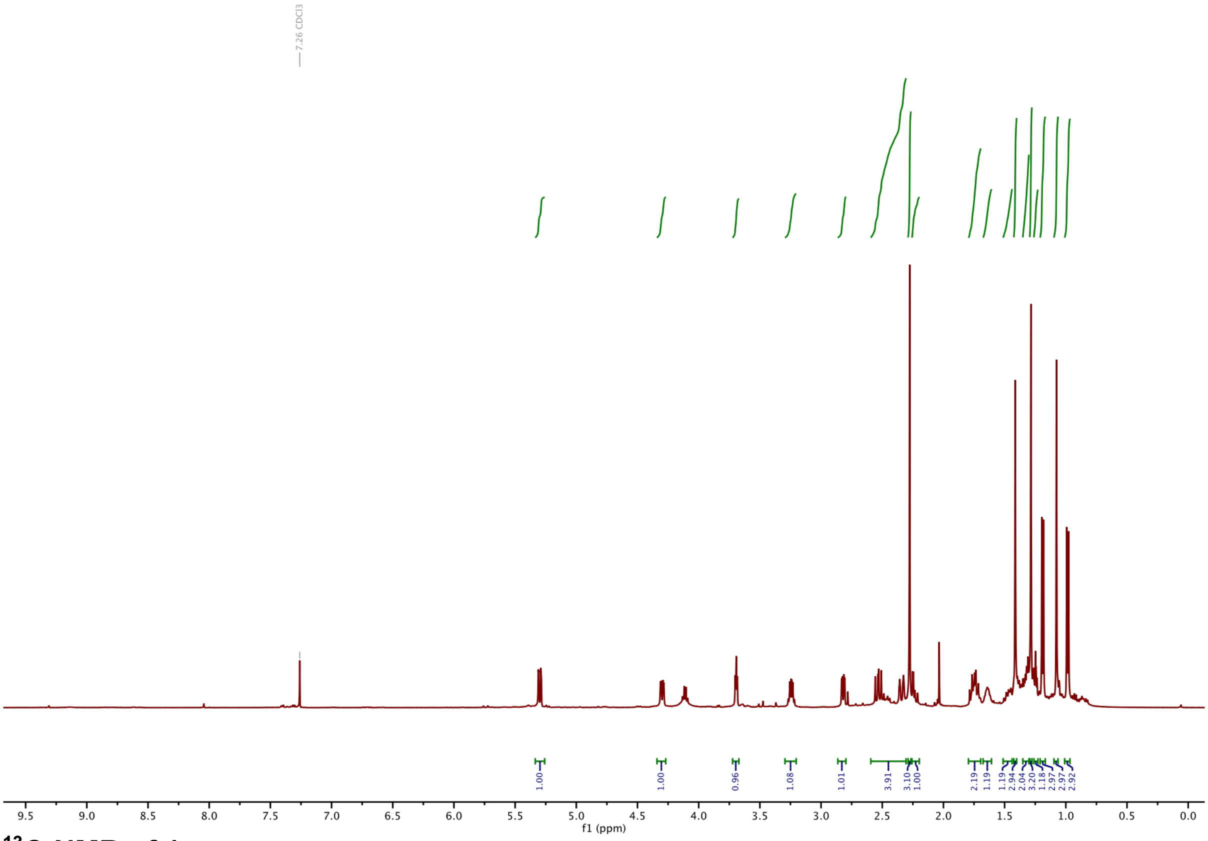

**[13]C-NMR of 1:**

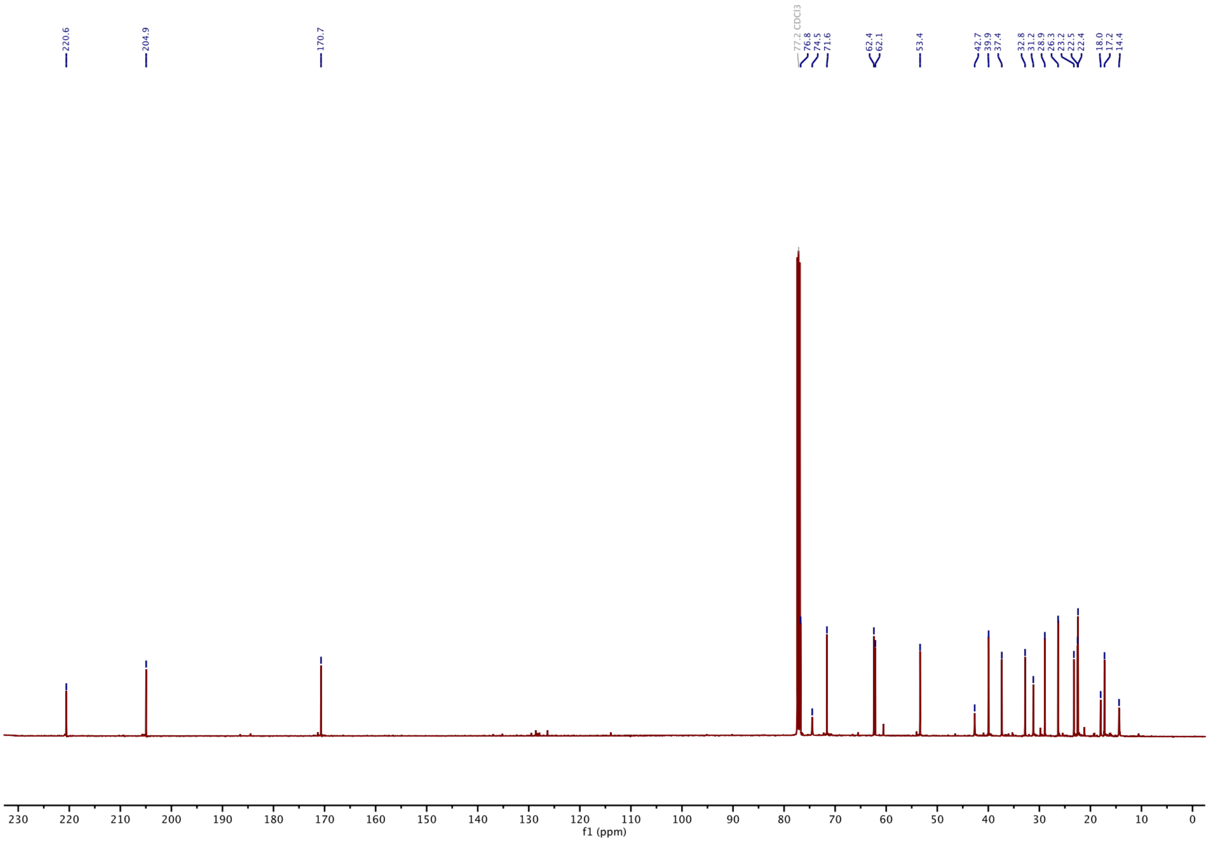

#### [1]H-NMR of 2:

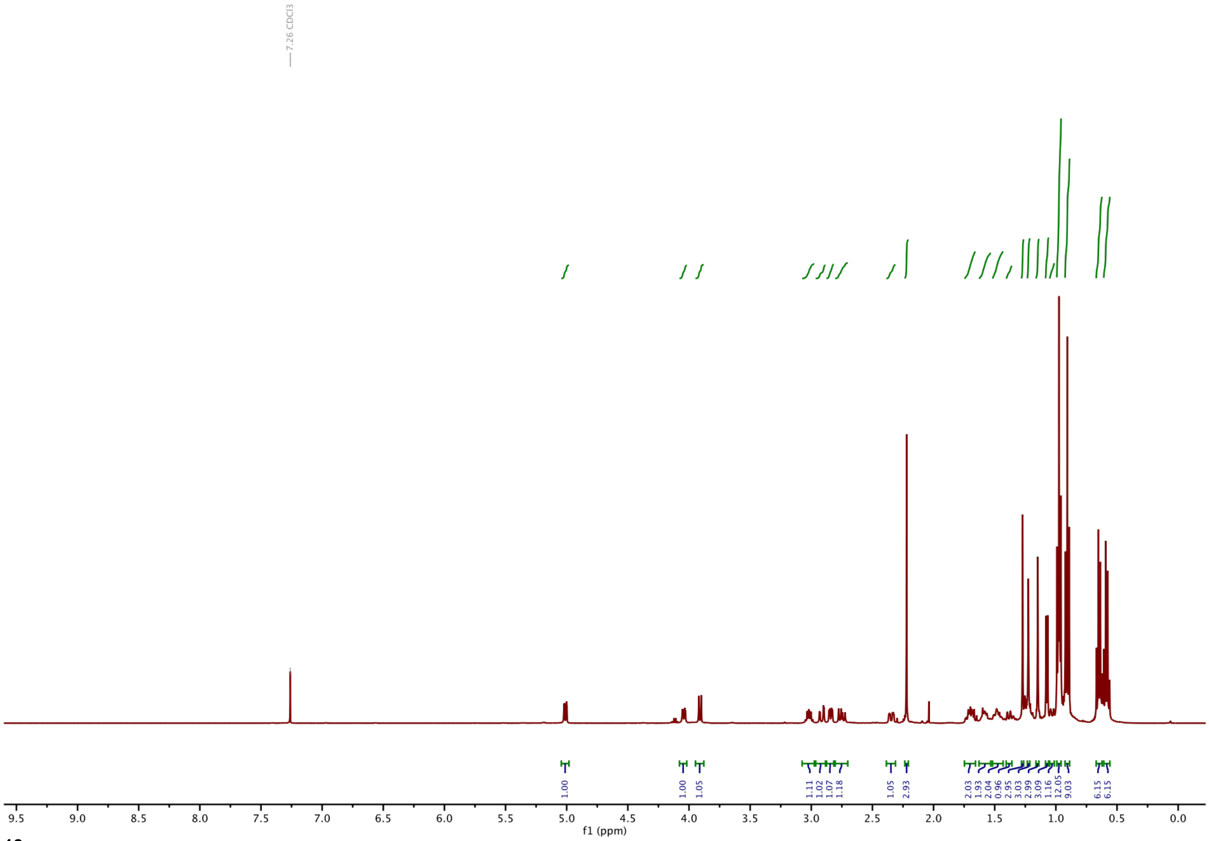

#### [13]C-NMR of 2:

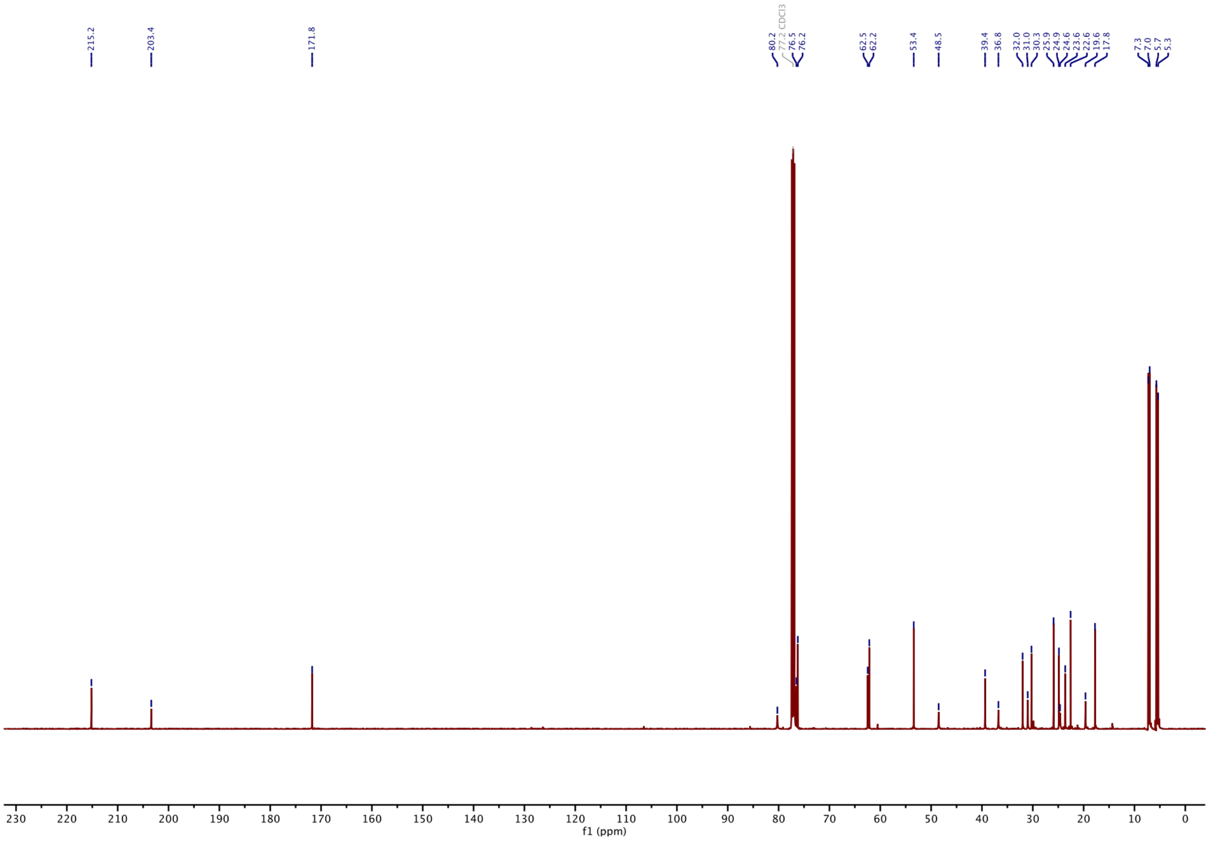

#### $^{1}$H-NMR of 3:

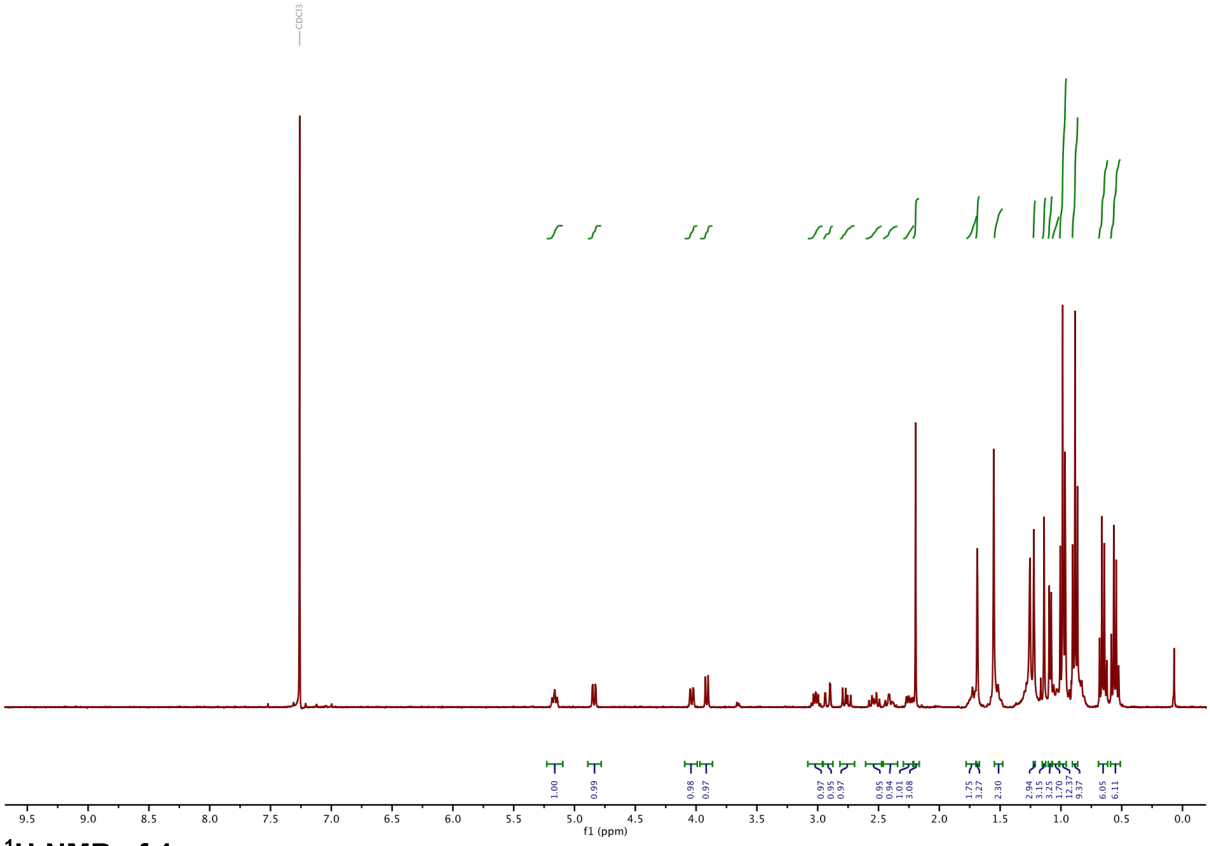

#### $^{1}$H-NMR of 4:

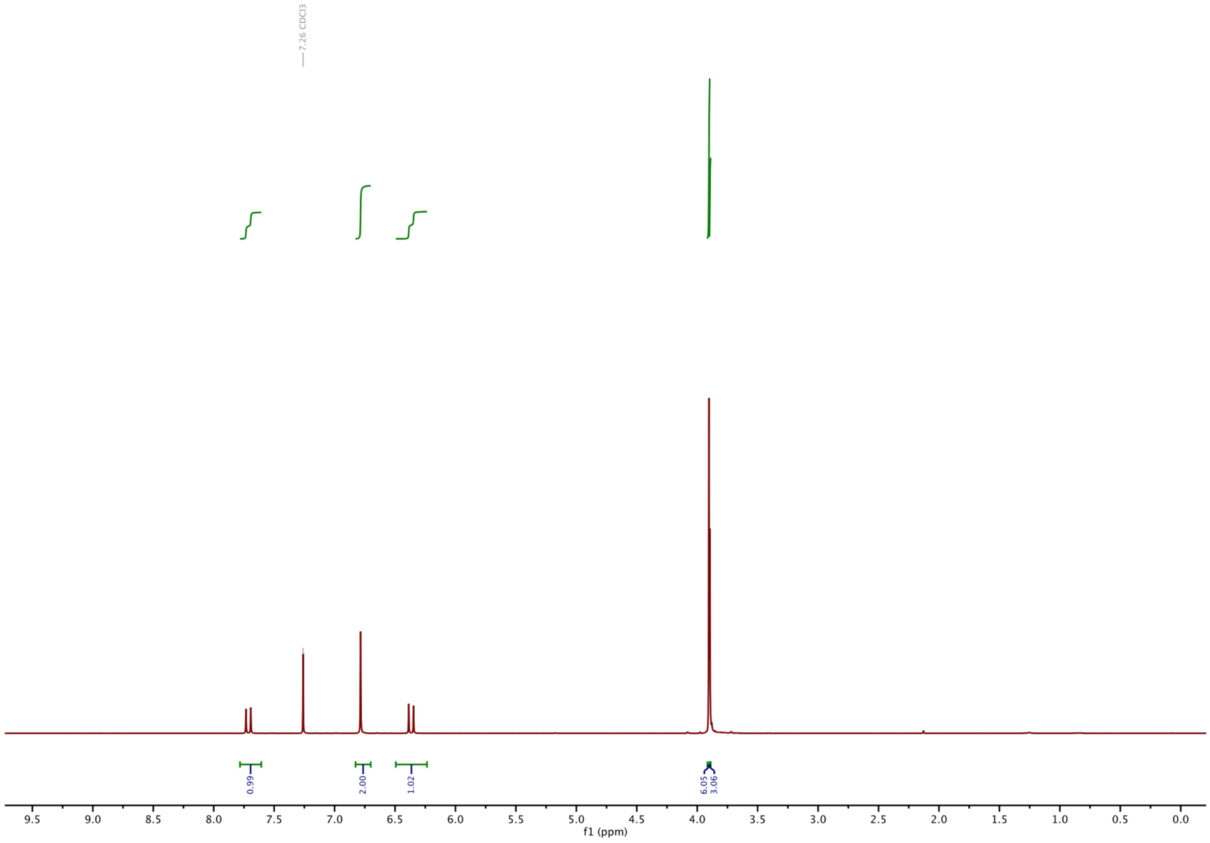

#### $^{13}$C-NMR of 4:

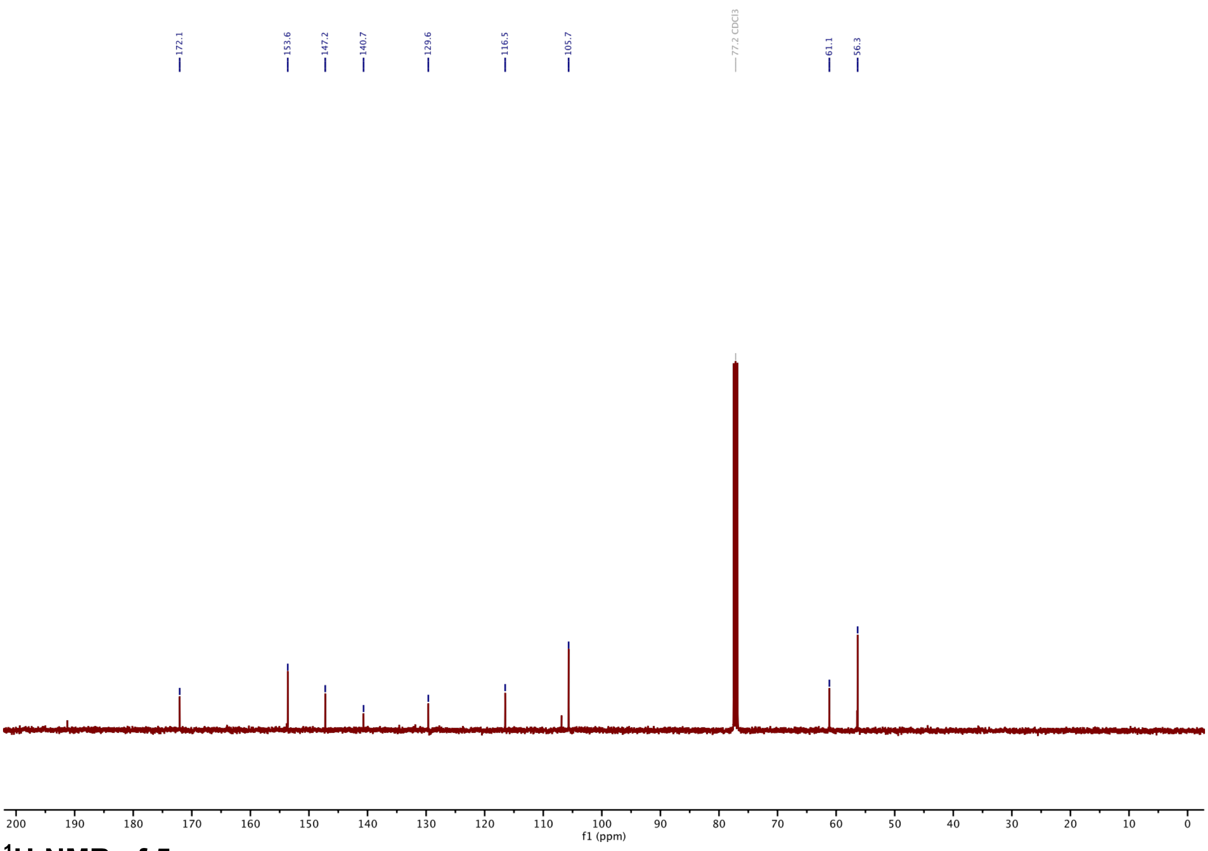

#### $^{1}$H-NMR of 5:

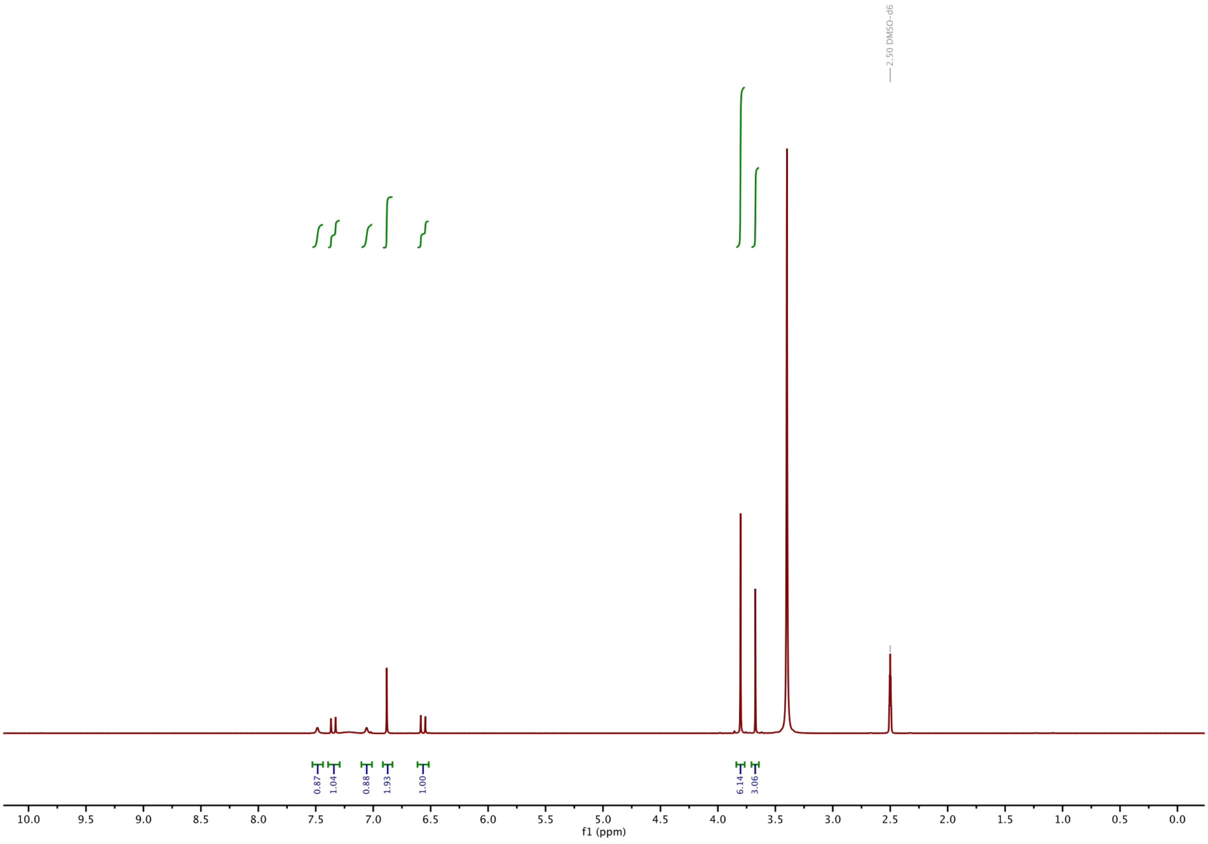

#### $^{13}$C-NMR of 5:

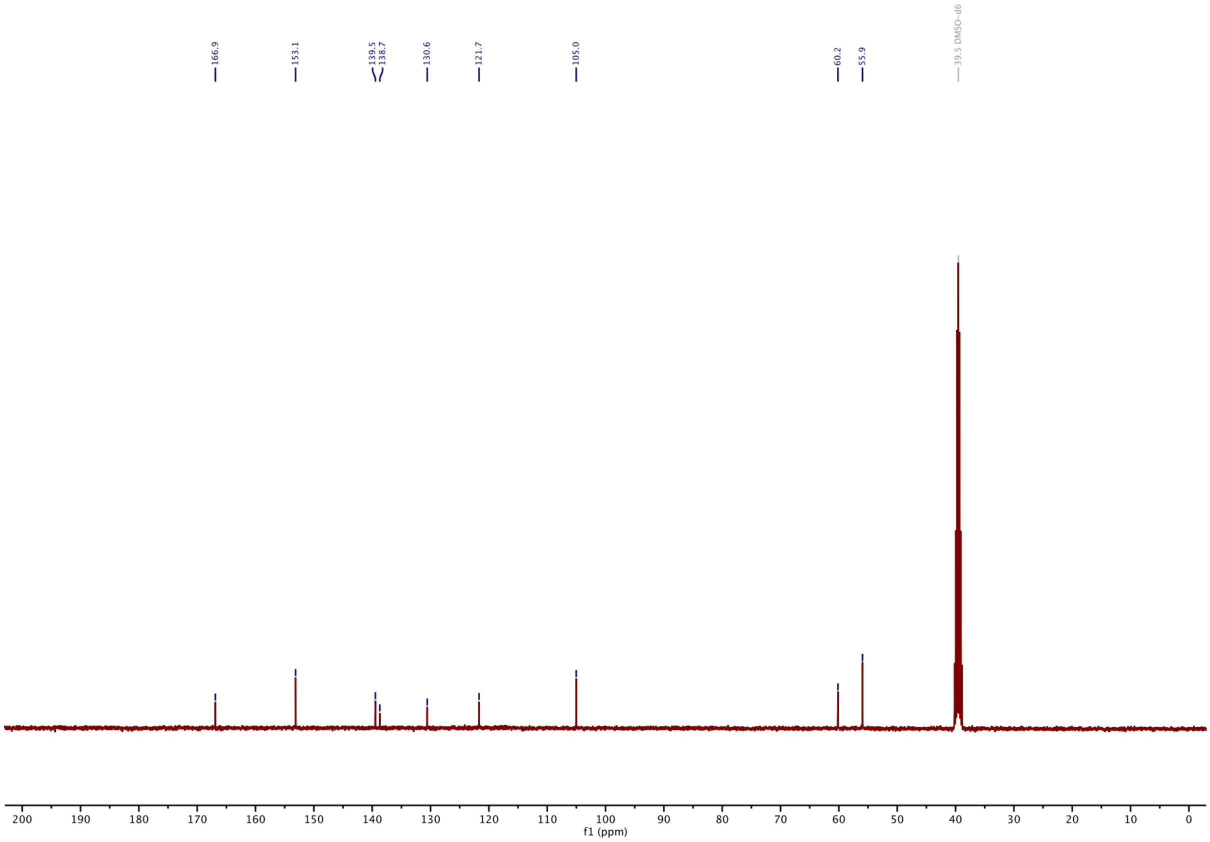

#### $^{1}$H-NMR of 6:

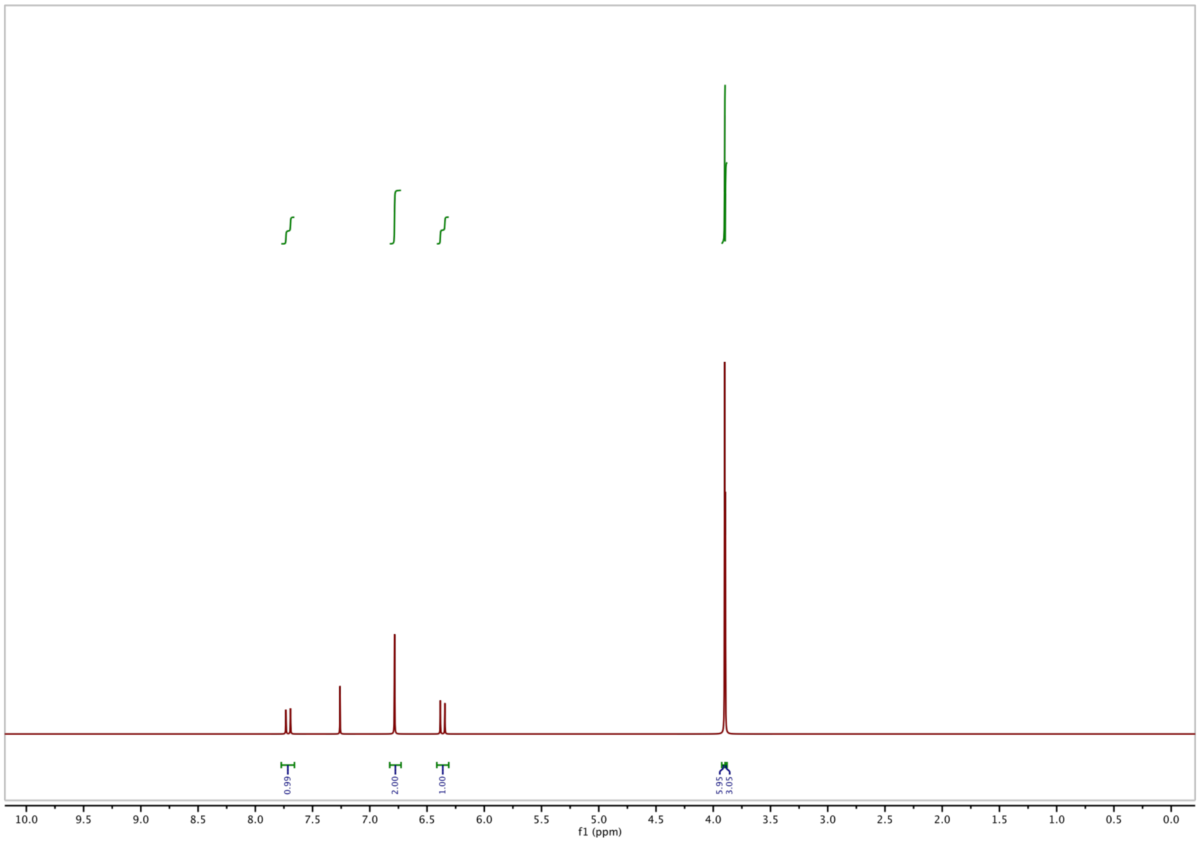

#### $^{13}$C-NMR of 6:

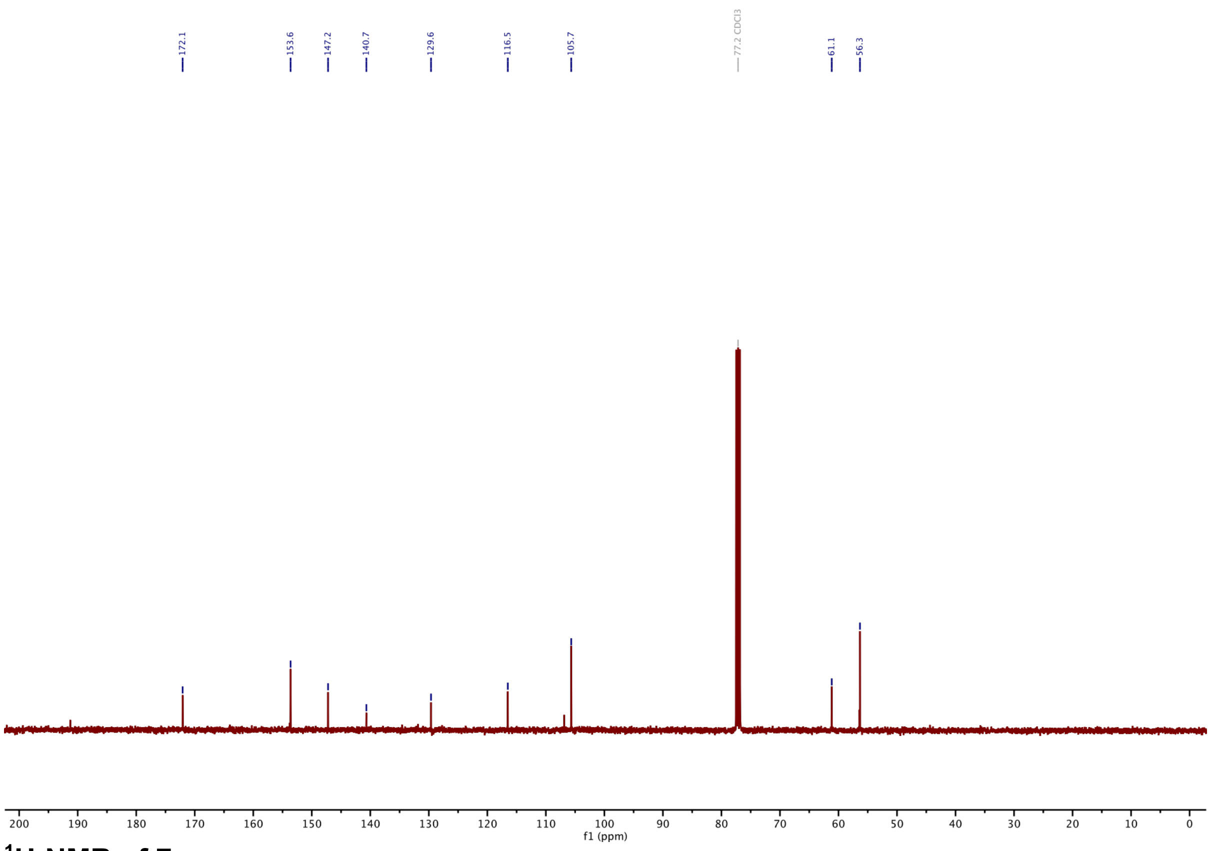

#### $^{1}$H-NMR of 7:

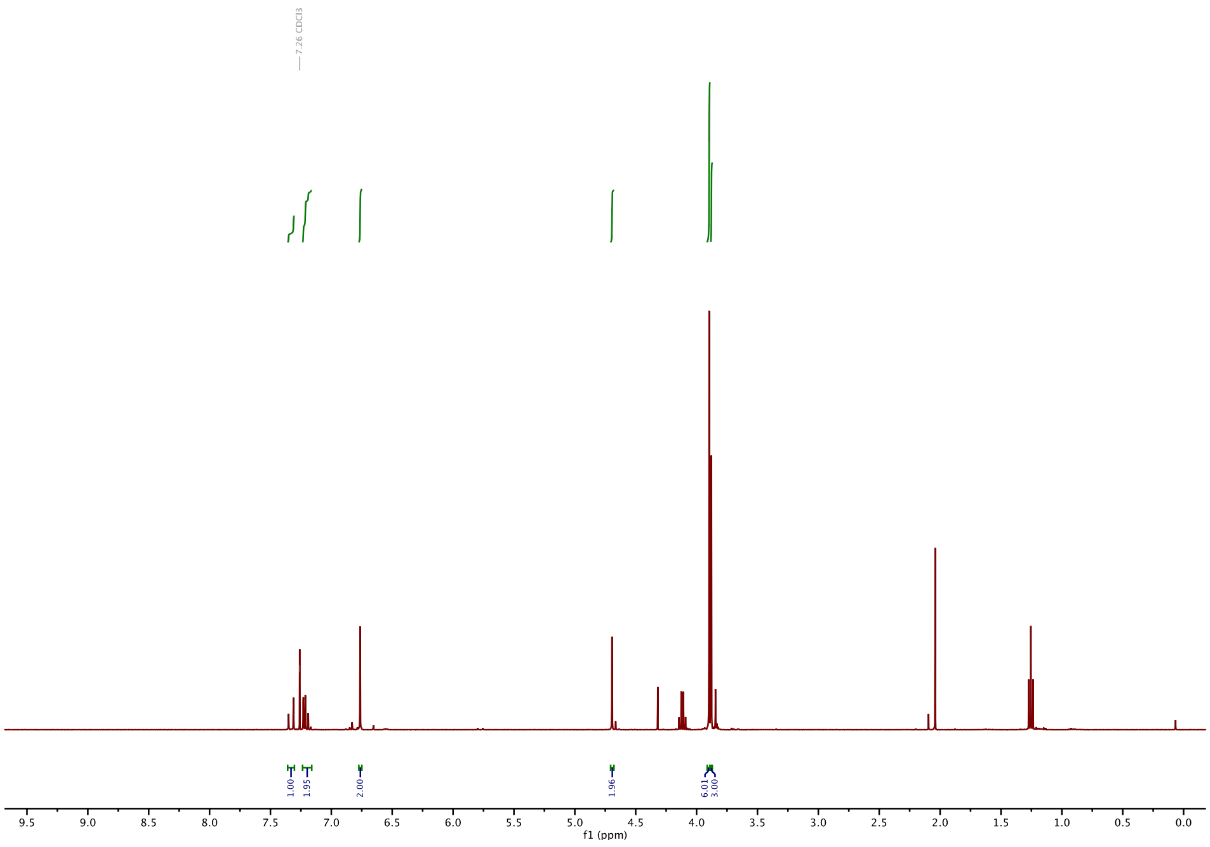

#### [13]C-NMR of 7:

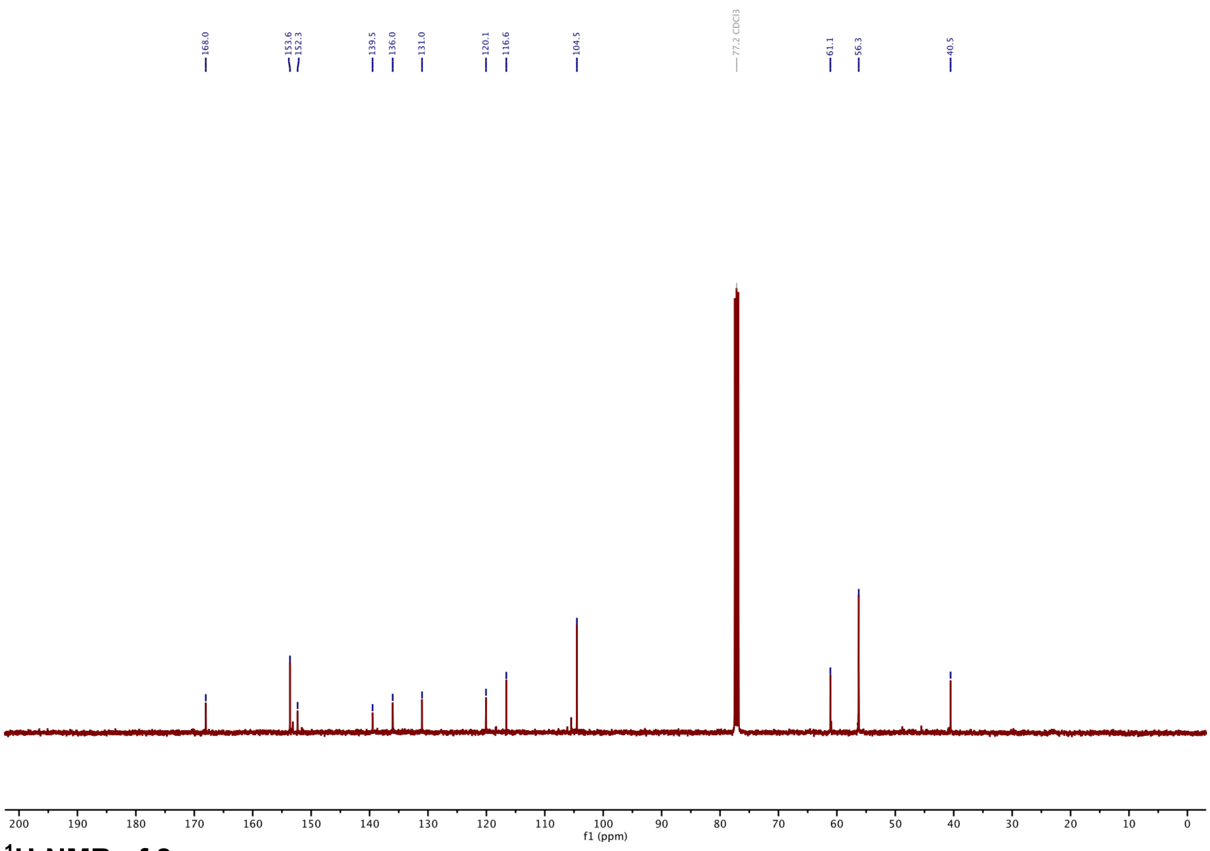

#### [1]H-NMR of 8:

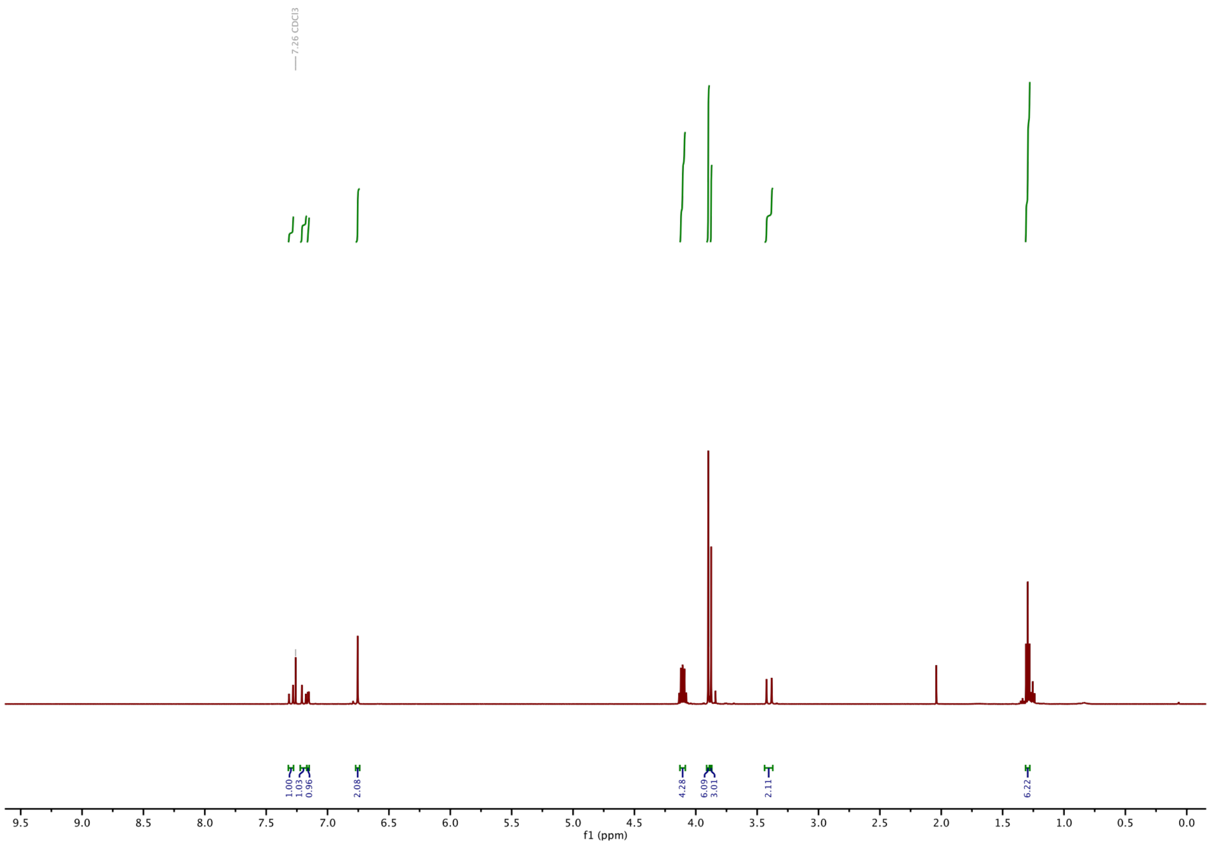

**$^{13}$C-NMR of 8:**

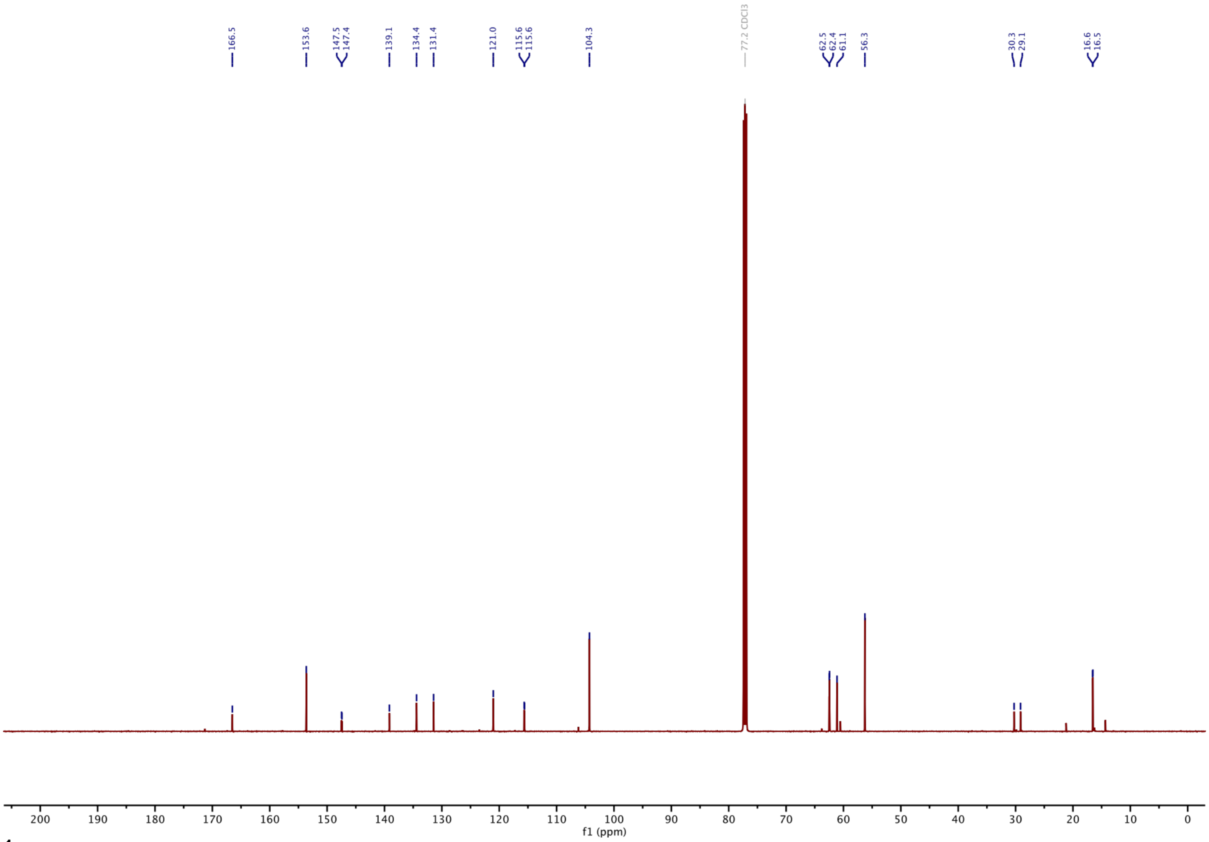

**$^{1}$H-NMR of 11a:**

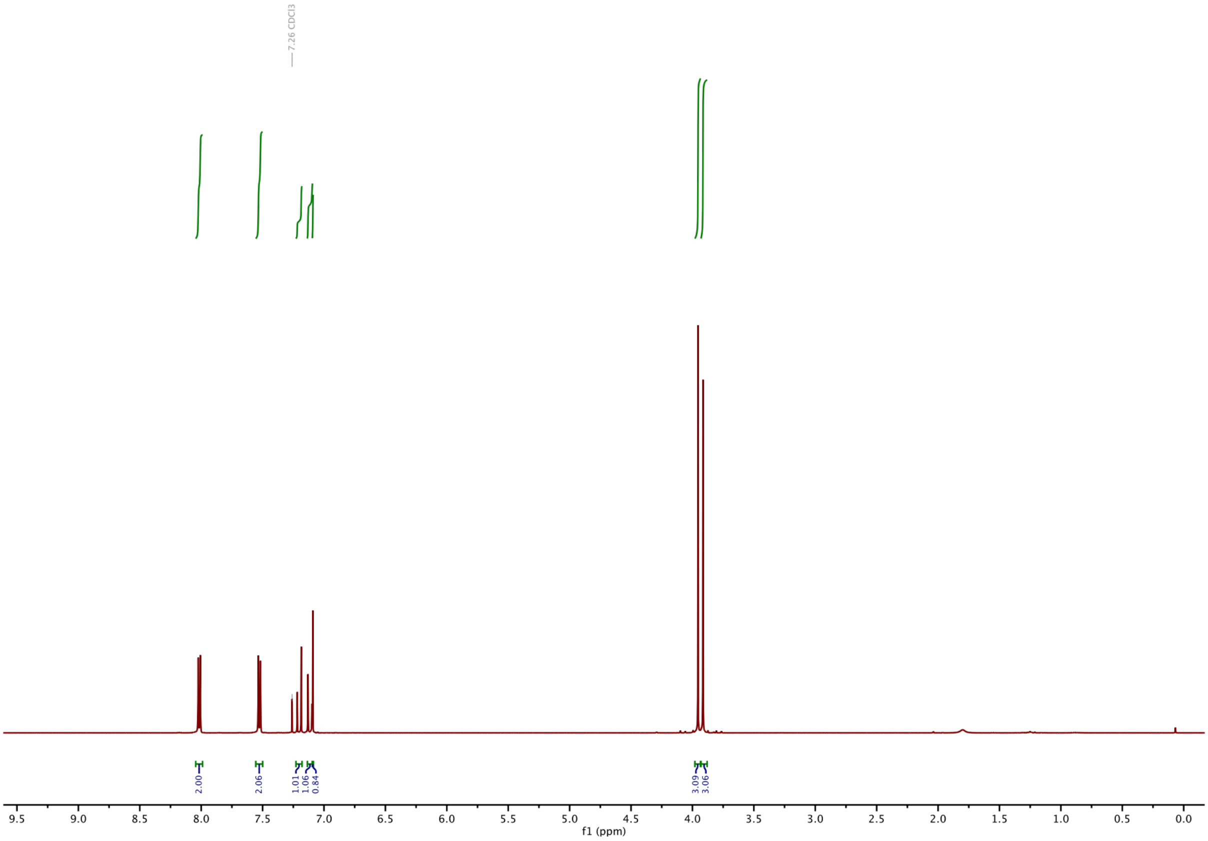

#### $^{13}$C-NMR of 11a:

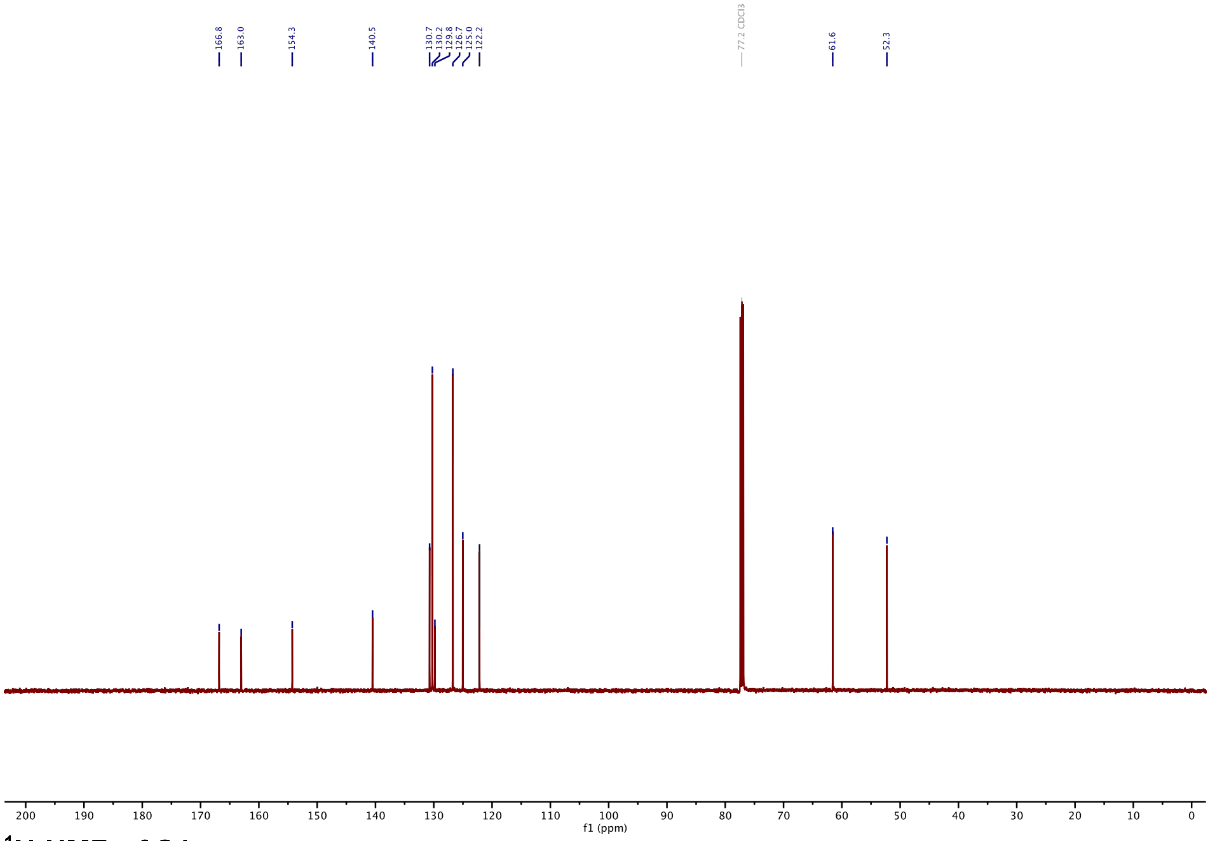

#### $^{1}$H-NMR of S1:

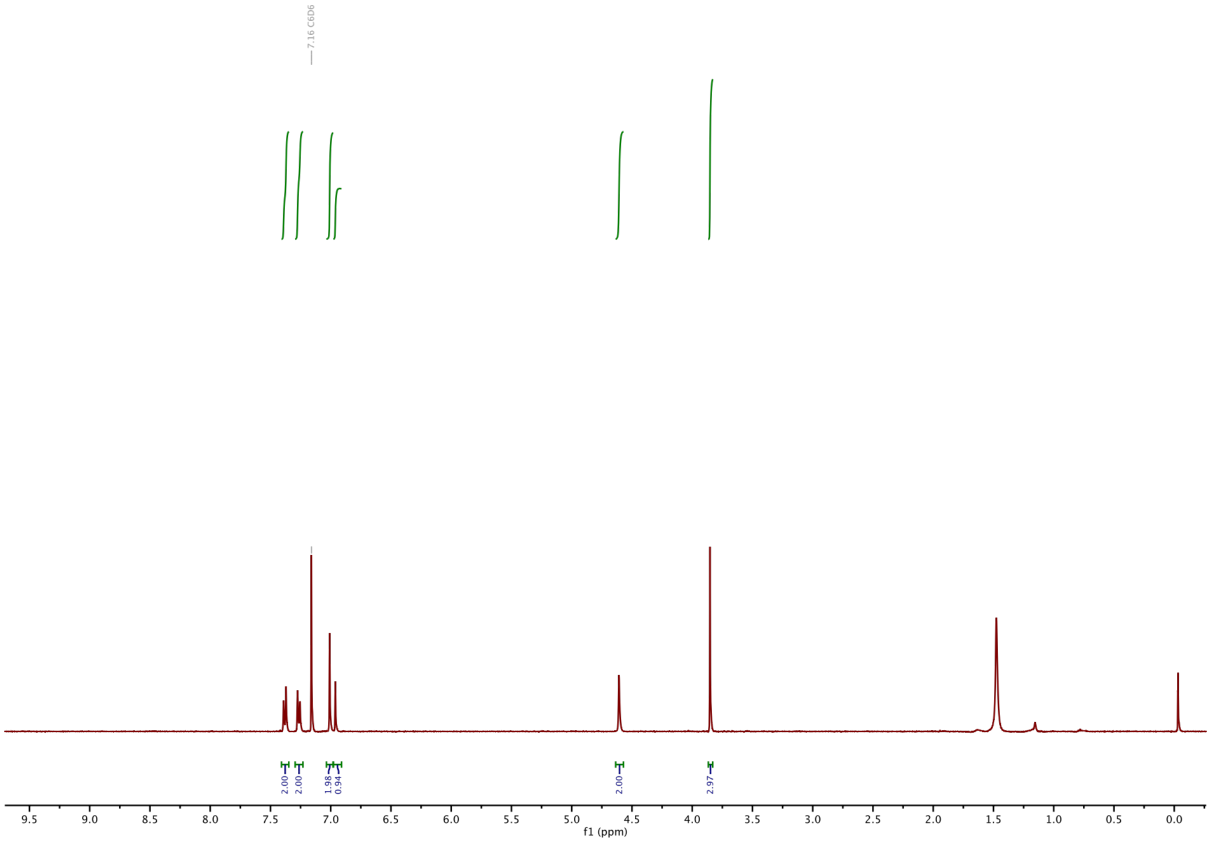

#### $^{13}$C-NMR of S1

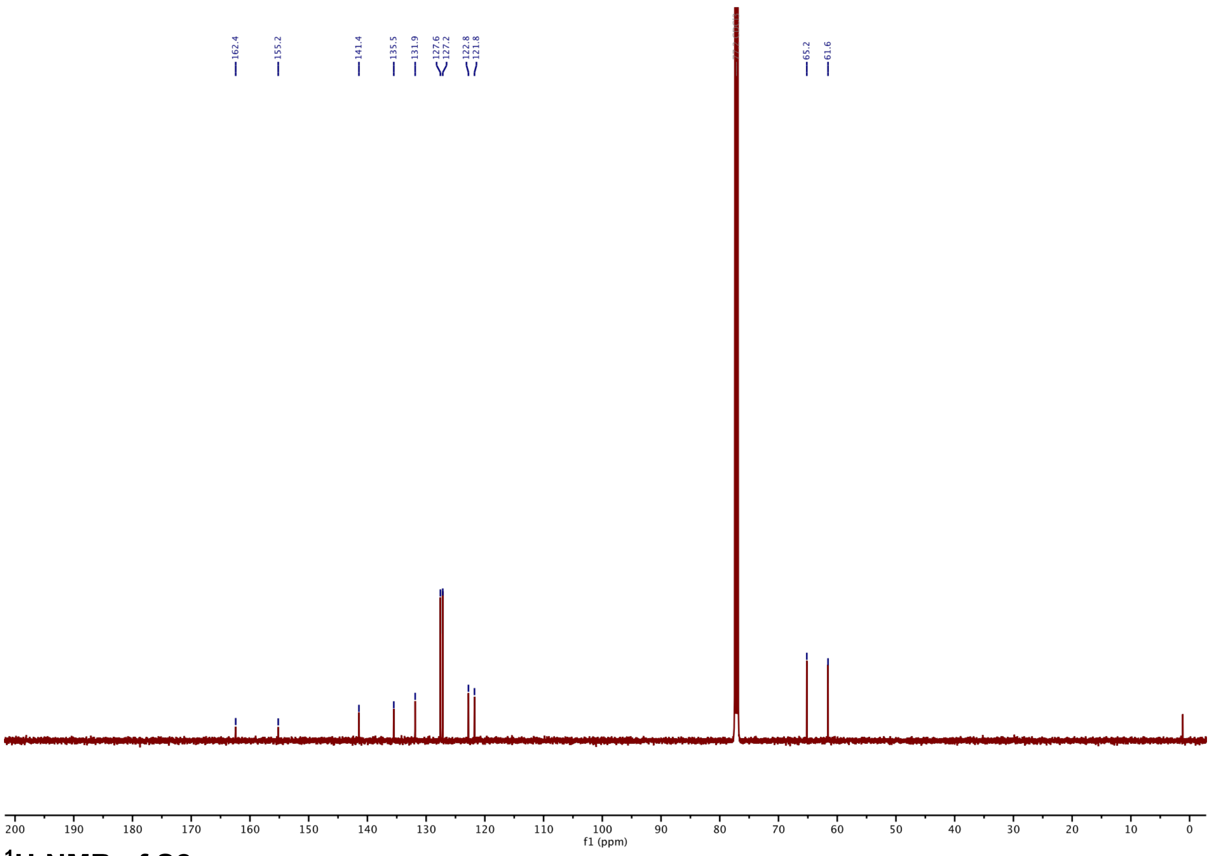

#### $^{1}$H-NMR of S2:

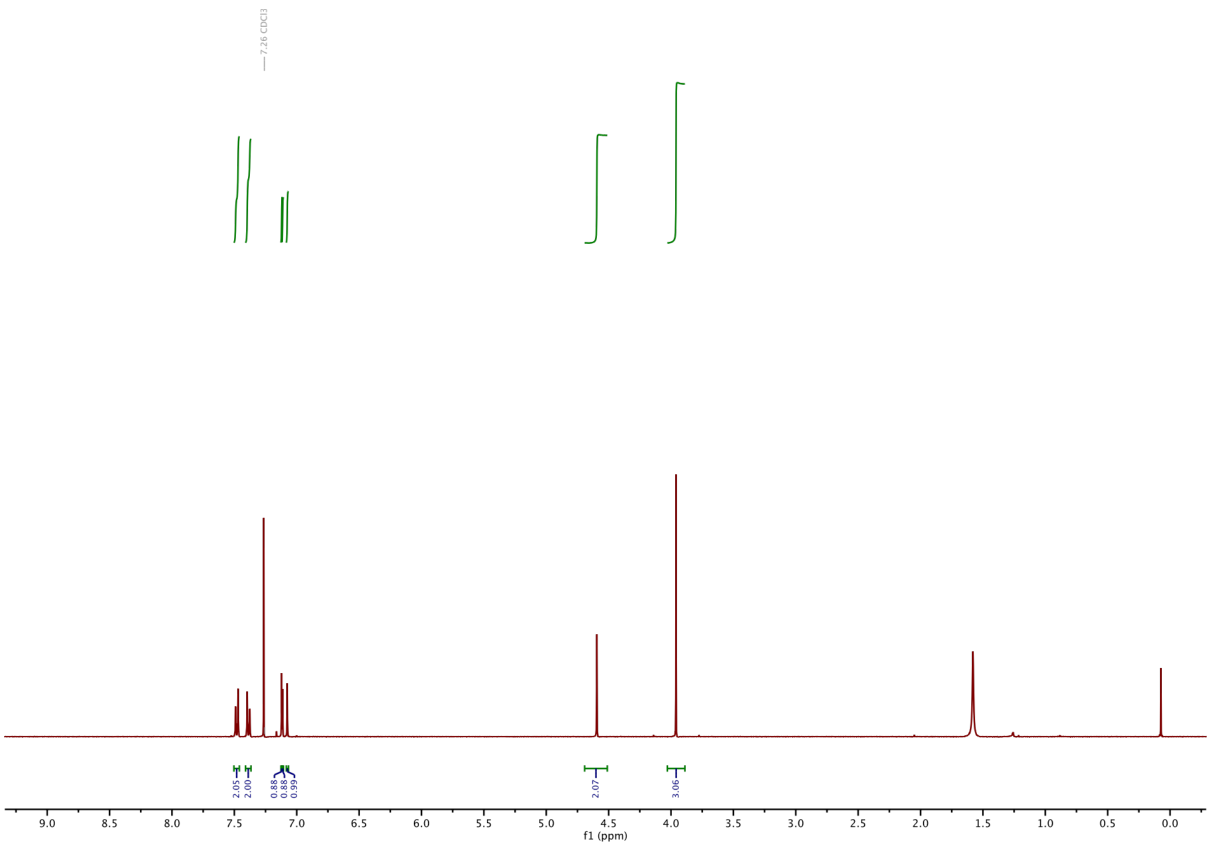

#### $^{13}$C-NMR of S2:

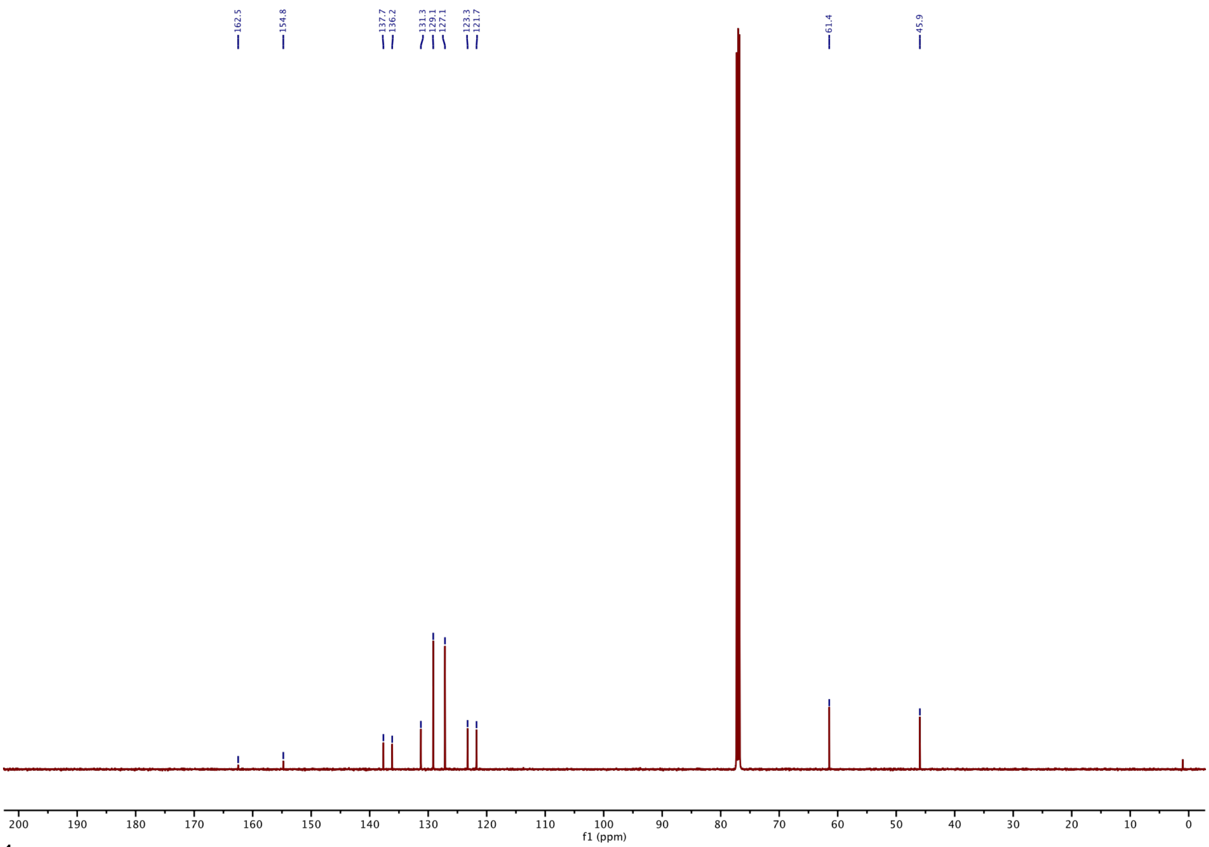

#### $^{1}$H-NMR of 12a:

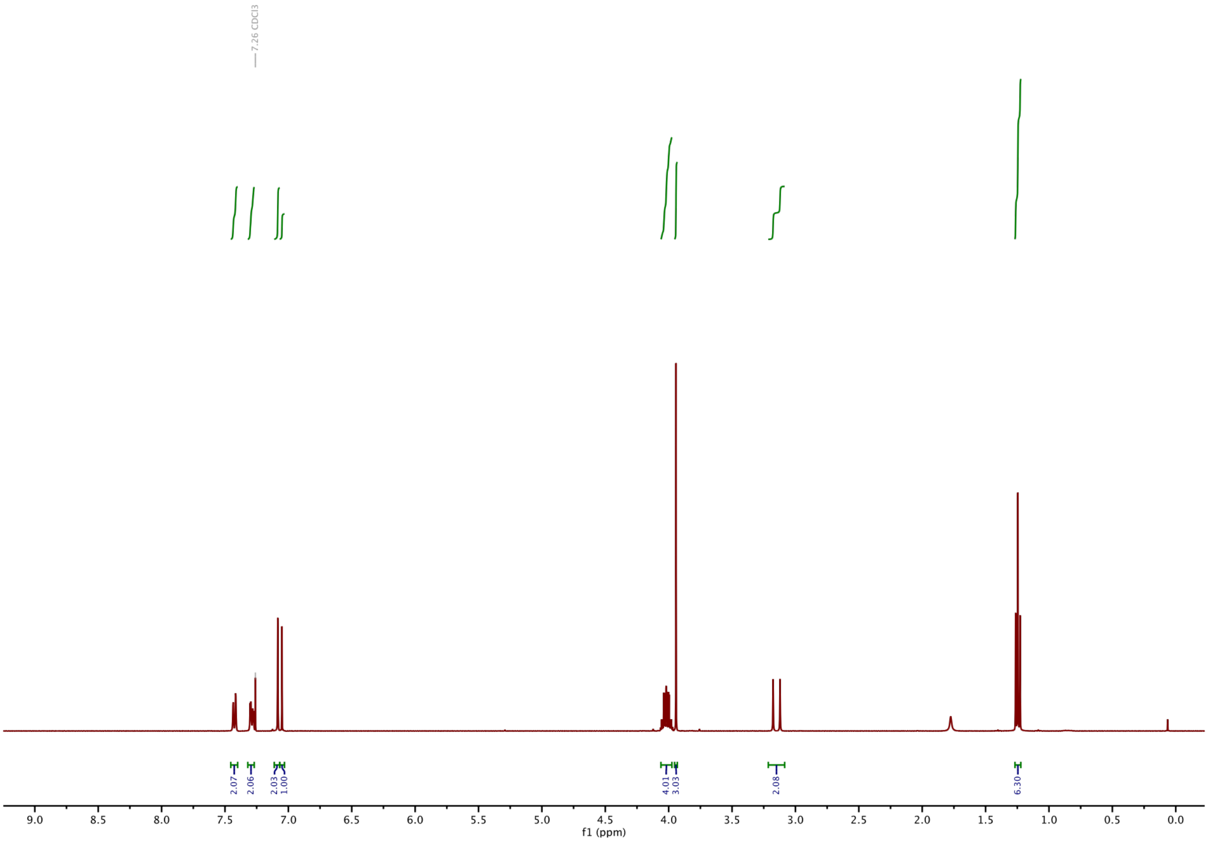

#### $^{13}$C-NMR of 12a:

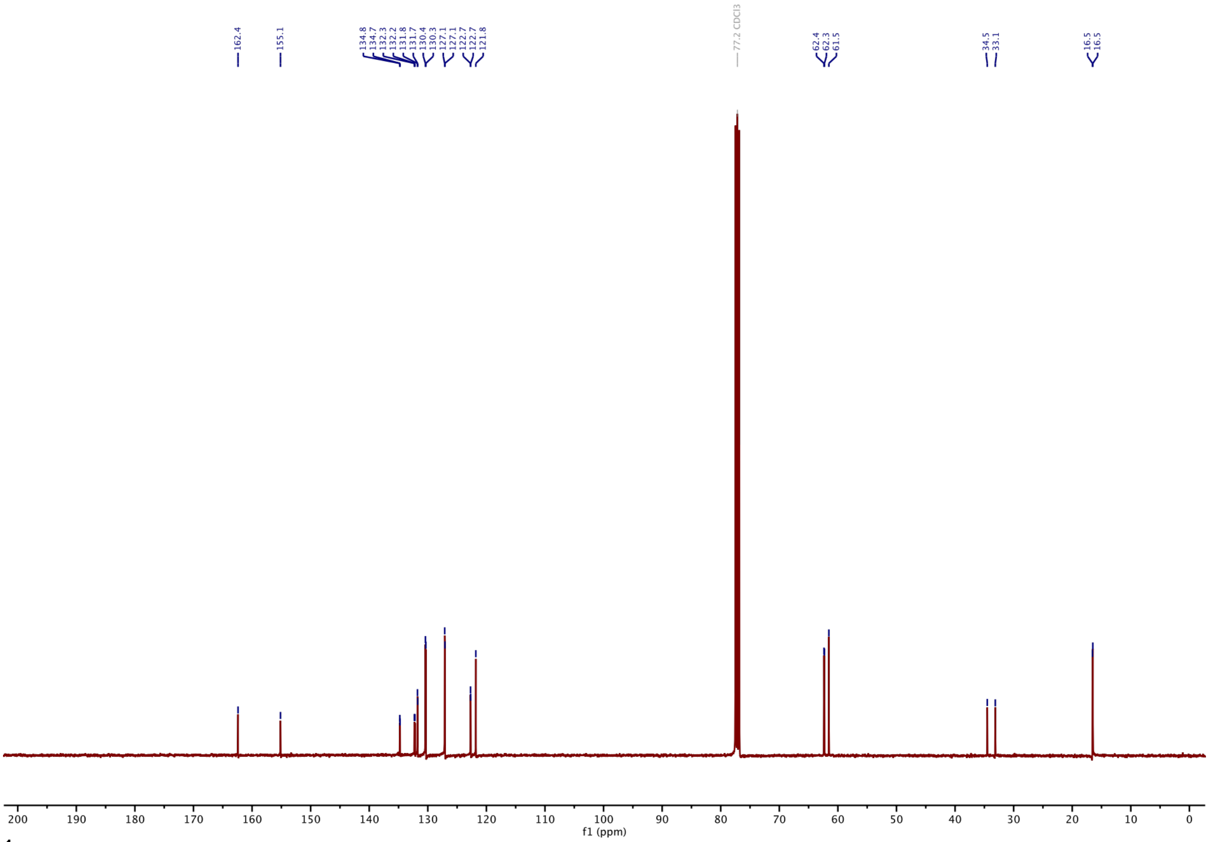

#### $^{1}$H-NMR of 11b:

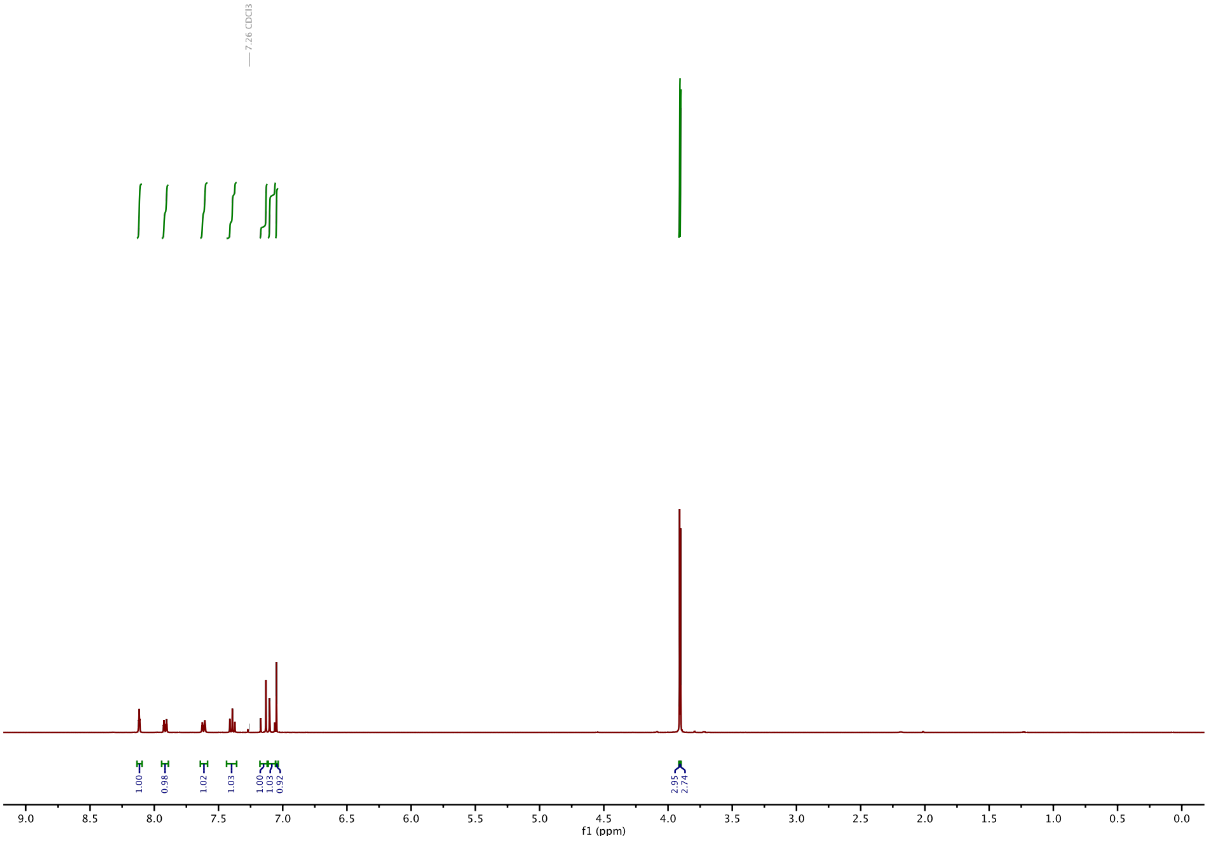

#### $^{13}$C-NMR of 11b:

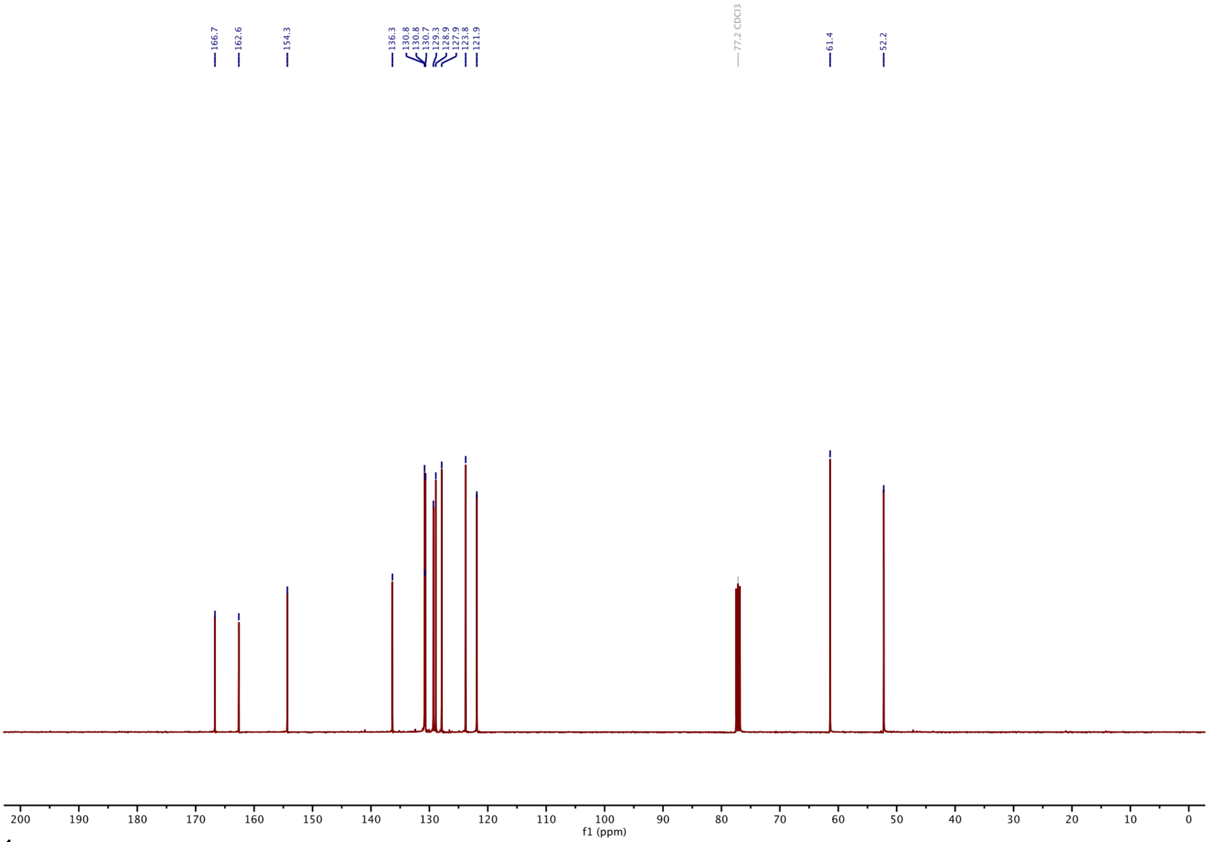

#### $^{1}$H-NMR of S3:

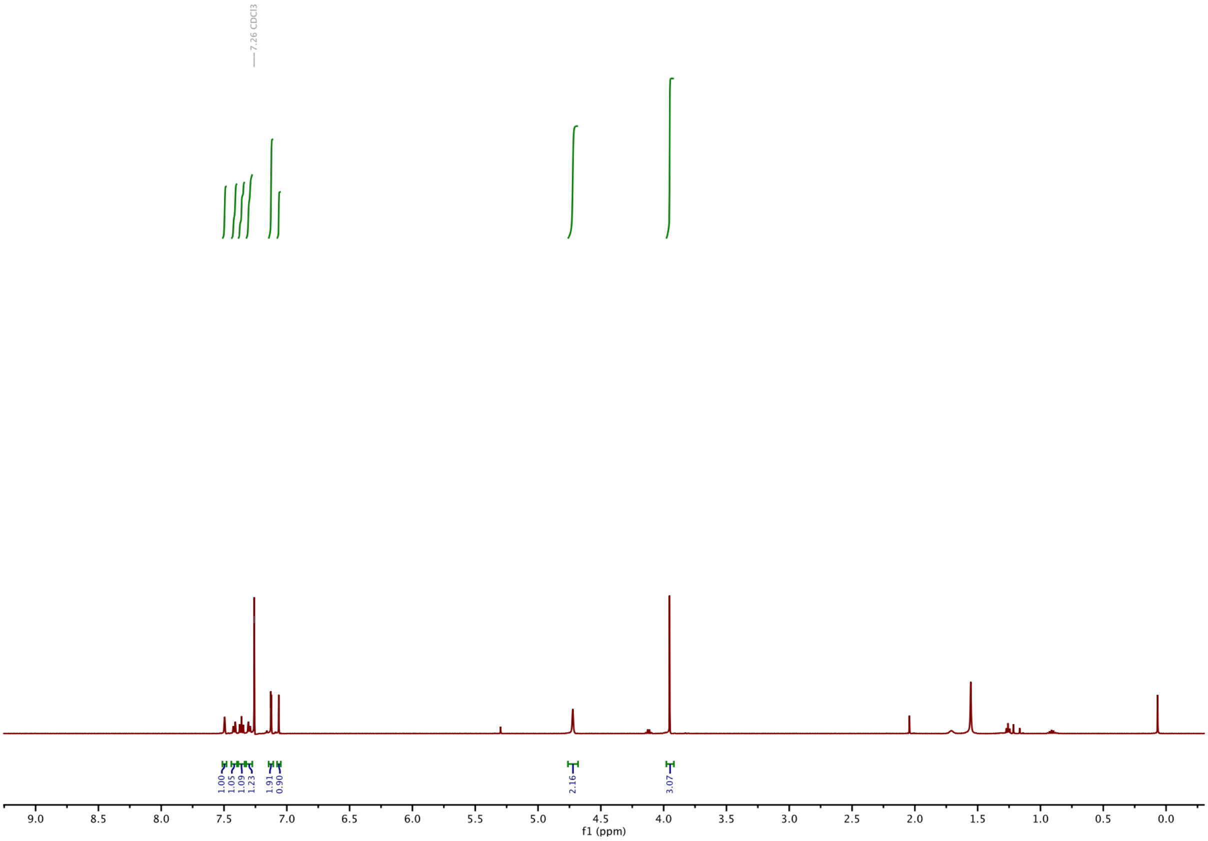

**$^{13}$C-NMR of S3:**

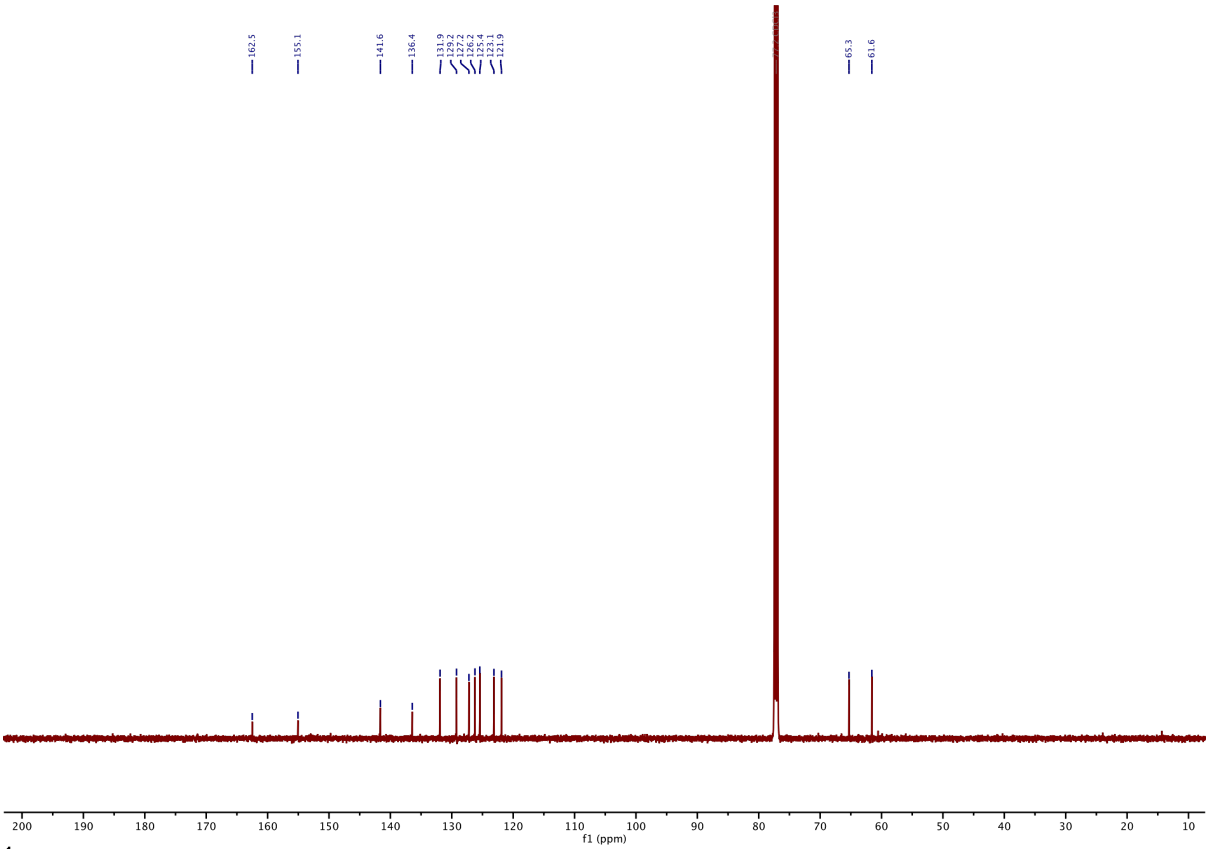

**$^{1}$H-NMR of S4:**

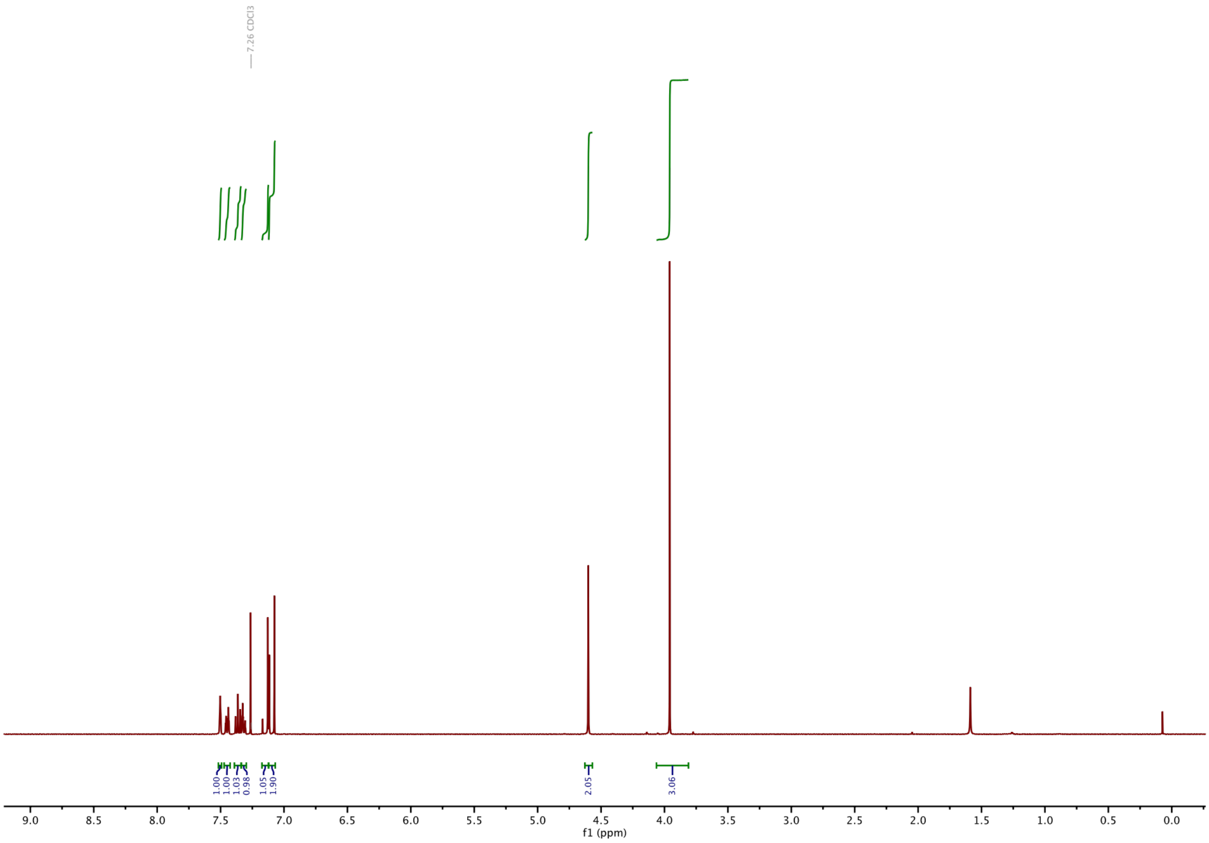

#### $^{13}$C-NMR of S4:

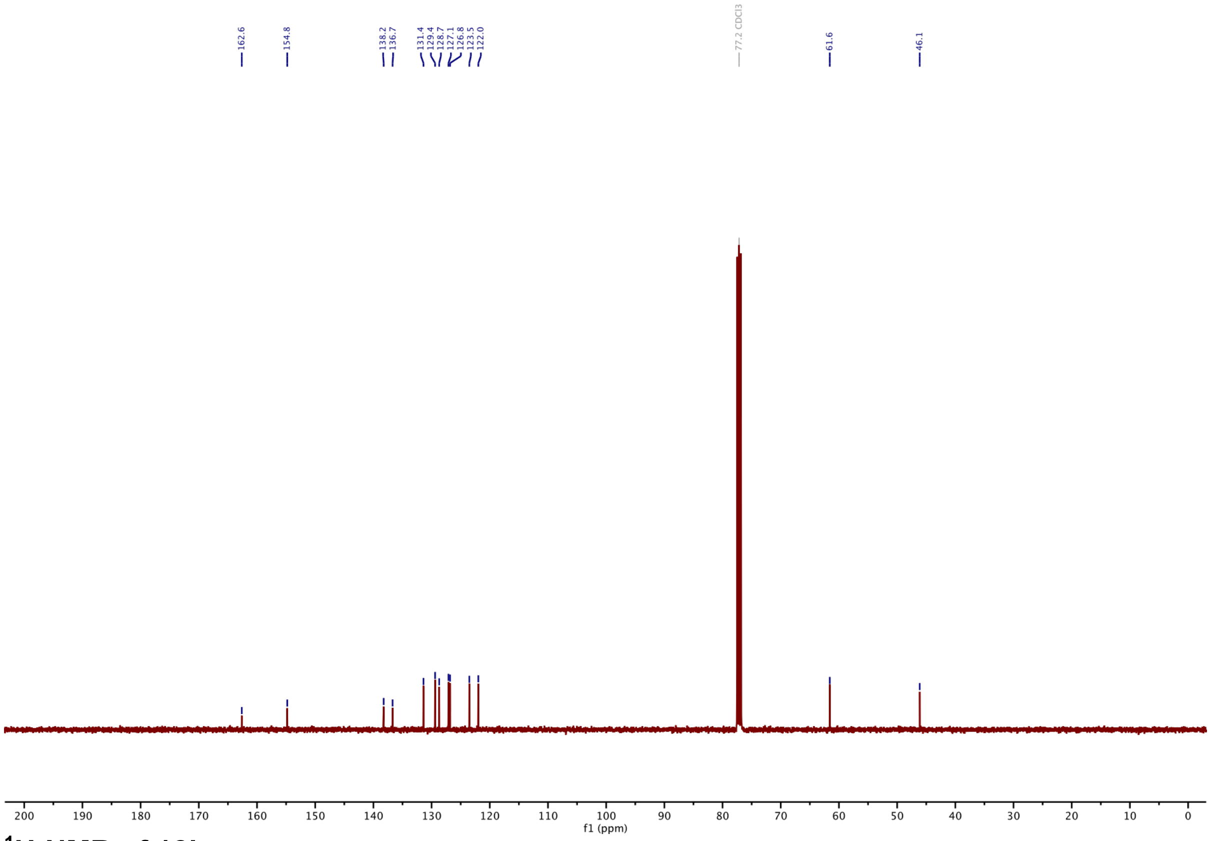

#### $^{1}$H-NMR of 12b:

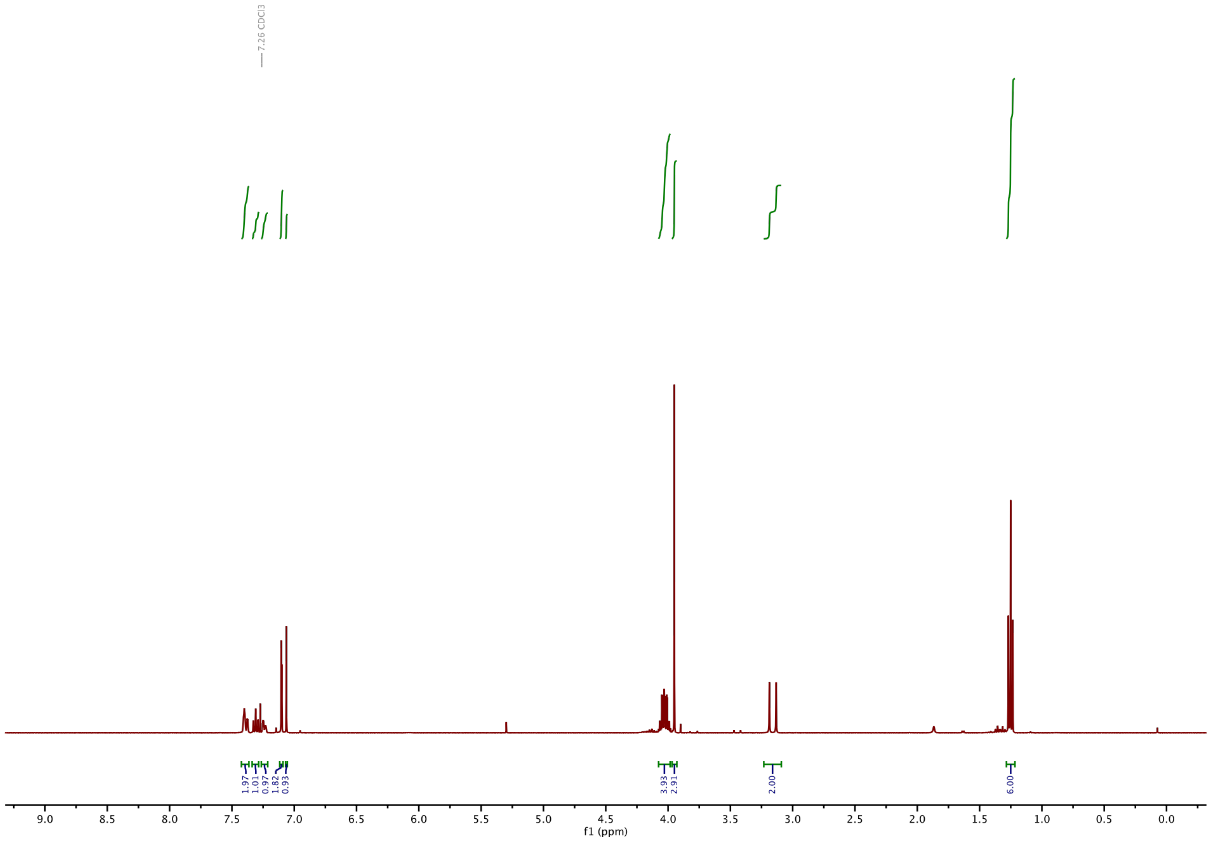

#### $^{13}$C-NMR of 12b:

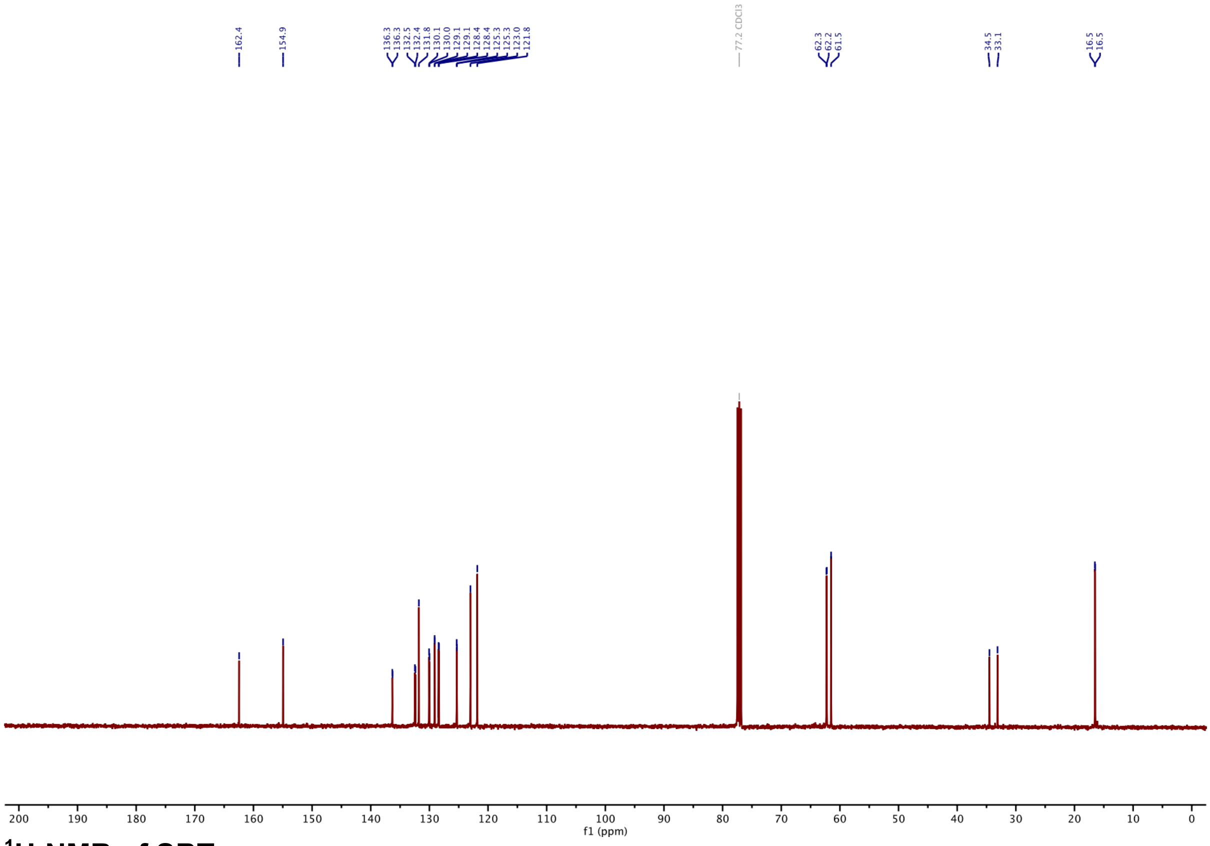

#### $^{1}$H-NMR of SBTax:

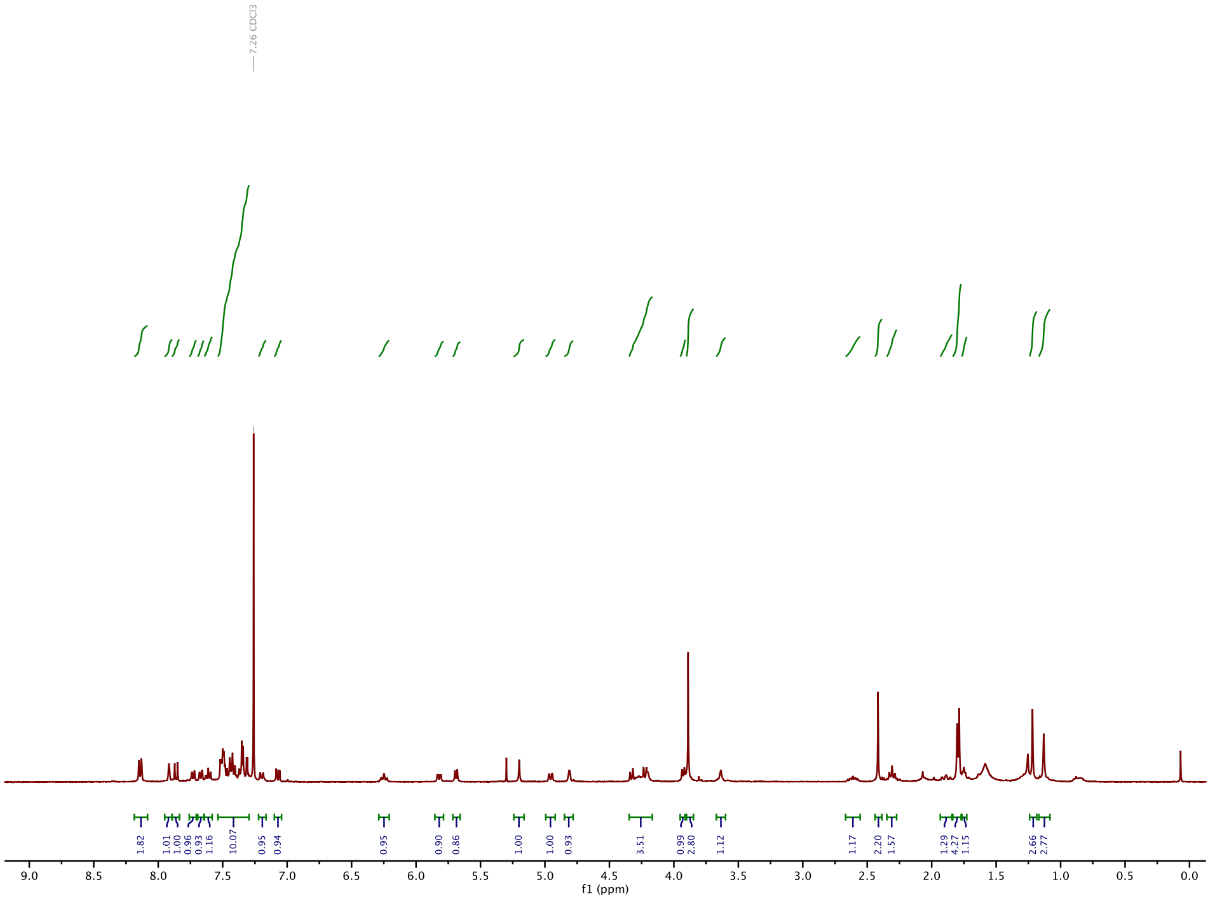

**$^{13}$C-NMR of SBTax:**

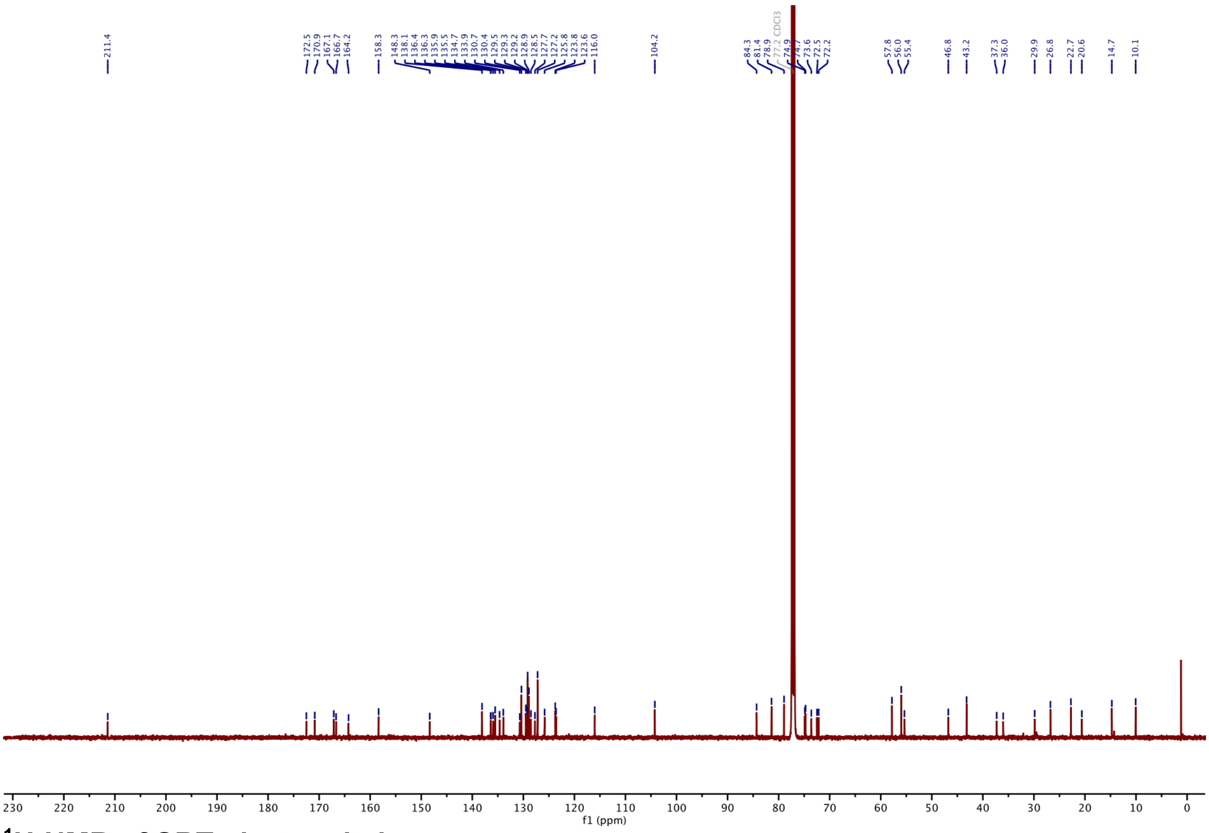

**$^{1}$H-NMR of SBT photoswitch:**

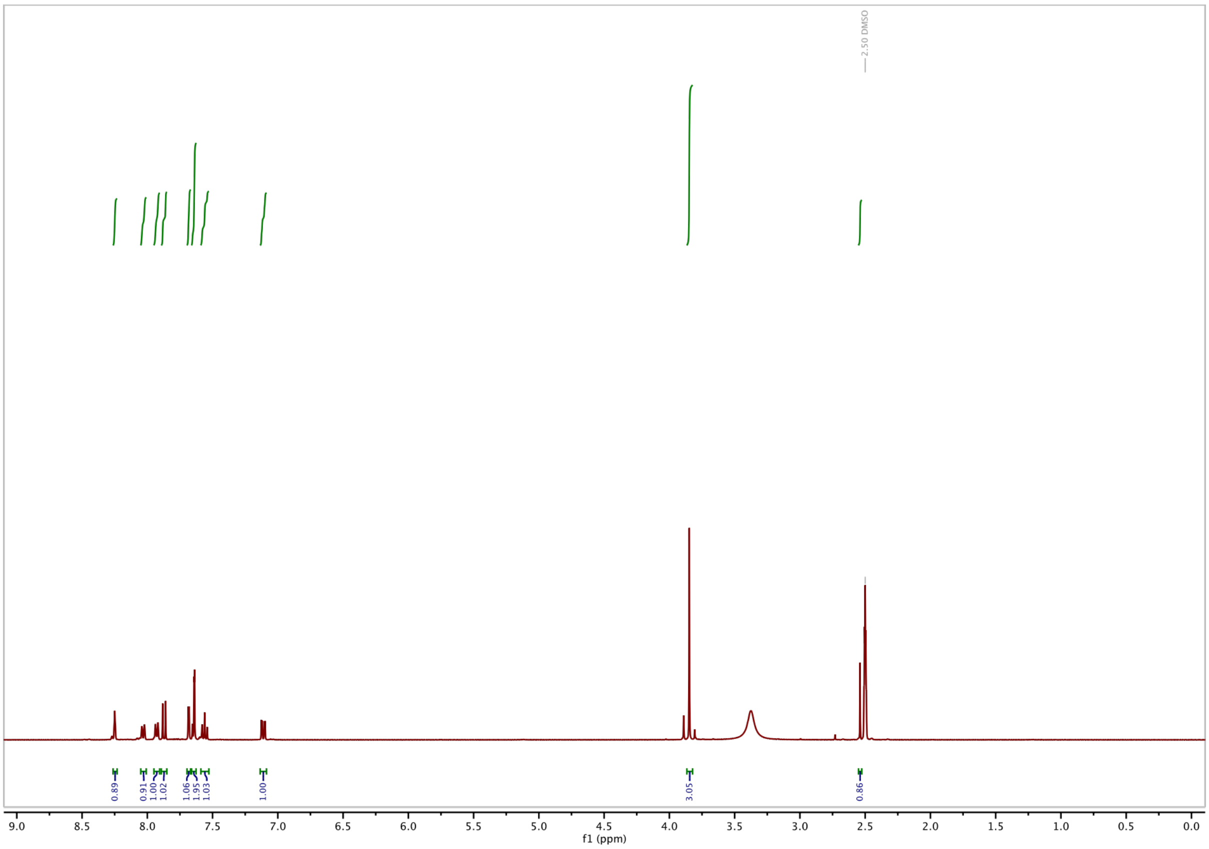

**<u>$^{13}$C-NMR of SBT photoswitch:</u>**

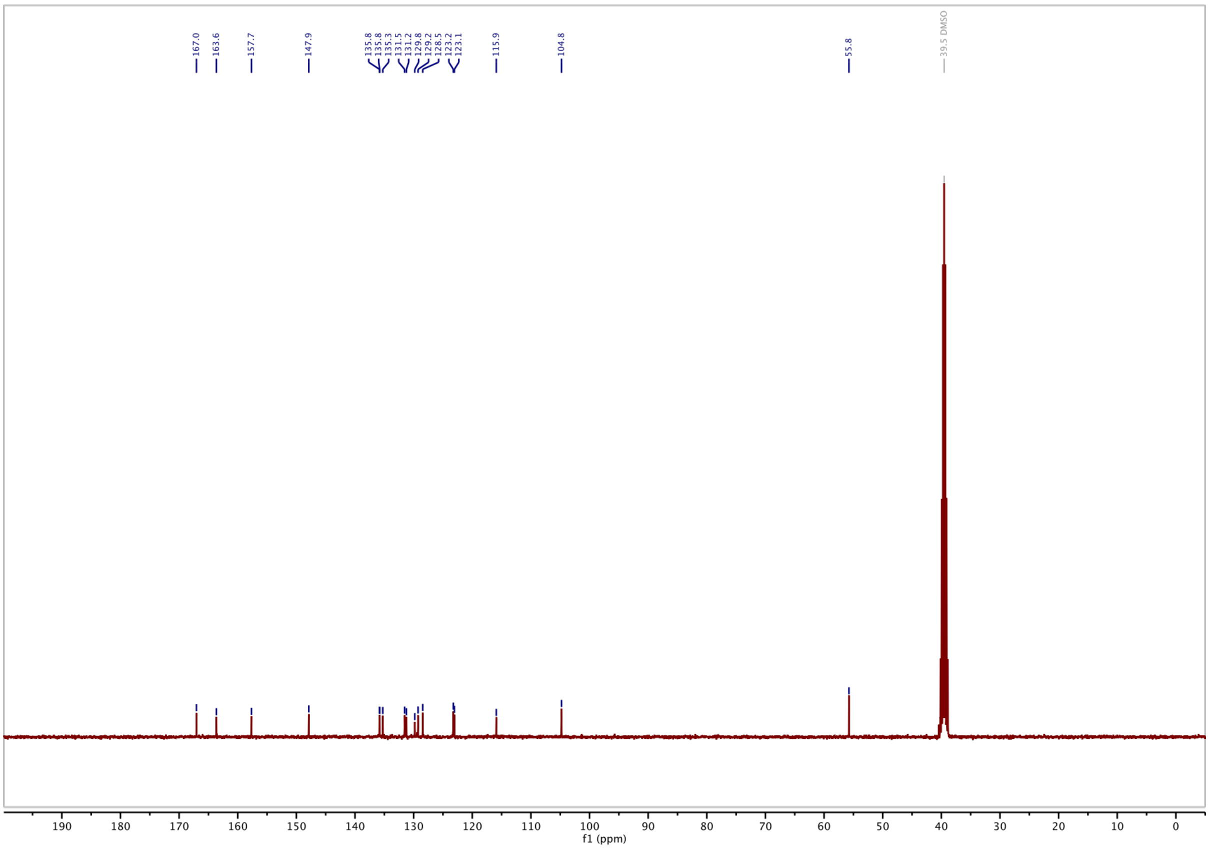

### Supporting Information Bibliography
